# Extracellular Domains of CAR Reprogram T-Cell Metabolism Without Antigen Stimulation

**DOI:** 10.1101/2023.04.03.533021

**Authors:** Aliya Lakhani, Ximin Chen, Laurence C. Chen, Mobina Khericha, Yvonne Y. Chen, Junyoung O. Park

## Abstract

Metabolism is an indispensable part of T-cell proliferation, activation, and exhaustion, yet the metabolism of chimeric antigen receptor (CAR)-T cells remains incompletely understood. CARs are comprised of extracellular domains that determine cancer specificity, often using single-chain variable fragments (scFvs), and intracellular domains that trigger signaling upon antigen binding. Here we show that CARs differing only in the scFv reprogram T-cell metabolism differently. Even in the absence of antigens, some CARs increase proliferation and nutrient uptake in T cells. Using stable isotope tracers and mass spectrometry, we observe basal metabolic fluxes through glycolysis doubling and amino acid uptake overtaking anaplerosis in CAR-T cells harboring rituximab scFv, unlike other similar anti-CD20 scFvs. Disparate rituximab and 14g2a-based anti-GD2 CAR-T cells are similarly hypermetabolic and channel excess nutrients to nitrogen overflow metabolism. Since CAR-dependent metabolic reprogramming alters cellular energetics, nutrient utilization, and proliferation, metabolic profiling should be an integral part of CAR-T cell development.

## Introduction

Chimeric antigen receptors (CARs) are synthetic proteins that activate T cells upon binding cognate antigens^1^. CAR-T cell therapies targeting CD19 or B-cell maturation antigen (BCMA) have received FDA approval for the treatment of B-cell leukemia, lymphoma, and multiple myeloma^2–4^. Despite recent successes, the design of new CARs relies on heuristics rather than quantitative metrics^5^.

Metabolism fuels T cells by supplying cellular energy and biochemical building blocks necessary for proliferation, cytokine production, and cytotoxicity. The inner workings of T-cell metabolism have become increasingly accessible due to improvements in analytical techniques such as electrochemical probes, NMR, and mass spectrometry^6–9^. Upon activation, T cells undergo metabolic reprogramming to meet the increased energetic and biosynthetic demand^10, 11^. Naïve T cells are metabolically quiescent; they have slow glycolysis and rely more on fatty acid oxidation and oxidative phosphorylation to produce ATP^10–12^. The initiation of T-cell activation is triggered by T-cell receptor (TCR) signaling and costimulation, resulting in T-cell proliferation, differentiation, and production of effector molecules like cytokines^13, 14^. The TCR activates signaling pathways that lead to increased glucose uptake^10–14^. Activated T cells use the increased glucose flux to support energy metabolism, the pentose phosphate pathway (PPP), one-carbon metabolism, and fatty acid synthesis^15–17^. Similarly, when CAR-T cells encounter cognate antigens, they mount an immune response by increasing nutrient uptake and shifting to anabolism to support proliferation, differentiation, and effector functions^11, 18–21^. However, persistent antigen stimulation exhausts T cells by impairing oxidative phosphorylation and inducing mitochondrial oxidative stress to limit nucleotide biosynthesis and proliferative capacity^22^. These metabolic alterations impact anti-tumor responses and contribute to the development of terminal dysfunction in CAR-T cells^22, 23^.

As different cancer targets require different CAR-T cells, we asked whether the choice of CAR would affect T-cell metabolism. The antigen specificity of the CAR molecule is dictated by its extracellular ligand-binding domain, which is most commonly a single-chain variable fragment (scFv). The scFv is a fusion protein that combines the variable regions of the heavy chain (V_H_) and light chain (V_L_) of an antibody. Within each variable region, the scFv is comprised of three complementarity determining regions (CDRs) interspersed among four framework regions (FRs). CARs that are different only in their scFv domain but otherwise identical have been to target diverse antigens and tumor types^5^. Previous studies have shown that the binding affinities of scFvs can affect the selectivity of CAR molecules by influencing the CAR-T cells’ ability to distinguish target cells with different antigen expression levels^24, 25^. However, if and how the choice of scFv in CAR protein design affects CAR-T cell metabolism remain unknown.

To answer this question, we characterized the basal metabolism of a panel of human T cells expressing each of seven (one anti-CD19, five anti-CD20, and one anti-GD2) CARs with identical transmembrane, costimulatory, and signaling domains using liquid chromatography-mass spectrometry (LC-MS). Even in the absence of antigens, the expression of CARs altered the proliferation and nutrient requirements of CAR-T cells. The nature of metabolic reprogramming was CAR-dependent. Using ^13^C and ^15^N tracers and quantitative flux modeling, we observed metabolic fluxes through glycolysis doubling and amino acid uptake increasing to overtake anaplerosis in CAR-T cells harboring a rituximab anti-CD20 scFv compared to EGFRt control T cells. In contrast, CAR-T cells harboring anti-CD19 scFv and other anti-CD20 scFvs with high sequence identity to the rituximab scFv upregulated metabolic activities to a lesser extent. Interestingly, 14g2a anti-GD2 and rituximab anti-CD20 CAR-T cell variants with divergent scFv sequences exhibited similarly exorbitant nutrient requirements and metabolic fluxes. These hypermetabolic CAR-T cells used the excess amino acid intake to operate nitrogen overflow metabolism, secreting nucleobases, non-essential amino acids, and ammonia. The impact of the extracellular environment on T-cell proliferation depended on scFv-specific CAR-T cell metabolism. Ammonia addition decreased the proliferation rate of rituximab anti-CD20 CAR-T cells while alanine addition increased the proliferation of anti-CD19 and non-rituximab-based anti-CD20 CAR-T cells. Since metabolism is fundamental to cellular function and proliferation and CAR domains alter T-cell metabolism, we propose the broad adoption of metabolic profiling and engineering of CAR-T cells as a critical step in the development of new CAR constructs for adoptive immunotherapy.

## Results

### CARs increase T-cell proliferation and reshape the T-cell metabolome

A key step in CAR-T cell therapy development is to eliminate inefficacious CAR candidates before performing time-consuming *in vivo* evaluation. Here, we sought to identify metabolic phenotypes that could serve as telltale signs of CAR-T cell efficacy.

We generated a panel that consists of one (FMC63-based) anti-CD19 CAR and five anti-CD20 CARs, as well as an EGFRt control (**Fig. 1a**). The anti-CD20 CARs were constructed with scFv domains derived from the antibodies Leu16 and rituximab^26–28^ or with a hybrid “RFR-LCDR” scFv that comprises the FRs of rituximab flanking the CDRs of Leu16 (**Fig. 1b)**. Furthermore, we generated rituximab.AA and RFR-LCDR.AA CARs, which contain insertions of two alanine residues between the transmembrane and cytoplasmic signaling domains of the CAR; alanine insertion had previously been shown to improve the *in vivo* anti-tumor efficacy of these CARs^29^. The anti-CD19 CAR contained an IgG4 hinge extracellular spacer while the anti-CD20 CARs contained an IgG4 hinge-CH2-CH3 spacer; each spacer was optimized for the target antigen^29, 30^. All CAR proteins contained the identical CD28 transmembrane and cytoplasmic domains and CD3ζ cytoplasmic domain for intracellular signaling. Thus, the CARs in our panel differed only in non-signaling residues.

**Figure 1.**
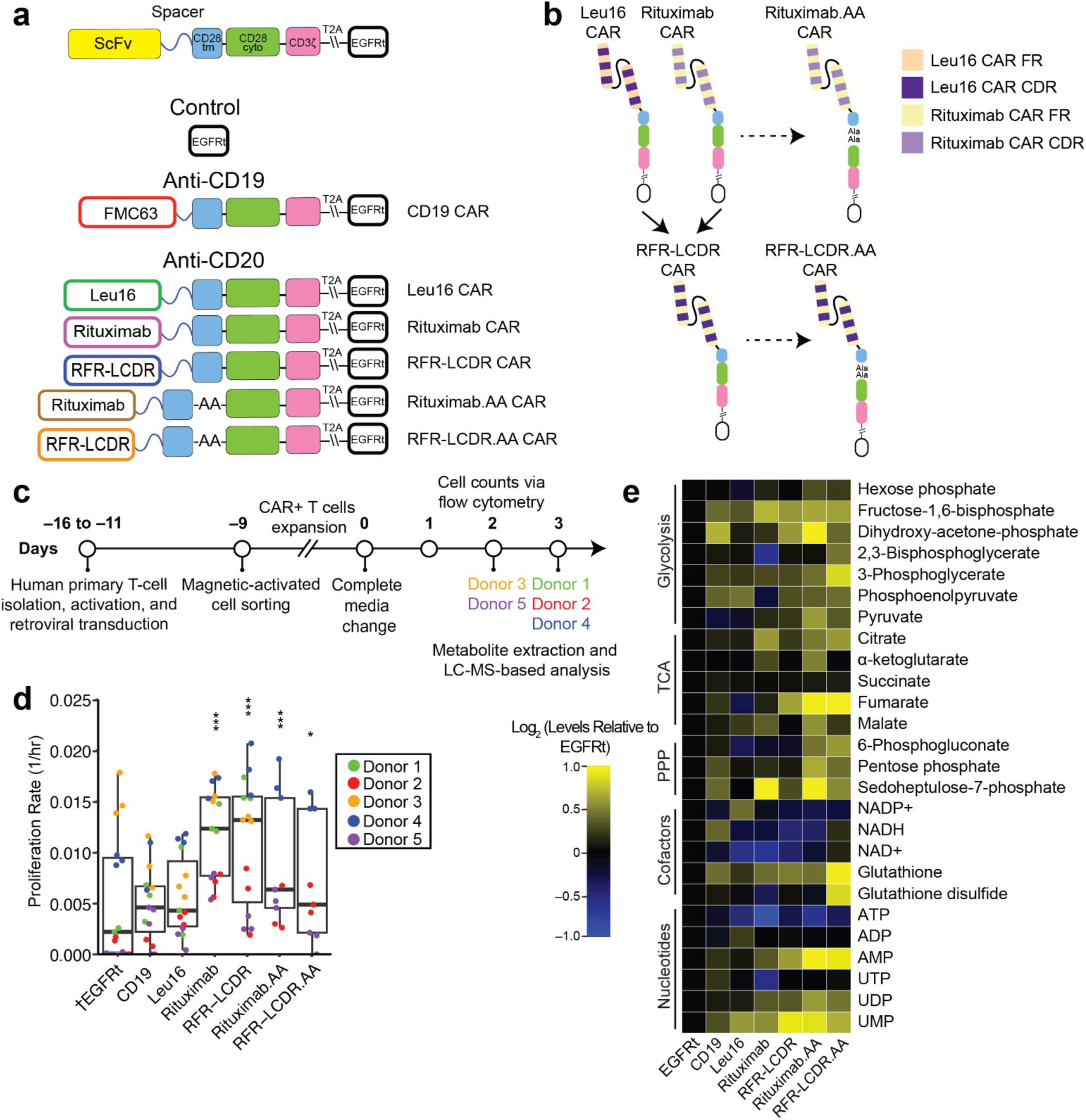
Construction, proliferation, and metabolome of CAR-T cells. **(a)** CAR proteins consist of extracellular domains for antigen binding and intracellular domains for signaling. The scFvs, which determine antigen specificity, were derived from three monoclonal antibodies (FMC63, Leu16, and rituximab) and fused to extracellular spacers followed by CD28 transmembrane (tm) and cytoplasmic (cyto) domains as well as CD3ζ signaling domain. Truncated EGFR (EGFRt) was produced via a self-cleaving (T2A) peptide and used as a transduction marker. Two alanine residues were placed between the transmembrane and intracellular domains in Rituximab.AA and RFR-LCDR.AA CARs. **(b)** RFR-LCDR CAR was constructed by hybridizing the framework regions (FR) of rituximab and the complementarity determining regions (CDR) of Leu16. **(c)** The timeline marks T-cell isolation, transduction, expansion, and measurement of proliferation and metabolome. CAR-T cells were generated from human primary T cells between T–16 and T–11 days. T cells were enriched for CAR^+^ expression by magnetic cell sorting and activated using CD3/CD28 Dynabeads on T–9 days. Cells were expanded for 9 days and seeded in fresh RPMI + dFBS media on day 0. Cells were counted for 2 or 3 days, after which their metabolites were extracted for LC-MS analysis. **(d)** Proliferation rates were determined based on the cell number changes between day 1 and the last time point measurements for each donor’s cells. T cells expressing rituximab-based (rituximab, RFR-LCDR, rituximab.AA, and RFR-LCDR.AA) anti-CD20 CARs grew faster than EGFRt control T cells. Each box shows the quartiles. The whiskers extend to the minimum and maximum values that are within 1.5-fold of the interquartile range. **(e)** CAR-T cells showed CAR-dependent metabolite abundances. Within each row, yellow and blue colors indicate higher and lower levels of a metabolite compared to the EGFRt control T cell. Statistical significance of proliferation rates was determined by a linear mixed-effects model (see Methods) in reference to EGFRt control T cell (†). \**p*<0.05, \*\**p*<0.01, \*\*\**p*<0.001.

We evaluated the impact of CAR protein expression on CAR-T cell expansion and metabolism during *ex vivo* culture (**Fig. 1c**). CAR-T cells maintained their cell numbers and viability for at least 20 days *ex vivo*, indicating their health and suitability for metabolic profiling during this period (**Extended Data Fig. 1a**). To assess how different scFvs affect CAR-T cell proliferation, we counted cells each day for up to three days in RPMI media supplemented with 10% dialyzed FBS (dFBS). The use of dFBS disambiguates the metabolic effects of small molecules found in FBS. A mixed-effects model was used to discern the effects of CARs on the proliferation of T cells originating from different donors. All four rituximab-based (i.e., rituximab, rituximab.AA, RFR-LCDR, and RFR-LCDR.AA) CAR-T cells grew significantly faster than EGFRt control T cells (**Fig. 1d**, **Extended Data Fig. 1b**, and **Supplementary Table 1**).

To investigate whether the disparate growth effects correlate with metabolism, we measured intracellular metabolite levels of CAR-T cells by LC-MS. T cells displayed metabolomic profiles that differed with CAR proteins expressed (**Extended Data Fig. 2**). Focusing on central carbon metabolism (**Fig. 1e** and **Supplementary Table 2**), we observed that rituximab-based CAR-T cells had significantly elevated levels of fructose-1,6-bisphosphate (FBP) compared to EGFRt control T cells, reflecting enhanced glycolytic flux^31^ (**Extended Data Fig. 3a**). Rituximab CAR-T cells also possessed higher levels of the TCA cycle intermediates citrate, α-ketoglutarate, and malate than EGFRt control T cells (**Extended Data Fig. 3b**). While all CAR-T cells maintained redox homeostasis, rituximab CAR-T cells had a lower ATP/ADP ratio compared to EGFRt cells (**Extended Data Fig. 3c**). These observations suggested CAR-dependent reprogramming of T-cell energy metabolism.

**Figure 2.**
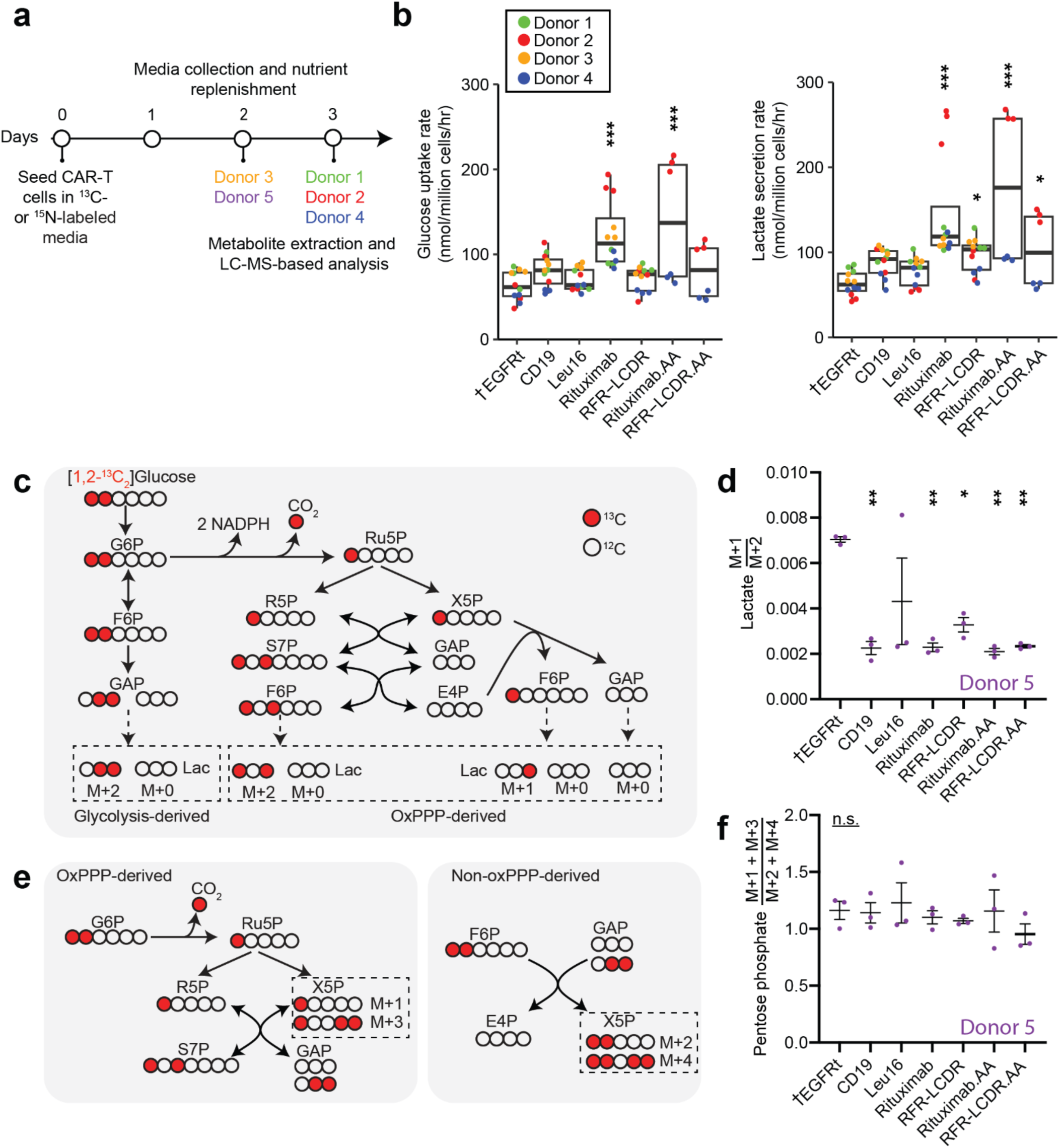
Glycolysis and pentose phosphate pathway activity in CAR-T cells. **(a)** The experimental timeline marks T-cell seeding in isotope-labeled medium, medium collection, and measurement of intracellular and extracellular metabolites. CAR-T cells were seeded in fresh ^13^C-or ^15^N-labeled media at t=0 hours. Media samples were collected for LC-MS analysis between Day 1 and the last time point measurements for each donor’s cells, after which cells were also harvested for intracellular metabolite measurement. No media samples were collected for Donor 5. **(b)** Glucose uptake and lactate secretion rates in rituximab and rituximab.AA anti-CD20 CARs were elevated. Each box shows the quartiles. The whiskers extend to the minimum and maximum values that are within 1.5-fold of the interquartile range. **(c-f)** CAR-T cells were cultured in media containing [1,2-^13^C_2_]glucose for 48 hours. **(c)** Singly ^13^C-labeled (M+1) lactate is only generated through the oxPPP while doubly ^13^C-labeled (M+2) lactate comes from either glycolysis or the oxPPP. **(d)** The (M+1)/(M+2) ratio indicates the activity of oxPPP relative to glycolysis. All but Leu16 CAR-T cells showed lower relative OxPPP activity than EGFRt control T cells. **(e)** Depending on its synthesis route (oxPPP or non-oxPPP), pentose phosphate contains odd or even numbers of ^13^C atoms. Pentose phosphate M+1 and M+3 originate from oxPPP whereas M+2 and M+4 originate from non-oxPPP. **(f)** The odd-to-even ^13^C-labeling ratio of pentose phosphate indicated that the relative usage of oxPPP and non-oxPPP for the nucleotide precursor did not change with CAR expression. Panels (d) and (f) show the mean ± the standard error of the mean (s.e.m.) with n=3. Statistical significance of glucose uptake and lactate secretion rates was determined by using a linear mixed-effects model (see Methods) in reference to the EGFRt control T cell (†). Statistical significance of labeling measurement was determined by two-tailed Student’s *t* test in reference to the EGFRt control T cell (†). \**p*<0.05, \*\**p*<0.01, \*\*\**p*<0.001, n.s. not statistically significant.

**Figure 3:**
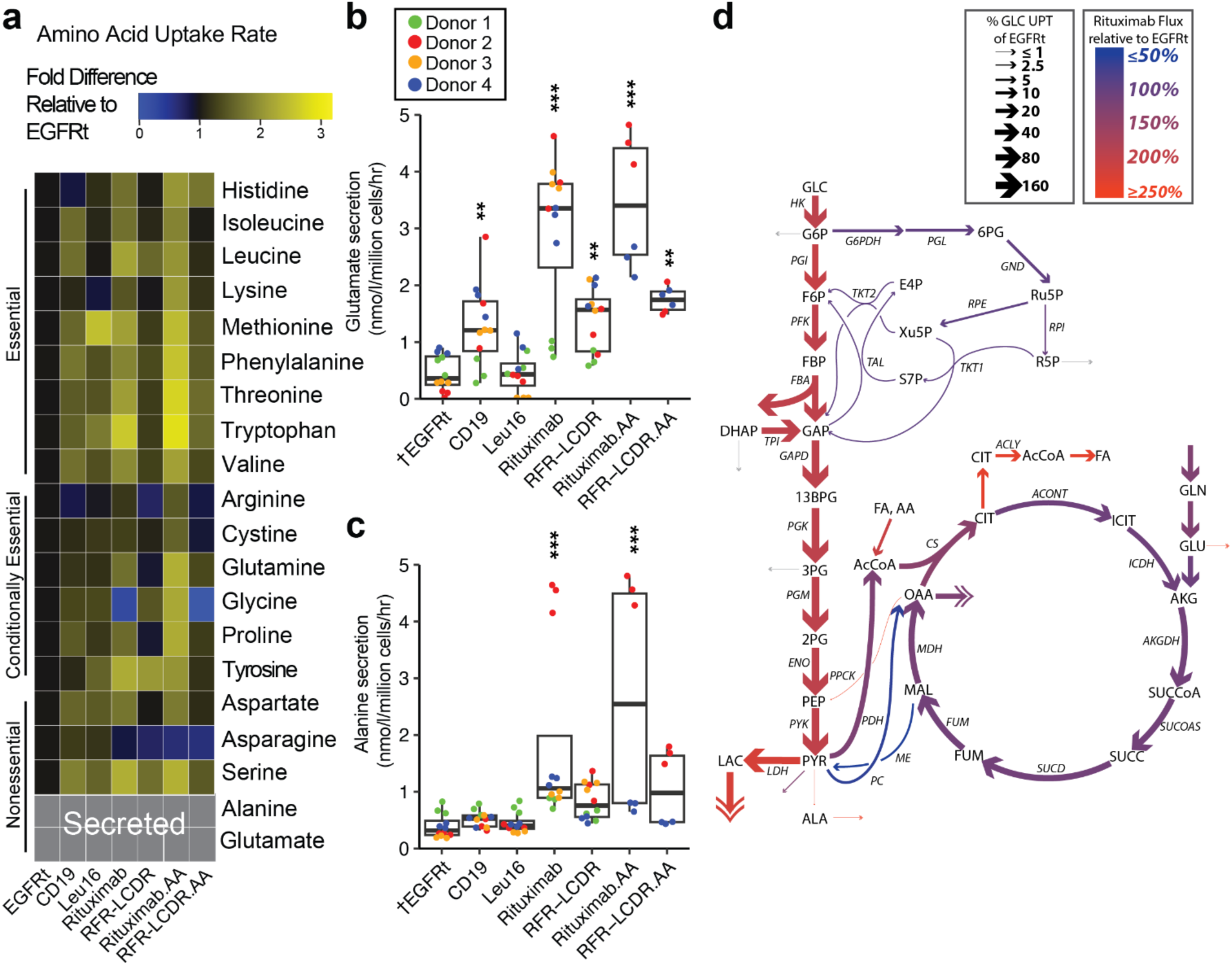
Rituximab-based CD20 CARs increase T-cell metabolic fluxes. **(a)** Amino acid uptake rates were mostly elevated in CAR-T cells compared to EGFRt control T cells. The data represent the mean of measurements from Donors 1-4 cells. **(b)** Glutamate secretion rates were faster in anti-CD19 and rituximab-based anti-CD20 CAR-T cells compared to the EGFRt control T cells. **(c)** Rituximab and rituximab.AA CAR-T cells secreted alanine significantly faster than EGFRt control T cells. Each box in (b) and (c) shows the quartiles. The whiskers extend to the minimum and maximum values that are within 1.5-fold of the interquartile range. **(d)** Metabolic fluxes through central carbon metabolism were compared between EGFRt control T cells and rituximab anti-CD20 CAR-T cells, which showed the largest differences in uptake and secretion rates as well as isotope labeling patterns (**Extended Data** Fig. 5). Metabolic flux analysis was performed using nutrient uptake rates, byproduct secretion rates, and intracellular metabolite labeling patterns in cells that were fed [1,2 ^13^C_2_]glucose, [U-^13^C_6]_glucose, or 50% [U-^13^C_5_^15^N_2_]glutamine, which were obtained from Donors 1-6 cells (see Methods). Arrow widths show the magnitudes of fluxes, normalized to glucose uptake (GLC UPT), for EGFRt T cells. Red and blue colors indicate higher and lower fluxes in rituximab CAR-T cells over EGFRt control T cells, respectively, while gray indicates low confidence flux shifts. Statistical significance for glutamate and alanine secretion in panels (b) and (c) was determined by using a linear mixed-effects model (see Methods) in reference to the EGFRt control T cell (†). \**p*<0.05, \*\**p*<0.01, \*\*\**p*<0.001.

### CAR expression activates aerobic glycolysis

A primary goal of metabolism is to produce energy and biochemical building blocks. Our observation of increased proliferation in the four rituximab-based CAR-T cells implied increased metabolic fluxes. We first compared glycolytic fluxes between CAR-T cells by measuring glucose uptake rates and lactate secretion rates (**Fig. 2a**). Glucose uptake rates doubled in rituximab and rituximab.AA CAR-T cells compared to EGFRt control T cells (**Fig. 2b**). Interestingly, Leu16 and rituximab CARs with 92% scFv sequence identity displayed disparate glucose uptake (**Supplementary Tables 3 and 4**). All four rituximab-based CAR-T cells displayed stronger fermentative glycolysis, converting higher fractions of glucose into lactate, compared to EGFRt control T cells (**Fig. 2b** and **Extended Data Fig. 4a**).

**Figure 4:**
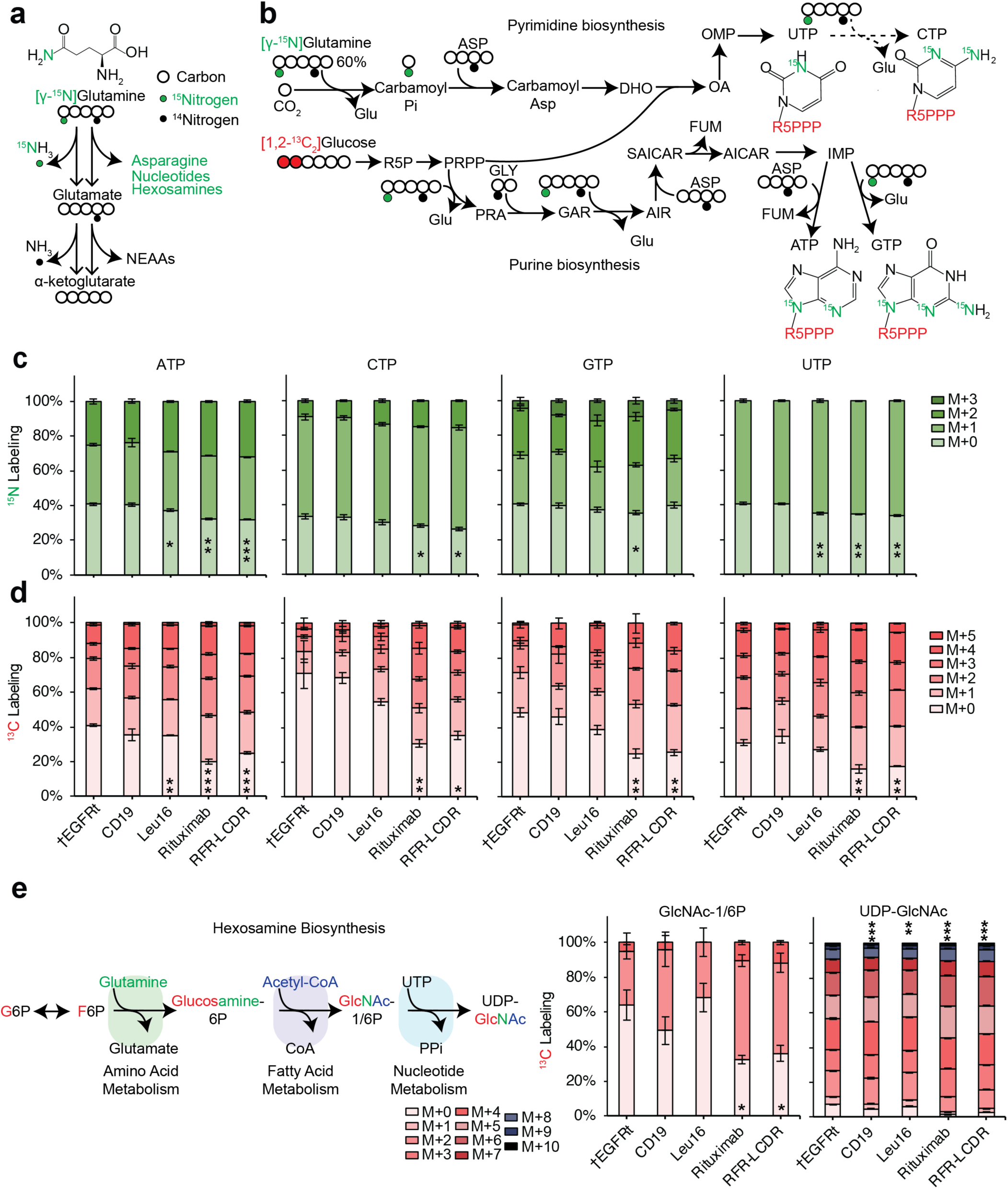
Tracing nitrogen and carbon in nucleotide and hexosamine biosynthesis. **(a)** The fates of ^15^N of [γ-^15^N]glutamine include ammonia, asparagine, nucleotides, and hexosamines. **(b)** Nucleotide biosynthesis map shows how stable isotopes of [γ-^15^N]glutamine and [1,2-^13^C_2_]glucose are incorporated into pyrimidines and purines. **(c)** EGFRt and CAR-T cells from Donor 4 were cultured in media containing [γ-^15^N]glutamine for 72 hours. Nucleotides were labeled more in anti-CD20 CAR-T cells, especially those with rituximab and RFR-LCDR CARs, than in the EGFRt control T cells. **(d-e)** EGFRt and CAR-T cells from Donor 5 were cultured in media containing [1,2-^13^C_2_]glucose for 48 hours. **(d)** Nucleotides were labeled more in rituximab and RFR-LCDR CAR-T cells than in the EGFRt control T cells. The greater labeling fractions in the same (48-and 72-hour) time periods indicated faster nucleotide turnover in rituximab and RFR-LCDR CAR-T cells. **(e)** Hexosamine biosynthesis pathway incorporates glucose carbons and glutamine nitrogen into pathway intermediates. N-acetylglucosamine-1/6-phosphate (GlcNAc-1/6P) and UDP-N-acetylglucosamine (UDP-GlcNAc) were labeled more in rituximab and RFR-LCDR CAR-T cells than EGFRt control T cells, indicating their increased turnover in the CAR-T cells. ^13^C-labeling fractions were corrected for natural isotope abundance and impurities. Panels (c-e) show the mean ± the standard error of the mean (s.e.m.) with n=3. Statistical significance was determined by two-tailed Student’s *t* test in reference to the EGFRt T cell (†) for M+0 labeling. \**p*<0.05, \*\**p*<0.01, \*\*\**p*<0.001, n.s. not statistically significant.

To probe how CAR-T cells utilized glucose, we traced [1,2-^13^C_2_]glucose through glycolysis and the PPP (**Fig 2c**). Glucose-6-phosphate (G6P) is the branching point between glycolysis and the oxidative pentose phosphate pathway (oxPPP), which generates NADPH and the ribose ring of nucleotides. Glycolytic intermediates fructose-6-phosphate (F6P) and glyceraldehyde-3-phosphate (GAP) participate reversibly in the non-oxidative PPP (non-oxPPP). When metabolized through the oxPPP, [1,2-^13^C_2_]glucose uniquely generates singly labeled (M+1) pentose phosphate and triose phosphate, whereas glycolysis generates doubly labeled (M+2) triose phosphate. Thus, the ratio of M+1 to M+2 lactate indicates the oxPPP flux relative to glycolytic flux. Across CAR-T cells, we observed lower fractions of glucose flux being diverted to oxPPP compared to EGFRt control T cells (**Fig. 2d**).

We examined the contribution of oxPPP versus non-oxPPP to pentose phosphate, a precursor to nucleic acid synthesis. Depending on its synthesis route (oxPPP or non-oxPPP), pentose phosphate becomes M+1 or M+2 labeled on its first two carbons (**Fig 2e**). The reversibility of the transketolase reaction can add two ^13^C-labeling in the last three positions of pentose, generating M+3 and M+4 pentose phosphate, respectively. The odd-to-even ratio of pentose phosphate labeling showed that CAR-T cells sourced the ribose ring of nucleotides similarly (**Fig 2f**). Taken together, these observations suggested that CAR-T cells maintain low oxPPP fluxes relative to glycolysis and rituximab-based CAR-T cells turn on aerobic glycolysis.

### CAR expression upshifts nutrient utilization

Cell proliferation is linked to amino acid uptake as proteins account for the majority of cell biomass^32^. We measured the rates of amino acid uptake across CAR-T cells by extracellular metabolomics. All CAR-T cells showed overall faster nutrient uptake than EGFRt control T cells, with rituximab and rituximab.AA CAR-T cells increasing uptake the most (**Fig. 3a**). All T-cell variants consumed glutamine the fastest among all amino acids (**Supplementary Table 5**). Amino acids influence T-cell activation, function, and fate^33^. T-cell activation initially depends on extracellular asparagine although prolonged activation upregulates asparagine synthetase and decreases asparagine dependence^34^. Elevated arginine uptake is linked to superior T-cell survival and anti-tumor activity^7^. Rituximab and rituximab.AA CAR-T cells showed the highest increase in glutamine and arginine uptake along with decreased asparagine dependence, suggesting that they may be poised for rapid activation and initiation of anti-tumor activity.

The altered amino acid uptake profiles pointed to the different abilities of CAR-T cell variants to activate and fight tumors in disparate tumor microenvironments.

All but Leu16 variants of CAR-T cells secreted glutamate faster than EGFRt control T cells (**Fig. 3b**). The increased glutamate secretion suggested that up to 15% of glutamine uptake was “wasted” by CAR-T cells instead of being used for protein synthesis or energy generation (**Extended Data Fig. 4b**). Despite the dependence of T cells on extracellular alanine for protein synthesis^35^, rituximab and rituximab.AA CAR-T cells also secreted alanine at significantly elevated rates compared to control cells (**Fig. 3c**). To see if extracellular alanine would deter alanine secretion, we cultured cells in RPMI-1640 medium supplemented with 10% FBS, which resulted in a final alanine concentration of ∼150 µM (ref.^35^). While anti-CD19 and Leu16 anti-CD20 CAR-T cells, as well as EGFRt control T cells, consumed alanine, all four rituximab-based CAR-T cells still secreted alanine at significant rates (**Extended Data Fig. 4c**). The uptake and secretion measurements across CAR-T cell variants pointed to CAR-dependent nutrient utilization strategies.

We next examined the TCA cycle, the central hub of various nutrient metabolism^36^, by tracing [U-^13^C_6_]glucose and [U-^13^C_5_]glutamine (**Extended Data Fig. 5a,b** and **Supplementary Tables 6 and 7**). Since the effect of alanine insertion (.AA) on nutrient uptake and byproduct secretion was minor compared to that of scFv choices, we henceforth focused on scFv-specific metabolic differences. The ^13^C-labeling fractions of the TCA cycle intermediates from [U-^13^C_6_]glucose revealed that glucose contributed fewer carbons to αKG, which is the first carbonaceous metabolite in glutaminolysis, in rituximab and RFR-LCDR CAR-T cells than EGFRt and other CAR-T cells (**Extended Data Fig. 5c**). Glutamine contributions to the TCA cycle across the CAR-T cells varied to a lesser extent, but αKG gained significantly more ^13^C from [U-^13^C_5_]glutamine in rituximab CAR-T cells (**Extended Data Fig. 5d**). Glutamine contributed more to the TCA cycle metabolites downstream of citrate than glucose did on a carbon basis (**Extended Data Fig. 5e**). The flipped carbon contributions between citrate and αKG suggested substantial diversion of citrate away from the TCA cycle toward fatty acid synthesis.

**Figure 5:**
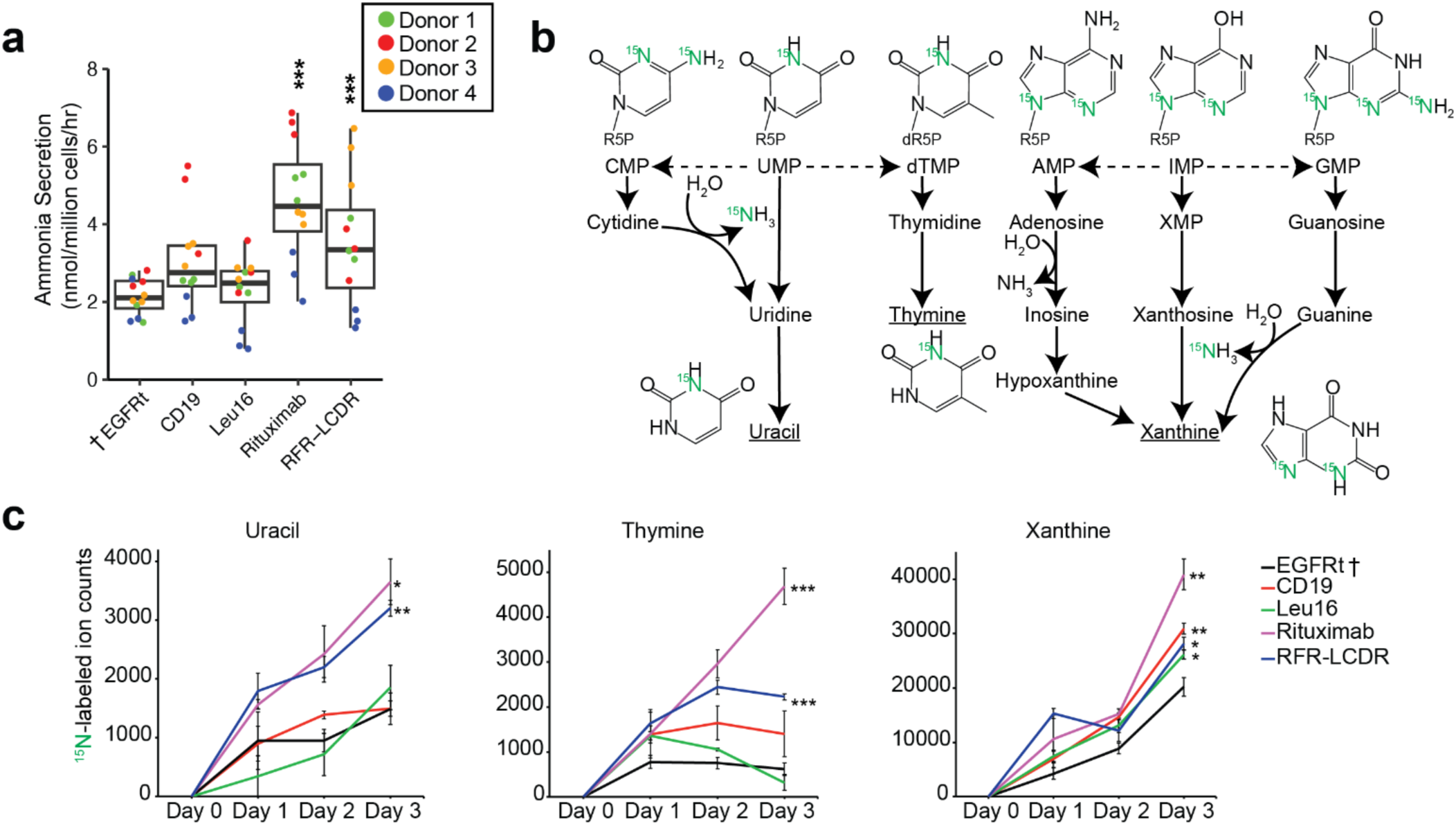
Secretion of nitrogenous metabolites by T cells. **(a)** Ammonia secretion were faster in rituximab and RFR-LCDR CAR-T cells than EGFRt control T cells. Each box shows the quartiles. The whiskers extend to the minimum and maximum values that are within 1.5-fold of the interquartile range. **(b)** Nucleotide degradation pathway shows the fates of ^15^N from [γ-^15^N]glutamine to nucleobases. Uracil, thymine, and xanthine (underlined) were secreted by T cells. **(c)** EGFRt and CAR-T cells from Donor 4 were cultured in media containing [γ-^15^N]glutamine for 72 hours. ^15^N-labeled pyrimidine and purine nucleobases were measured from media samples collected each day. Ion counts for xanthine represent the sum of M+1 and M+2 ^15^N-labeled ions. Plots show the mean ± the standard error of the mean (s.e.m.) with n=3. Statistical significance for ammonia secretion was determined by using a linear mixed-effects model (see Methods) in reference to EGFRt control T cell (†). Statistical significance in panel (c) was determined by two-tailed Student’s *t* test in reference to the EGFRt T cell in Donor 4 (†). \**p*<0.05, \*\**p*<0.01, \*\*\**p*<0.001.

When comparing each CAR-T cell variant against EGFRt control T cells, rituximab CAR-T cells displayed the greatest departure in both ^13^C tracing and transport fluxes (**Fig. 2b**, **Fig. 3a-c**, and **Extended Data Fig. 5**). We thus set out to quantify central carbon metabolism fluxes in these two T-cell variants. For ^13^C-based metabolic flux analysis (^13^C-MFA), we developed a carbon-balanced flux model of central carbon metabolism and glutaminolysis and computed flux values that best fit experimental data for nutrient uptake, byproduct secretion, and ^13^C-labeling patterns in cells that were fed [1,2-^13^C_2_]glucose, [U-^13^C_6_]glucose, or [U-^13^C_5_]glutamine (**Supplementary Tables 8**-**10**). The main intracellular flux in both T-cell variants was glycolysis. Rituximab CAR-T cells extensively utilized aerobic glycolysis (**Fig. 3d**). Compared to EGFRt control T cells, rituximab CAR-T cells lowered the anaplerotic flux through pyruvate carboxylase (PC) to compensate for the faster amino acid uptake and increased the channeling of the acetyl group of acetyl-CoA into fatty acid synthesis rather than oxidizing it. The differences in the TCA cycle turning and the PPP were muted (**Fig. 3d**).

### Rituximab-based CAR-T cells increase nucleotide and hexosamine turnover

We sought to understand the effect of increased amino acid uptake on the nitrogen metabolism of CAR-T cells. Since glutamine was the greatest source of nitrogen for T cells (**Supplementary Table 5**), we cultured CAR-T cells in medium containing 50% [γ-^15^N]glutamine and traced its heavy nitrogen by LC-

MS. The possible fates of nitrogen on the amide of glutamine include proteins, asparagine, nucleotides, hexosamines, urea, and ammonia (**Fig. 4a**). The ^15^N from [γ-^15^N]glutamine is incorporated into both purines and pyrimidines with up to one ^15^N (M+1) in UTP, two ^15^N (M+2) in CTP and ATP, and three ^15^N (M+3) in GTP (**Fig. 4b**). In the same duration (72 hours) of [γ-^15^N]glutamine feeding, all nucleotides in rituximab-based CAR-T cells gained more ^15^N than EGFRt T cells (**Fig. 4c**, **Extended Data Fig. 6a**, and **Supplementary Table 11**). Consistent with the faster turnover of nucleotides in these two CAR-T cells, in the same duration (48 hours) of [1,2-^13^C_2_]glucose feeding, the ribose rings of their nucleotides gained significantly more ^13^C labeling (**Fig. 4d**, **Extended Data Fig. 6b**, and **Supplementary Table 12**). It has been shown that effector T cells have limited pyrimidine biosynthesis capacity, making them sensitive to pyrimidine starvation^37, 38^. Thus, the elevated pyrimidine biosynthesis capacity observed in rituximab-based CAR-T cells would preserve effector T-cell population.

**Figure 6:**
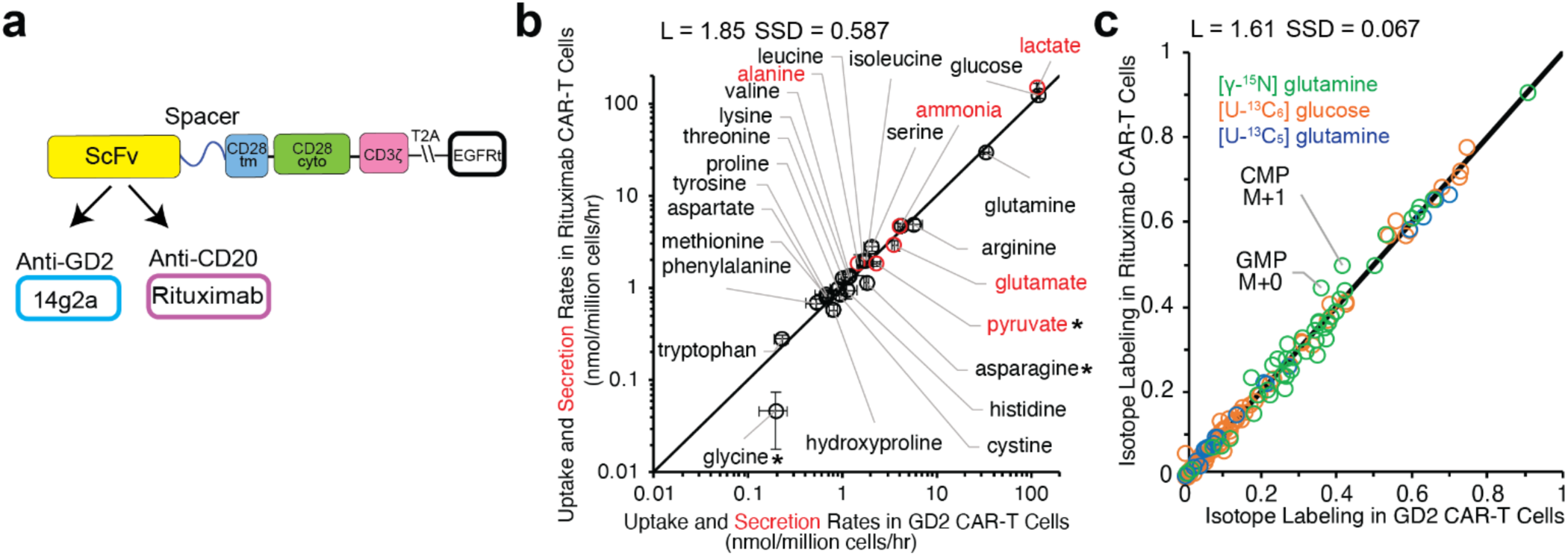
Similarly hypermetabolic rituximab anti-CD20 and 14g2a anti-GD2 CAR-T cells. **(a)** The 14g2a-based anti-GD2 CAR was constructed similarly to the rituximab anti-CD20 CAR, with the scFv as the only difference. **(b)** Rituximab anti-CD20 and 14g2a anti-GD2 (GD2) CAR-T cells from Donors 3 and 4 displayed similar rates of nutrient uptake (black) and byproduct secretion (red). SSD is the sum of squared differences, and L is the total distance between individual points and the line of unity. Plot shows the mean ± the standard error of the mean (s.e.m.) with n=12 for rituximab CAR-T cells and n=6 for GD2 CAR-T cells. **(c)** Metabolites in rituximab and GD2 CAR-T cells, which were fed [γ-^15^N]glutamine, [U-^13^C_6_]glucose, and [U-^13^C_5_]glutamine, were labeled similarly. Metabolites with a distance of at least 0.05 are labeled. The transport fluxes and isotope labeling patterns, which encode intracellular flux information, revealed that anti-GD2 CAR-T cells were metabolically most similar to rituximab CAR-T cells (*cf.* **Extended Data** Fig. 9). Statistical significance in panel (b) was determined by two-tailed Student’s *t* test. \**p*<0.05.

Rituximab and RFR-LCDR CAR-T cells also increased the turnover of hexosamine biosynthesis pathway intermediates N-acetylglucosamine phosphate (GlcNAc-1/6P) and UDP-N-acetylglucosamine (UDP-GlcNAc) (**Fig. 4e**). These metabolites may indicate the cellular nutritional status^39^ as UDP-GlcNAc synthesis requires carbohydrates, glutamine nitrogen, an acetyl group, and a pyrimidine. UDP-GlcNAc is a nucleotide sugar that is a donor substrate for protein and lipid glycosylation^40^. UDP-GlcNAc is also used for O-GlcNAcylation, which regulates signaling pathways and transcription^41, 42^. During early stages of activation, T cells have been shown to increase flux towards O-GlcNAcylation as a regulatory modification to control protein function by increasing the levels of UDP-GlcNAc^43–46^. These observations suggest that increased nutrient uptake by rituximab and RFR-LCDR CAR-T cells may entail systems-level alteration of biochemical networks beyond metabolism.

### Rituximab CAR-T cells operate nitrogen overflow metabolism

Glutamine passes on nitrogen to glutamate and ammonia via the glutaminase reaction (**Fig. 4a**). Consistent with their increased glutamine uptake and glutamate secretion (**Fig. 3a,b**), rituximab and RFR-LCDR CAR-T cells secreted ammonia faster than EGFRt control T cells (**Fig. 5a**). To test whether ammonia generation is reversible (i.e., it can be incorporated back into cellular metabolism), we added ^15^NH_4_Cl to culture media and traced ^15^N (**Extended Data Fig. 7a**). Glutamate dehydrogenase and carbamoyl phosphate synthetase can take ammonia as a substrate to form glutamate and carbamoyl phosphate (as well as downstream amino acids, pyrimidines, and urea). While these metabolites were minimally ^15^N-labeled, we observed small increases in ^15^N labeling of glutamate in anti-CD19 and rituximab-based CAR-T cells (**Extended Data Fig. 7b**) and a decrease in ^15^N labeling of carbamoyl phosphate in rituximab CAR-T cells compared to EGFRt control T cells (**Extended Data Fig. 7c**). The presence of ^15^N in glutamine was more substantial than in glutamate albeit still low. We observed significantly lower ^15^N labeling in glutamine in anti-CD19 and rituximab-based CAR-T cells, which indicated their lower glutamine synthetase activity (**Extended Data Fig 7d**). These results suggested that while CARs affect both the production and the (limited) reincorporation of ammonia by T cells, T cells do not utilize ammonia as a meaningful nitrogen source.

To account for increased amino acid uptake in rituximab and RFR-LCDR CAR-T cells, we searched for other secreted nitrogenous products (**Fig. 5b**). When providing cells with [γ-^15^N]glutamine and measuring their labeling over multiple days, rituximab CAR-T cells produced ^15^N-labeled uracil, thymine, and xanthine the fastest out of all the T cells tested (**Fig. 5c** and **Supplementary Table 13**). Taken together, CAR-T cells with highly upregulated amino acid uptake operate nitrogen overflow metabolism, secreting ammonia, alanine, glutamate, and nucleobases. The inefficient nitrogen metabolism found in such hypermetabolic CAR-T cells may portend poor anti-tumor efficacy, especially if cancer cells also secrete nitrogenous byproducts^47^. The excess pyrimidine biosynthesis capacity, however, may serve as a reserve flux to support differentiation and proliferation on demand in rituximab-based CAR-T cells ^37, 38^.

To discern the effects of secretion products that may accumulate in the tumor microenvironment^48–50^, we treated cells with 300 µM alanine, 800 µM ammonia, 30 mM lactate, or 800 µM ammonia and 30 mM lactate. Extracellular alanine presence exerted CAR-dependent positive and negative effects on proliferation rates (**Extended Data Fig. 8**). Those that absorbed alanine (EGFRt T cells and anti-CD19 and Leu16 anti-CD20 CAR-T cells; **Extended Data Fig. 4c**) proliferated faster with alanine supplementation while the opposite was true for RFR-LCDR CAR-T cells. High ammonia levels have been shown to induce T-cell metabolic reprogramming, increase exhaustion, and decrease proliferation^47^. The addition of ammonia and/or lactate led to slower proliferation for rituximab and RFR-LCDR CAR-T cells, the same ones that elevated the secretion of ammonia and lactate (**Extended Data Fig. 8**). Our observations suggested that the impact of the extracellular environment on T-cell proliferation is CAR-dependent and hypermetabolic CAR-T cells (e.g., rituximab) may self-aggravate proliferation in the tumor microenvironment.

### Anti-GD2 and rituximab anti-CD20 CAR-T cells are similarly hypermetabolic

With rituximab CAR elevating carbon and nitrogen metabolism of T cells the most among the six anti-CD20 and anti-CD19 CARs evaluated, we were curious how it compares against a 14g2a-based anti-GD2 CAR, which has been reported to activate T cells by ligand-independent aggregation of CAR proteins mediated through the FR of scFv^51^. We constructed the anti-GD2 CAR identically to the anti-CD20 CARs, except for the scFv domain, which was derived from the 14g2a antibody (**Fig. 6a**). We compared the metabolic activities of rituximab anti-CD20 CAR-T cells and 14g2a anti-GD2 CAR-T cells by measuring their nutrient uptake and byproduct secretion rates as well as the isotope labeling patterns of intracellular metabolites. Transport fluxes showed some significant differences between the two CAR-T cell variants, but as a whole, their metabolic fluxes were near the line of unity (**Fig. 6b**). The two divergent CAR-T cell variants also exhibited similar metabolite labeling patterns upon feeding [U-^13^C_6_]glucose, [U-^13^C_5_^15^N_2_]glutamine, and [γ-^15^N]glutamine (**Fig. 6c**). Comparing rituximab and 14g2a CAR-T cells, the total distances from the individual fluxes and labeling data points to the line of unity were

1.85 and 1.61, respectively, the smallest among all pair-wise comparisons with 14g2a CAR-T cells (**Extended Data Fig. 9a,b**). The resemblance between rituximab and 14g2a CAR-T cells extended to the secretion of nucleobases (**Extended Data Fig. 9c**). Thus, despite their sequence identity being only 64% (**Supplementary Tables 3 and 4**), 14g2a anti-GD2 CAR-T cells were similarly hypermetabolic as rituximab anti-CD20 CAR-T cells.

Metabolism was reprogrammed across the seven CAR-T cell variants to different extents. While the effect of extracellular domains (i.e., scFvs) was dominant, modifying the intracellular non-signaling component (i.e., two-alanine insertions) exerted small but measurable effects. Rituximab and rituximab.AA CAR-T cells differed in thymine and xanthine secretion (**Extended Data Fig. 9d**). We also observed small but significant differences in nucleotide and hexosamine biosynthesis with and without alanine insertions (**Extended Data Fig. 10**). Our results showed that reprogramming of the basal metabolism of CAR-T cells is prevalent but unpredictable on the basis of non-signaling residues. While CARs that have similar sequences and target the same antigen (e.g., Leu16, rituximab, and RFR-LCDR) can differentially reprogram T-cell metabolism, two CARs that have low sequence identity and target different antigens (e.g., rituximab and 14g2a) do not necessarily result in disparate metabolism.

## Discussion

CARs enable T cells to recognize and attack cancer cells. The most successful CAR-T cell therapies to date target hematological malignancies that express CD19 or BCMA. Expanding the applicability of immunotherapy to other diseases entails the tuning of antigen specificity and affinity as well as intracellular signaling. Development of new CARs involves the combinatorial assembly of peptide domains responsible for these parameters, expressing CAR proteins in T cells, and identifying promising CAR designs through screening in tissue cultures and animal models. Understanding how the domains of CAR proteins impact the intracellular biochemical networks of T cells would streamline this process by providing an in-depth look at T-cell physiology and reducing trial and error.

Metabolism is a network of biochemical reactions that supports T-cell proliferation and anti-cancer activity by supplying cellular energy and biochemical building blocks. Different intracellular co-stimulatory and signaling domains can differentially modulate CAR-T cells’ metabolism and efficacy. CARs have been classified according to modifications in their intracellular signaling domains. First-generation CARs consist of CD3ζ signaling modules alone, and second-and third-generation CARs include the addition of single or multiple costimulatory domains (e.g., 4-1BB and CD28). A study comparing T cells expressing 4-1BB-versus CD28-containing CARs reported that 4-1BB CARs lead to mitochondrial-enriched, memory T cells with superior metabolic capacity, antitumor activity, and persistence; in contrast, CD28 are more likely to yield short-lived effector T cells that rely on heightened glycolysis, a phenotype that could be useful in contexts where long-term CAR effects are intolerable^52^. CAR-T cells that simultaneously targeted two tumor-associated antigens while providing both CD28 and 4-1BB costimulation promote both glycolysis and mitochondrial respiration^53^. T-cell metabolism and fate are clearly influenced by the selection of intracellular signaling domains. CAR ectodomains (e.g., extracellular spacers) have also been shown to affect the anti-tumor efficacy of CAR-T cells despite their lack of intrinsic signaling function^29, 54–56^. However, the effect of the scFv and non-signaling portions of CAR proteins on metabolism remains unexplored. The present work sheds light on systems-level metabolic reprogramming of CAR-T cells that accompanies the modification of the extracellular scFv and intracellular non-signaling portions.

We quantified cell proliferation, metabolic fluxes, and mapped metabolic pathway utilization across seven CAR-T cell variants that only differed in non-signaling portions of CAR proteins. We varied the extracellular components with five different scFvs targeting three different antigens (CD19, CD20, and GD2) and, for two of the anti-CD20 scFvs, we inserted two alanine residues between the transmembrane and intracellular domains. Our work highlighted three important lessons: *i*) mere expression of CAR can increase T-cell proliferation and metabolic activity; *ii*) replacement of non-signaling components of CAR differentially rewires metabolism; and *iii*) CARs with similar scFv sequences can have disparate metabolic profiles while CARs with disparate scFv sequences can lead to similar hypermetabolism in T cells.

All seven CAR-T cell variants increased overall nutrient uptake, and a subset of them also increased proliferation rates. The choice of scFv affected metabolic fluxes to a larger extent than alanine insertion in the intracellular non-signaling portion of CAR. Unexpectedly, we saw substantial alanine production by the four rituximab-based (rituximab, RFR-LCDR, rituximab.AA, RFR-LCDR.AA) CAR-T cells. This observation was in contrast to other anti-CD20 (Leu16) CAR-T cells, anti-CD19 CAR-T cells, and EGFRt control T cells, which consumed alanine, and activated T cells, which have been shown to take in extracellular alanine for protein synthesis^35^. These anomalies hinted at a potential metabolic imbalance in rituximab-based CAR-T cells: cells consumed more nutrients than necessary for biomass, cytokine, and energy generation.

Lactate secretion is the epitome of overflow metabolism, in which cells secrete seemingly useful molecules^57, 58^. Rituximab CAR-T cells, which consumed glucose and amino acids the fastest, secreted excess carbons the fastest through lactate. What about the excess nitrogen? In addition to alanine, we observed increased secretion of glutamate, ammonia, and nucleobases by rituximab CAR-T cells. While the accumulation of ammonia has been shown to have detrimental effects on T cells^47^, how the other nitrogenous molecule buildup affects T cells remains to be seen. The inefficient nitrogen metabolism may undermine the hyperactive CAR-T cells’ ability to proliferate and attack cancer cells in a tumor microenvironment containing high levels of nitrogen waste.

On a fundamental level, our observations showed that the scFv domain has impacts on CAR and CAR-T cells beyond antigen specificity. Although we found that similar non-signaling components of CARs differentially rewire metabolism, different CARs do not necessarily result in different T-cell metabolism. Rituximab anti-CD20 CAR-T cells and 14g2a anti-GD2 CAR-T cells showed similarly hyperactive metabolism despite the low sequence identity. Overall, our observations imply that simply swapping out one scFv for another may entail systems-level rewiring of cellular biochemical networks even if the rest of the CAR protein, including all signaling domains, remain the same. Despite the apparent modularity of the peptide domains of CARs and the widespread practice of combinatorial assembly of the domains to develop new CAR proteins^5^, a holistic assessment of CAR proteins is warranted to learn how different domains within a given CAR molecule interact and how CAR proteins interact with each other and other cellular components.

A limitation of our study was a batch effect that is attributable to both donor-intrinsic differences (e.g., genetic factors, age, sex, diet, etc.) and batch-to-batch differences in CAR-T cell generation and cellular metabolite extraction. To account for donor-to-donor variability, we used mixed-effects models. The measurement of the isotope labeling fractions of intracellular metabolites and extracellular metabolite levels obviates the concerns associated with cellular metabolite extraction. Improvement in human primary T-cell transduction, expansion, and metabolite extraction will enable more sensitive detection of CAR-dependent metabolic reprogramming^59, 60^.

The different metabolic rates and pathway usage across the CAR-T cell variants suggested that different CAR-T cells may function optimally in different nutrient environments^17, 61^, and that the optimal CAR construct for a given tumor may be dependent on the specific tumor microenvironment. CAR-T cells with fast basal metabolic activity could be poised to unleash rapid initial anti-cancer cytotoxicity by swiftly ramping up the supply of necessary cellular machinery and energy. On the other hand, less metabolically active cells may be less effective at killing cancer cells initially but may sustain their effectiveness longer by having a more “malleable” metabolism. With increasing knowledge of the metabolism of clinically successful (and unsuccessful) CAR-T cells, metabolic reprogramming would serve as an important prognostic marker for streamlined CAR design, and metabolic engineering should be an integral task in successful CAR-T cell therapy development.

## Materials and Methods

### Construction of scFvs and CARs

The anti-CD19 scFv was derived from FMC63 monoclonal antibody (mAb)^62^. The plasmid encoding the scFv sequence of rituximab was provided by Dr. Anna M. Wu of UCLA and City of Hope^63^. The plasmid encoding the scFv derived from Leu16 mAb was provided by Dr. Michael C. Jensen of the Seattle Children’s Research Institute^64^. AbYsis was used to obtain the V_L_ and V_H_ sequences of the anti-GD2 scFv from the 14g2a mAb (Protein Data Bank entry 4TUJ). Anti-CD20 CARs were constructed by fusing an scFv (in V_L_-V_H_ orientation), an extracellular IgG4 hinge-CH2-CH3 spacer containing L235E N297Q mutations^56^, the CD28 transmembrane and cytoplasmic domains, the CD3ζ cytoplasmic domain, a T2A self-cleaving peptide sequence, and a truncated epidermal growth factor receptor (EGFRt) with the MSCV backbone. The 14g2a-based anti-GD2 CAR was constructed identically to the anti-CD20 CARs except for the scFv (in V_H-_V_L_ orientation for anti-GD2). The anti-CD19 CAR was constructed with the FMC63-based scFv (in V_L_-V_H_ orientation), an IgG4 hinge extracellular spacer, and the identical transmembrane and cytoplasmic domains as the anti-CD20 and anti-GD2 CARs. The IgG4 hinge for the CD19 CAR was based on previous studies showing different structural requirements for optimal CD19 and CD20 antigen targeting^30, 56^. CARs containing two alanine residues were created using DNA sequences inserted between the CD28 transmembrane and cytoplasmic domains using isothermal assembly. RFR-LCDR and RFR-LCDR.AA anti-CD20 CARs were constructed as gene block fragments (Integrated DNA Technologies) and assembled using digestion-ligation with MreI and BstBI. EGFRt was used as a transduction and sorting marker^29^. Percent identity matrix and sequence homology of scFV was determined using T-Coffee^65^.

### Retrovirus production and human primary CAR-T cell generation

Retrovirus was produced by transiently co-transfecting HEK 293T cells with a plasmid encoding a CAR or control construct and pRD114/pHIT60 virus-packaging plasmids (courtesy of Dr. Steven Feldman of the National Cancer Institute) using linear polyethylenimine (25 kDa; Polysciences). After 48 and 72 hours, the supernatants were collected, filtered using a membrane filter (0.45 μm), and pooled. Peripheral blood mononuclear cells (PBMCs) were isolated from healthy donor blood (courtesy of the UCLA Blood and Platelet Center) by density gradient centrifugation using Ficoll-Paque PLUS (GE Healthcare). On the day of isolation, CD14^−^/CD25^−^/CD62L^+^ naïve/memory T cells were enriched by depleting CD14^+^ and CD25^+^ cells and subsequently enriching CD62L^+^ cells by using microbeads with respective specificity (Miltenyi) and magnetic-activated cell sorting (MACS; Miltenyi)^66^. The naïve/memory T cells were stimulated with CD3/CD28 Dynabeads (ThermoFisher) at a 3:1 cell-to-bead ratio for one week. Two to three days into Dynabeads stimulation, T cells were transduced with the retroviral supernatant. After removing Dynabeads, CAR-T cells and EGFRt control T cells were enriched from the resulting T cell population by staining EGFRt with biotinylated cetuximab (Eli Lilly), labeling with anti-biotin microbeads (Miltenyi), and using MACS. T cells were expanded in RPMI-1640 media supplemented with 10% heat-inactivated FBS (HI-FBS) as well as recombinant human IL-2 (50 U/mL; ThermoFisher) and IL-15 (1 ng/mL; Miltenyi).

### CAR-T cell culture conditions

At the start of experiments for metabolic profiling and proliferation measurement, each CAR-T cell variant was moved to three wells in a 12-well plate containing RPMI-1640 media supplemented with 10% dialyzed fetal bovine serum (dFBS), 50 U/mL IL-2, and 1 ng/mL IL-15 unless otherwise specified. Partial media change was performed by removing a third of consumed media and adding the same volume of fresh media every 24 hours to prevent nutrient depletion. For isotope labeling experiments, CAR-T cells were cultured in media containing [1,2-^13^C_2]_glucose, [U-^13^C_6_]glucose, 50% [U-^13^C_5_^15^N_2_]glutamine, and/or 50% [γ-^15^N]glutamine (Cambridge Isotope Laboratories) for 48-72 hours. Labeled media were prepared from RPMI-1640 without glucose (Gibco) by adding the desired isotopically labeled form of glucose to a final concentration of 2 g/L and by adding the desired isotopically labeled form of glutamine to a final concentration of 0.6 g/L glutamine (resulting in 50% labeled glutamine). To trace nitrogen from ammonia, 800 µM ^15^NH_4_Cl was added to media. For proliferation measurement under different extracellular conditions, 300 µM alanine, 800 µM NH_4_Cl, and/or 30 mM lactate was added to media (**Supplementary Table 14**).

### Proliferation rate measurement

Proliferation rate was measured based on how cell numbers changed over time starting 24 hours after seeding the CAR-T cells in RPMI media supplemented with 10% dFBS. On each day, 10 µL of the well-mixed cultures were mixed with 10 µL of 1% flow cytometry staining buffer, and the mixture was run on a MACSQuant VYB flow cytometer, which reported total cell numbers and viability. The total cell count per well (C) was determined by accounting for the total culture volume in each well. Proliferation rate (µ) was computed using linear regression on log-transformed total per-well cell counts over time throughout the experimental periods of up to 72 hours (**Supplementary Table 1**). If cell counts decreased over time, the apparent proliferation rate was set to 0 hr^−1^:

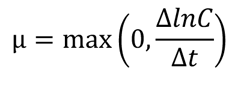

### Metabolite extraction and measurement

Four hours prior to metabolite extraction, culture media was partially replaced with fresh but otherwise identical media to avoid any nutrient depletion. As metabolites turnover fast, metabolism was quenched and metabolites were extracted quickly^67, 68^. Cells were vacuum-filtered onto nylon membrane filters (0.45 μm; Millipore), resting on a glass filter support. Each filter was quickly submerged cell-side down in 400 μL of a –20°C pre-cooled extraction solvent mixture (40:40:20 HPLC-grade acetonitrile/methanol/water) in individual wells of a 6-well plate. The extraction continued for 20 minutes at –20°C. The filters were flipped and washed thoroughly with the extraction solvent. Metabolite extracts were collected into Eppendorf tubes and centrifuged at 17,000 x g in 4°C for 10 minutes. Supernatants were dried under nitrogen gas and reconstituted in HPLC-grade water at 10-15 μL per 1 million cells.

Cell extracts were analyzed on a Vanquish Duo UHPLC system coupled to a Q Exactive Plus orbitrap mass spectrometer (Thermo) by electrospray ionization. The LC separation was performed on a XBridge BEH Amide XP column (150 mm × 2.1 mm, 2.5 μm particle size, Waters, Milford, MA) using a gradient of solvent A (95:5 water/acetonitrile with 20 mM ammonium acetate, 20 mM ammonium hydroxide, pH 9.4) and solvent B (acetonitrile)^69^. Two independent LC flow paths were operated in tandem, staggered by 20 minutes. The gradient for each flow path was 0 min, 90% B; 2 min, 90% B; 3 min, 75% B; 7 min, 75% B; 8 min, 70% B; 9 min, 70% B; 10 min, 50% B; 12 min, 50% B; 13 min, 25% B; 14 min, 25% B; 16 min, 0% B; 20 min, 0% B; 23 min, 0% B; 27 min, 25% B; 30 min, 45% B; 33 min, 90% B; 40 min, 90% B. The flow rate was 150 μL min^−1^. Injection volume was 5 μL, and autosampler and column temperature were 4°C and 25°C, respectively. The MS operated in negative and positive ion modes with a resolution of 140,000 to scan the mass-to-charge (m/z) range of 60-2000. With retention times determined by authenticated standards, resulting mass spectra and chromatograms were identified and integrated using the Metabolomic Analysis and Visualization Engine (MAVEN)^70^ (**Supplementary Tables 2, 6, 7, 9, 11-13**).

### Nutrient uptake and byproduct secretion measurement

Media samples were collected from each culture well every 24 hours during partial replacement of media. Glucose and lactate were measured by YSI7200 instrument (YSI, Yellow Springs, OH) and LC-MS (with internal standards). Ammonia concentration was determined using Ammonia Assay Kit (Sigma-Aldrich AA0100) following manufactured specifications. Assay is based on ammonia reacting with α-ketoglutarate and NADPH in the presence of L-glutamate dehydrogenase to form glutamate and NADP+. Spectrophotometer was used to measure the absorbance at 340 nm, which declines with the oxidation of NADPH, thus revealing ammonia levels. For LC-MS measurement of nutrients and byproducts in media, HPLC-grade methanol was added to fresh and consumed media samples at 80:20 methanol/sample, mixed well, and centrifuged at 17,000g and 4°C for 15 minutes to precipitate proteins. Clear supernatants were collected and analyzed by LC-MS. Metabolite concentrations in culture media were quantified at 24, 48, and 72 hours by normalizing their ion counts to those of fresh media samples with known concentrations. As CAR-T cells did not always proliferate exponentially, uptake and secretion rates were obtained by fitting the rates to measured culture densities and extracellular metabolite concentrations (see Supplementary Information and **Supplementary Table 5**).

### Metabolic flux analysis

EGFRt T cells and rituximab CAR-T cells were labeled with [1,2-^13^C_2_]glucose, [U-^13^C_6_]glucose, or [U-^13^C_5_^15^N_2_]glutamine for 48-72 hours, which is an ample amount of time for central carbon metabolites to reach isotopic steady states. Metabolite labeling measurements were corrected for natural ^13^C abundance and enrichment impurity of labeled substrate. Metabolic fluxes were obtained using the model of central carbon metabolism and glutaminolysis, the elementary metabolite unit (EMU) framework^71^, and mathematical optimization (**Supplementary Table 8**). Measurements of transport fluxes for glucose uptake, glutamine uptake, lactate secretion, glutamate secretion, alanine secretion, and pyruvate secretion were used to constrain the model (**Supplementary Table 5**). The ^13^C labeling patterns of intracellular metabolites contained information on internal metabolic fluxes (**Supplementary Table 9**). Measured M+0 FBP labeling fraction was inexplicably high. Based on FBP aldolase reaction mechanism, which generates comparable fractions of M+0 and M+4 FBP in the reverse reaction^68^, FBP M+0 and M+4 fractions were set equal to each other in the [1,2-^13^C_2_]glucose experiments for ^13^C-MFA. Since central carbon metabolites do not contain nitrogen and our LC-MS has sufficiently high resolution to distinguish between ^13^C and ^15^N labeling, the [U-^13^C_5_^15^N_2_]glutamine labeling experiment was practically the same as [U-^13^C_5_]glutamine tracing. A MATLAB-scripted algorithm was used to simulate mass isotopomer distributions with fluxes as a variable and identify the flux distribution that minimized the variance-weighted sum of squared residuals (SSR) between simulated and measured isotope labeling and uptake and secretion rates:

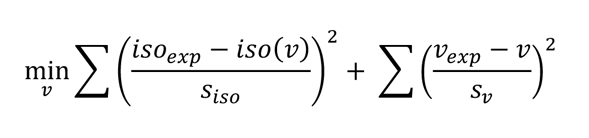

*υ* is the flux vector, *iso_exp_* is the experimentally observed isotope labeling, *iso(υ)* is the simulated isotope labeling, *υ_exp_* is the measured transport fluxes, and *s* denotes the standard deviation of the measured isotope labeling and fluxes. Flux distributions simulated all isotope tracer experiments simultaneously. The confidence interval (95%) of each reaction flux was obtained by searching for the minimum and the maximum flux values that increase the SSR by less than the χ^2^ cutoff (1 degree of freedom) of 3.84 (**Supplementary Table 10**)^72^.

### Statistical analysis

Statistical tests were performed by fitting linear mixed-effects models (LMER) using the statistical software R to the response variables. For each response variable, the linear mixed-effects models had CAR type and donor as the explanatory variables. The mixed-effects models were fit by the restricted maximum likelihood (REML) approach with CAR types as a fixed effect and donors as a random effect:

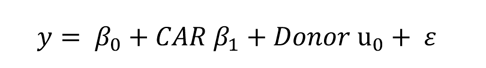

where *y* represents the response variable, *β_0_*, the intercept term, *β_1_*, the fixed-effect regression coefficients for the CAR variable, u_)_, the random effect for the Donor variable, and *ɛ*, the residuals. The random effect contributed to the intercept (*β*_0_), which was set to the reference measurement from EGFRt control T cells, but not the slopes (*β*_1_). With the best parameters that describe the random effects and the residuals, the fitted model computed the coefficients *β*_1_ for the fixed effects of CAR type. Statistical significance for each CAR was obtained by a t test comparing the full model with a simpler model, which contains one fewer parameter that corresponds to the CAR, using Satterthwaite’s method^73^.

## Data availability

Source data for Figures 1-6 are provided in Supplementary Tables 1-14 and the GitHub public repository: https://github.com/aliyalakhani/CAR-T-Cell-Metabolism

## Code availability

The code for metabolic flux analysis and statistical analysis is available on the GitHub public repository: https://github.com/aliyalakhani/CAR-T-Cell-Metabolism

## Acknowledgements

The authors would like to thank the members of the Park lab, the UCLA Metabolomics Center, and the UCLA Molecular Instrumentation Center as well as Drs. M. Nofal, L. B. Tanner, Y. Song, A. M. Wu, M. C. Jensen, and S. Feldman. This work was supported by the National Institute of General Medical Sciences of the National Institutes of Health under Award Number R35GM143127 (J.O.P.), the National Institutes of Health Biotechnology Training Grant under Award Number T32GM067555 (A.L.), the National Institutes of Health Instrumentation Grant Number 1S10OD016387-01, the Alliance for Cancer Gene Therapy Young Investigator Award (Y.Y.C.), the Cancer Research Institute Lloyd J. Old STAR Award (Y.Y.C.), and the UCLA Jonsson Comprehensive Cancer Center Intramural Award (Y.Y.C. and J.O.P.).

## Author Contributions

A.L. and J.O.P. designed the study and wrote the paper with feedback from Y.Y.C. A.L., X.C., L.C.C., M.K., and J.O.P. carried out the experiments. A.L. and J.O.P. analyzed the metabolomic and isotope-labeling results.

## References

1. Ho, P. & Chen, Y. Y. Synthetic Biology in Immunotherapy and Stem Cell Therapy Engineering. in Synthetic Biology 349–372 (Wiley-VCH Verlag GmbH & Co. KGaA, 2018). doi:10.1002/9783527688104.ch17.

2. Brudno, J. N. et al. T Cells Genetically Modified to Express an Anti-B-Cell Maturation Antigen Chimeric Antigen Receptor Cause Remissions of Poor-Prognosis Relapsed Multiple Myeloma. J. Clin. Oncol. 36, 2267–2280 (2018).

3. Cohen, A. D. et al. Safety and Efficacy of B-Cell Maturation Antigen (BCMA)-Specific Chimeric Antigen Receptor T Cells (CART-BCMA) with Cyclophosphamide Conditioning for Refractory Multiple Myeloma (MM). Blood 130, 505–505 (2017).

4. Majzner, R. G. & Mackall, C. L. Clinical lessons learned from the first leg of the CAR T cell journey. Nature Medicine (2019) doi:10.1038/s41591-019-0564-6.

5. Rafiq, S., Hackett, C. S. & Brentjens, R. J. Engineering strategies to overcome the current roadblocks in CAR T cell therapy. Nat. Rev. Clin. Oncol. 2019 173 17, 147–167 (2019).

6. Miccheli, A. et al. Metabolic profiling by 13C-NMR spectroscopy: [1,2-13C2]glucose reveals a heterogeneous metabolism in human leukemia T cells. Biochimie 88, 437–448 (2006).

7. Geiger, R. et al. L-Arginine Modulates T Cell Metabolism and Enhances Survival and Anti-tumor Activity. Cell 167, 829–842.e13 (2016).

8. Gu, M. et al. NF-κB-inducing kinase maintains T cell metabolic fitness in antitumor immunity. Nat. Immunol. 2021 222 22, 193–204 (2021).

9. Can, E. et al. Noninvasive rapid detection of metabolic adaptation in activated human T lymphocytes by hyperpolarized 13C magnetic resonance. Sci. Reports 2020 101 10, 1–8 (2020).

10. Carr, E. L. et al. Glutamine Uptake and Metabolism Are Coordinately Regulated by ERK/MAPK during T Lymphocyte Activation. J. Immunol. (2010) doi:10.4049/jimmunol.0903586.

11. Wang, R. et al. The Transcription Factor Myc Controls Metabolic Reprogramming upon T Lymphocyte Activation. Immunity (2011) doi:10.1016/j.immuni.2011.09.021.

12. Frauwirth, K. A. et al. The CD28 signaling pathway regulates glucose metabolism. Immunity 16, 769–777 (2002).

13. Michalek, R. D. et al. Estrogen-related receptor-α is a metabolic regulator of effector T-cell activation and differentiation. Proc. Natl. Acad. Sci. U. S. A. (2011) doi:10.1073/pnas.1108856108.

14. Blagih, J. et al. The Energy Sensor AMPK Regulates T Cell Metabolic Adaptation and Effector Responses InVivo. Immunity 42, 41–54 (2015).

15. Ma, E. H. et al. Serine Is an Essential Metabolite for Effector T Cell Expansion. Cell Metab. 25, 345–357 (2017).

16. Ron-Harel, N. et al. Mitochondrial Biogenesis and Proteome Remodeling Promote One-Carbon Metabolism for T Cell Activation. Cell Metab. 24, 104–117 (2016).

17. Klein Geltink, R. I., et al. Metabolic conditioning of CD8+ effector T cells for adoptive cell therapy. Nat. Metab. 2, (2020).

18. Bantug, G. R., Galluzzi, L., Kroemer, G. & Hess, C. The spectrum of T cell metabolism in health and disease. Nature Reviews Immunology vol. 18 19–34 (2018).

19. Menk, A. V. et al. Early TCR Signaling Induces Rapid Aerobic Glycolysis Enabling Distinct Acute T Cell Effector Functions. Cell Rep. (2018) doi:10.1016/j.celrep.2018.01.040.

20. Gerriets, V. A. & Rathmell, J. C. Metabolic pathways in T cell fate and function. Trends Immunol. 33, 168–173 (2012).

21. Buck, M. D., O’Sullivan, D. & Pearce, E. L. T cell metabolism drives immunity. Journal of Experimental Medicine vol. 212 (2015).

22. Vardhana, S. A. et al. Impaired mitochondrial oxidative phosphorylation limits the self-renewal of T cells exposed to persistent antigen. Nat. Immunol. 21, (2020).

23. Guo, Y. et al. Metabolic reprogramming of terminally exhausted CD8+ T cells by IL-10 enhances anti-tumor immunity. Nat. Immunol. 22, (2021).

24. Chmielewski, M., Hombach, A., Heuser, C., Adams, G. P. & Abken, H. T Cell Activation by Antibody-Like Immunoreceptors: Increase in Affinity of the Single-Chain Fragment Domain above Threshold Does Not Increase T Cell Activation against Antigen-Positive Target Cells but Decreases Selectivity. J. Immunol. 173, 7647–7653 (2004).

25. Liu, X. et al. Affinity-Tuned ErbB2 or EGFR Chimeric Antigen Receptor T Cells Exhibit an Increased Therapeutic Index against Tumors in Mice. Cancer Res. 75, 3596–3607 (2015).

26. Mössner, E. et al. Increasing the efficacy of CD20 antibody therapy through the engineering of a new type II anti-CD20 antibody with enhanced direct and immune effector cell - mediated B-cell cytotoxicity. Blood 115, 4393–4402 (2010).

27. Reff, M. et al. Depletion of B cells in vivo by a chimeric mouse human monoclonal antibody to CD20. Blood (1994) doi:10.1182/blood.v83.2.435.bloodjournal832435.

28. Uchiyama, S. et al. Development of novel humanized anti-CD20 antibodies based on affinity constant and epitope. Cancer Sci. 101, 201–209 (2010).

29. Chen, X. et al. Rational Protein Design Yields a CD20 CAR with Superior Antitumor Efficacy Compared with CD19 CAR. Cancer Immunol. Res. OF1–OF14 (2022) doi:10.1158/2326-6066.CIR-22-0504.

30. Zah, E., Lin, M. Y., Anne, S. B., Jensen, M. C. & Chen, Y. Y. T cells expressing CD19/CD20 bi-specific chimeric antigen receptors prevent antigen escape by malignant B cells. Cancer Immunol. Res. 4, 498 (2016).

31. Tanner, L. B. et al. Four Key Steps Control Glycolytic Flux in Mammalian Cells. Cell Syst. 7, (2018).

32. Hosios, A. M. et al. Amino Acids Rather than Glucose Account for the Majority of Cell Mass in Proliferating Mammalian Cells. Dev. Cell (2016) doi:10.1016/j.devcel.2016.02.012.

33. Kelly, B. & Pearce, E. L. Amino Assets: How Amino Acids Support Immunity. Cell Metab. 32, 154–175 (2020).

34. Hope, H. C., et al. Coordination of asparagine uptake and asparagine synthetase expression modulates CD8+ T cell activation. JCI Insight 6, (2021).

35. Ron-Harel, N. et al. T Cell Activation Depends on Extracellular Alanine. Cell Rep. 28, (2019).

36. Kaymak, I. et al. Carbon source availability drives nutrient utilization in CD8+ T cells. Cell Metab. 34, 1298–1311.e6 (2022).

37. Dimitrova, P. et al. Restriction of De Novo Pyrimidine Biosynthesis Inhibits Th1 Cell Activation and Promotes Th2 Cell Differentiation. J. Immunol. 169, (2002).

38. Scherer, S. et al. Pyrimidine de novo synthesis inhibition selectively blocks effector but not memory T cell development. Nat. Immunol. | 24, 26 (2023).

39. Wellen, K. E. & Thompson, C. B. Cellular metabolic stress: Considering how cells respond to nutrient excess. Mol. Cell 40, 323 (2010).

40. Bond, M. R. & Hanover, J. A. A little sugar goes a long way: The cell biology of O-GlcNAc. J. Cell Biol. 208, 869–880 (2015).

41. Love, D. C. & Hanover, J. A. The hexosamine signaling pathway: deciphering the ‘O-GlcNAc code’. Sci. STKE 2005, (2005).

42. Butkinaree, C., Park, K. & Hart, G. W. O-linked β-N-acetylglucosamine (O-GlcNAc): Extensive crosstalk with phosphorylation to regulate signaling and transcription in response to nutrients and stress. Biochim. Biophys. Acta - Gen. Subj. 1800, 96–106 (2010).

43. Golks, A., Tran, T. T. T., Goetschy, J. F. & Guerini, D. Requirement for O-linked N-acetylglucosaminyltransferase in lymphocytes activation. EMBO J. 26, 4368–4379 (2007).

44. Kearse, K. P. & Hart, G. W. Lymphocyte activation induces rapid changes in nuclear and cytoplasmic glycoproteins. Proc. Natl. Acad. Sci. U. S. A. 88, 1701–1705 (1991).

45. Swamy, M. et al. Glucose and glutamine fuel protein O-GlcNAcylation to control T cell self-renewal and malignancy. Nat. Immunol. 17, 712–720 (2016).

46. Donnelly, R. P. & Finlay, D. K. Glucose, glycolysis and lymphocyte responses. Mol. Immunol. 68, 513–519 (2015).

47. Bell, H. N. et al. Microenvironmental ammonia enhances T cell exhaustion in colorectal cancer. Cell Metab. 35, 134–149.e6 (2023).

48. Spinelli, J. B. et al. Metabolic recycling of ammonia via glutamate dehydrogenase supports breast cancer biomass. Science (80-.). 358, 941–946 (2017).

49. Tessem, M. B. et al. Evaluation of lactate and alanine as metabolic biomarkers of prostate cancer using 1H HR-MAS spectroscopy of biopsy tissues. Magn. Reson. Med. 60, 510–516 (2008).

50. Quinn, W. J. et al. Lactate Limits T Cell Proliferation via the NAD(H) Redox State. Cell Rep. 33, 108500 (2020).

51. Long, A. H. et al. 4-1BB costimulation ameliorates T cell exhaustion induced by tonic signaling of chimeric antigen receptors. Nat. Med. (2015) doi:10.1038/nm.3838.

52. Kawalekar, O. U. et al. Distinct Signaling of Coreceptors Regulates Specific Metabolism Pathways and Impacts Memory Development in CAR T Cells. Immunity 44, 380–390 (2016).

53. Hirabayashi, K. et al. Dual Targeting CAR-T Cells with Optimal Costimulation and Metabolic Fitness enhance Antitumor Activity and Prevent Escape in Solid Tumors. Nat. cancer 2, 904–918 (2021).

54. Guest, R. D. et al. The role of extracellular spacer regions in the optimal design of chimeric immune receptors: evaluation of four different scFvs and antigens. J. Immunother. 28, 203–211 (2005).

55. James, S. E. et al. Antigen sensitivity of CD22-specific chimeric T cell receptors is modulated by target epitope distance from the cell membrane. J. Immunol. 180, 7028 (2008).

56. Hudecek, M., et al. The Nonsignaling Extracellular Spacer Domain of Chimeric Antigen Receptors Is Decisive for In Vivo Antitumor Activity. Cancer Immunol. Res. 3, 125–135 (2015).

57. Basan, M. et al. Overflow metabolism in Escherichia coli results from efficient proteome allocation. Nature 528, (2015).

58. Reaves, M. L., Young, B. D., Hosios, A. M., Xu, Y. F. & Rabinowitz, J. D. Pyrimidine homeostasis is accomplished by directed overflow metabolism. Nature 500, (2013).

59. Sheldon, R. D., Ma, E. H., Decamp, L. M., Williams, K. S. & Jones, R. G. Interrogating in vivo T-cell metabolism in mice using stable isotope labeling metabolomics and rapid cell sorting. Nat. Protoc. doi:10.1038/s41596-021-00586-2.

60. Kilgour, M. K. et al. Principles of reproducible metabolite profiling of enriched lymphocytes in tumors and ascites from human ovarian cancer. doi:10.1038/s41596-022-00729-z.

61. Li, X. et al. Navigating metabolic pathways to enhance antitumour immunity and immunotherapy. Nat. Rev. Clin. Oncol. 2019 167 16, 425–441 (2019).

62. Nicholson, I. C. et al. Construction and characterisation of a functional CD19 specific single chain Fv fragment for immunotherapy of B lineage leukaemia and lymphoma. Mol. Immunol. 34, (1997).

63. Zettlitz, K. A. et al. ImmunoPET of Malignant and Normal B Cells with 89Zr-and 124I-Labeled Obinutuzumab Antibody Fragments Reveals Differential CD20 Internalization In Vivo. Clin. Cancer Res. 23, 7242–7252 (2017).

64. Jensen, M., Tan, G., Forman, S., Wu, A. M. & Raubitschek, A. CD20 is a molecular target for scFvFc:ζ receptor redirected T cells: Implications for cellular immunotherapy of CD10+ malignancy. Biol. Blood Marrow Transplant. 4, 75–83 (1998).

65. Madeira, F. et al. Search and sequence analysis tools services from EMBL-EBI in 2022. Nucleic Acids Res. 50, (2022).

66. Zah, E. et al. Systematically optimized BCMA/CS1 bispecific CAR-T cells robustly control heterogeneous multiple myeloma. Nat. Commun. 2020 111 11, 1–13 (2020).

67. Bennett, B. D., Yuan, J., Kimball, E. H. & Rabinowitz, J. D. Absolute quantitation of intracellular metabolite concentrations by an isotope ratio-based approach. Nat. Protoc. 3, (2008).

68. Park, J. O. et al. Metabolite concentrations, fluxes and free energies imply efficient enzyme usage. Nat. Chem. Biol. 12, 482–489 (2016).

69. Wang, L. et al. Peak Annotation and Verification Engine for Untargeted LC-MS Metabolomics. Anal. Chem. 91, (2019).

70. Seitzer, P., Bennett, B. & Melamud, E. MAVEN2: An Updated Open-Source Mass Spectrometry Exploration Platform. Metabolites 12, (2022).

71. Antoniewicz, M. R., Kelleher, J. K. & Stephanopoulos, G. Elementary metabolite units (EMU): A novel framework for modeling isotopic distributions. Metab. Eng. 9, (2007).

72. Antoniewicz, M. R., Kelleher, J. K. & Stephanopoulos, G. Determination of confidence intervals of metabolic fluxes estimated from stable isotope measurements. Metab. Eng. 8, 324–337 (2006).

73. Kuznetsova, A., Brockhoff, P. B. & Christensen, R. H. B. lmerTest Package: Tests in Linear Mixed Effects Models. J. Stat. Softw. 82, 1–26 (2017).

