## Extended Data for "Extracellular Domains of CAR Reprogram T-Cell Metabolism Without Antigen Stimulation"

Extended Data Figures 1-10

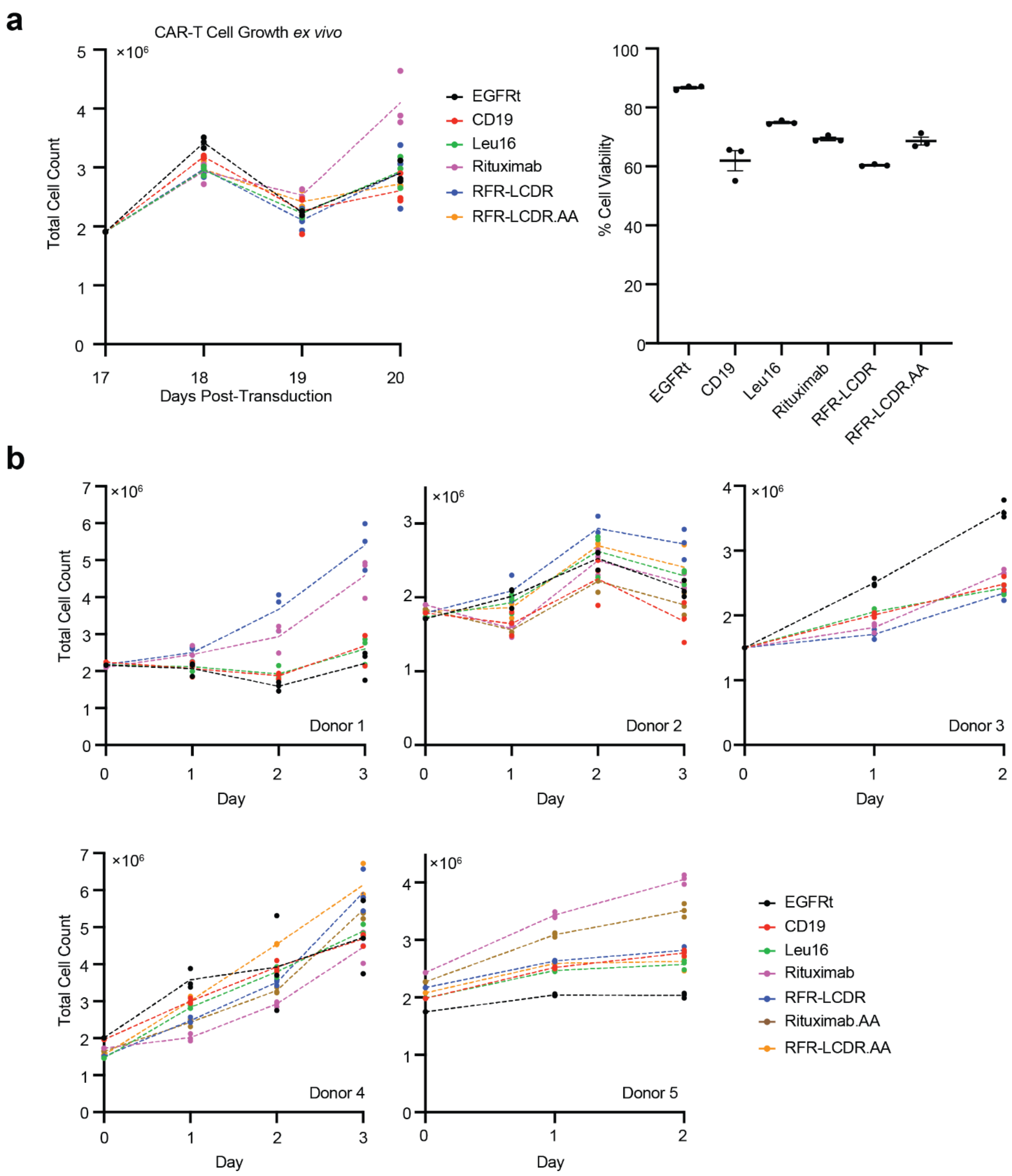


**Extended Data Figure 1. (a)** CAR-T cells are healthy up to 20 days post-retroviral transduction. Graph on left shows proliferation curve of CAR-T cells from days 17 to 20 post-transduction. Graph on right shows viability of CAR-T cells on day 20 post-transduction. **(b)** Growth curves for CAR-T cells engineered from each human donor's cells showed that most CAR-T cells and EGFRt control T cells grew after a complete media change to RPMI-1640 media supplemented with dFBS, IL-2, and IL-15 on day 0. Cell counts between day 1 and the last time point measurements for each donor's cells were used to calculate proliferation rates.

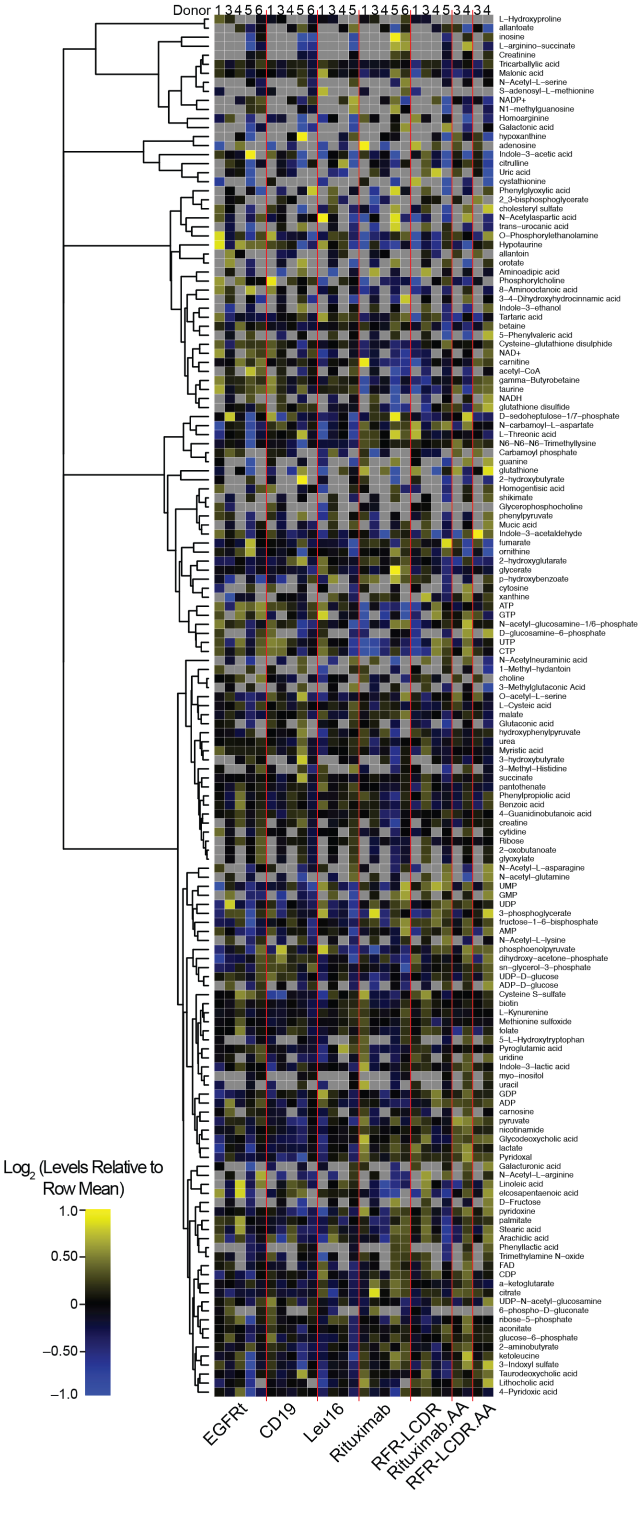


**Extended Data Figure 2.** The metabolome of EGFRt control T cells and CAR-T cells showed CAR-dependent metabolic profiles. To account for batch effects, each metabolomics sample was normalized to its median ion count. Within each row, yellow and blue colors indicate higher and lower levels of a metabolite compared to the row mean of the respective donor CAR-T cells.

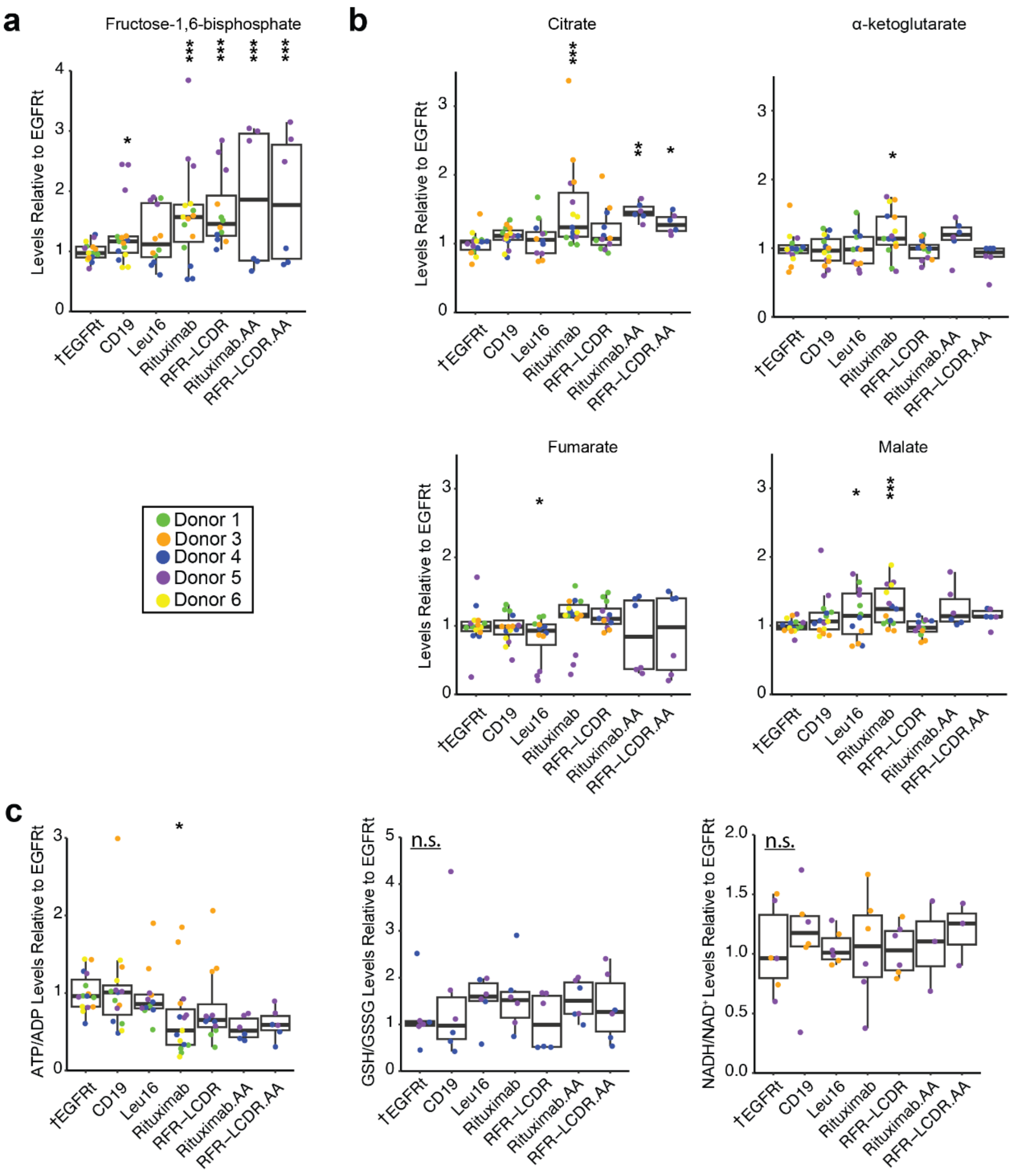


**Extended Data Figure 3.** Central carbon metabolites and co-factor ratios. **(a)** Fructose-1,6-bisphosphate levels were higher in the four rituximab-based CAR-T cells. **(b)** Three of the TCA cycle metabolites were higher in rituximab CAR-T cells **(c)** ATP/ADP was significantly lower in rituximab CAR-T cells. Glutathione/Glutathione disulfide (GSH/GSSG) and NADH/NAD+ ratios showed no statistical difference. Each box shows the quartiles. The whiskers extend to the minimum and maximum values that are within 1.5-fold of the interquartile range. Metabolite levels are normalized to EGFRt, and the ratios are taken from the normalized metabolite levels. Statistical significance was determined by a linear mixed-effects model (see Methods) in reference to EGFRt control T cell (†). *$p$<0.05, **$p$<0.01, ***$p$<0.001, n.s. not statistically significant.

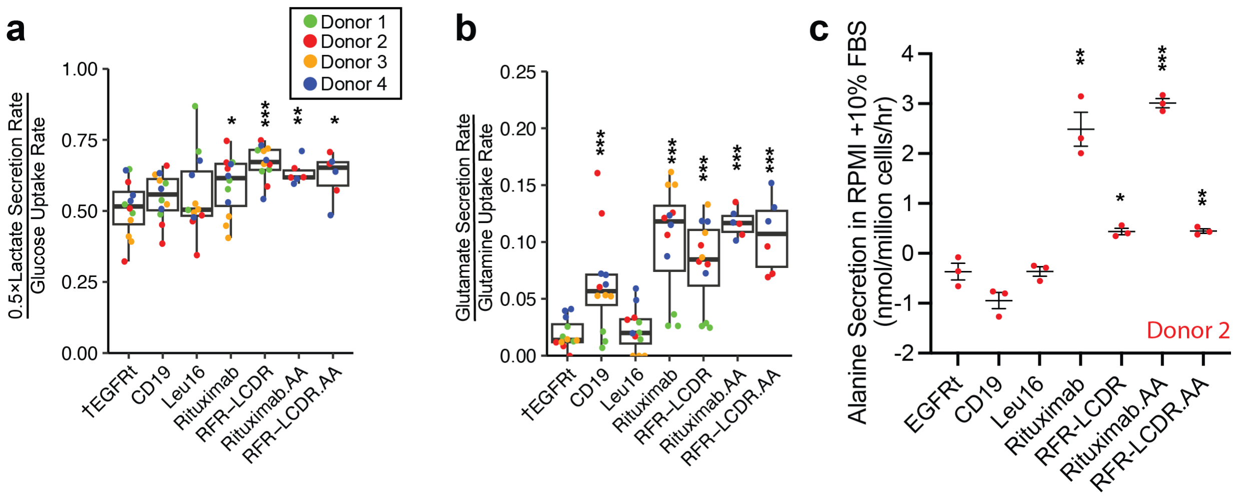


**Extended Data Figure 4. (a)** Higher fractions of glucose carbons were secreted as lactate by the four rituximab-based CAR-T cells than EGFRt control T cells. The lactate-to-glucose carbon flux ratio represents fermentative glycolytic activity. **(b)** In anti-CD19 and rituximab-based CAR-T cells, higher fractions of glutamine were diverted to glutamate secretion. **(c)** Even in the presence of alanine in the media, rituximab-based CAR-T cells secreted alanine at substantial rates unlike EGFRt control T cells, anti-CD19 CAR-T cells, and Leu16 anti-CD20 CAR-T cells, which consumed alanine. Each box in (a) and (b) shows the quartiles. The whiskers extend to the minimum and maximum values that are within 1.5-fold of the interquartile range. Statistical significance in panels (a) and (b) was determined by a linear mixed-effects model (see Methods) in reference to EGFRt control T cell (†). Panel (c) shows the mean ± the standard error of the mean (s.e.m.) with n=3. Statistical significance in panel (c) was determined by two-tailed Student's *t* test in reference to the EGFRt control T cell (†). *$p<0.05$, **$p<0.01$, ***$p<0.001$.

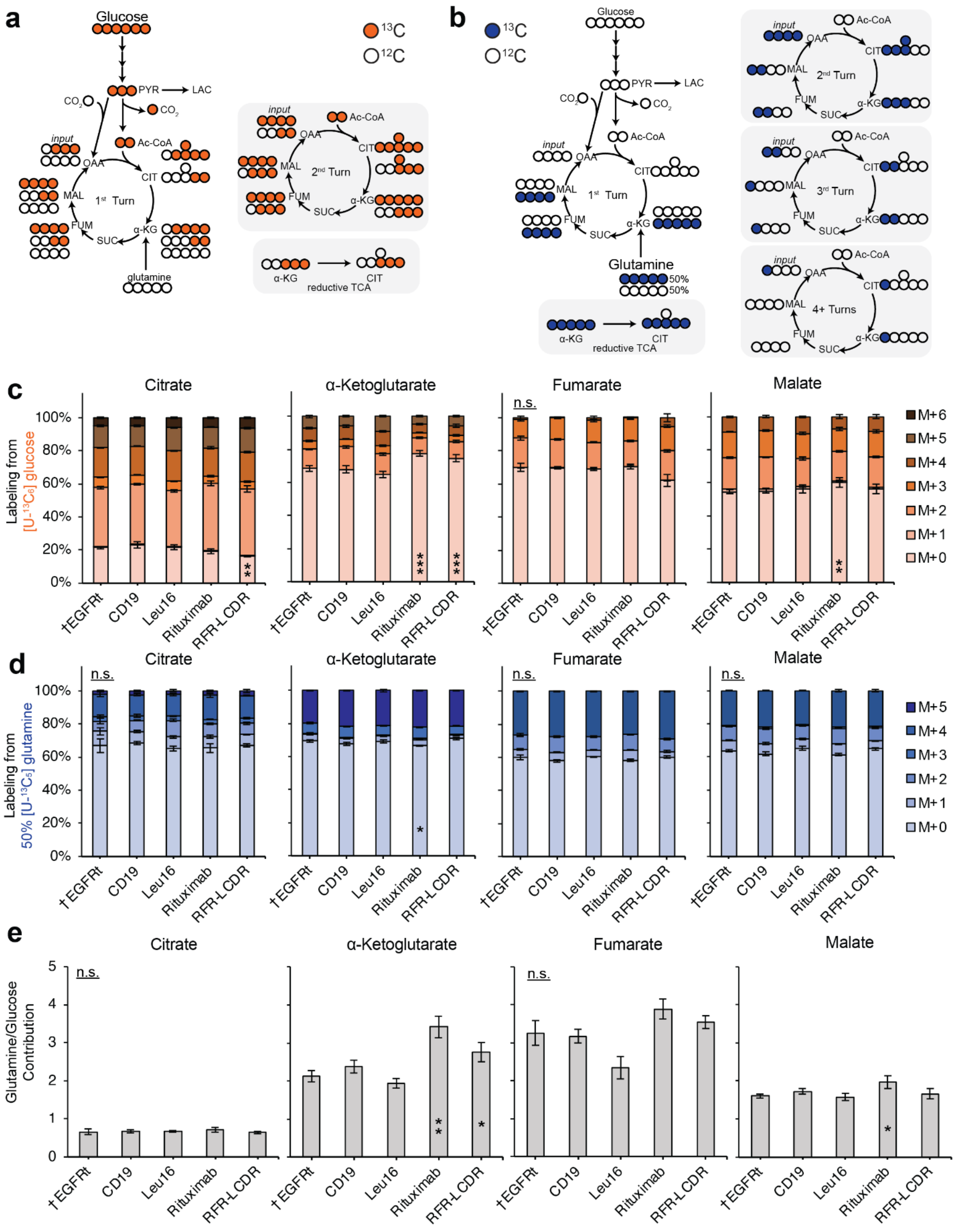


**Extended Data Figure 5.** The TCA cycle activity in CAR-T cells. **(a)** [U-$^{13}C_6$]glucose isotope tracing shows expected labeling patterns in the TCA cycle intermediates. **(b)** [U-$^{13}C_5$]glutamine isotope tracing shows expected labeling patterns in the TCA cycle intermediates. **(c)** The contribution of glucose to α-ketoglutarate and malate was lower in rituximab anti-CD20 CAR-T cells compared to that of EGFRt control T cells based on the significantly higher M+0 fraction in the former. CAR-T cells from Donors 1 and 4 were labeled for 72 hours in media containing [U-$^{13}C_6$]glucose. Labeling fractions were corrected for natural isotope abundance and impurities. **(d)** Glutamine contribution to a-ketoglutarate was significantly higher in rituximab anti-CD20 CAR-T cells than in EGFRt control T cells. CAR-T cells from Donor 3 were labeled for 48 hours in media containing 50% [U-$^{13}C_5$]glutamine. Labeling fractions were corrected for natural isotope abundance and impurities. **(e)** The relative contributions of glucose and glutamine on a carbon basis to the TCA cycle intermediates in the CAR panel indicated that citrate was derived more from glucose but the downstream TCA cycle metabolites were derived mainly from glutamine. Rituximab CAR-T cells displayed higher glutamine-to-glucose contribution ratios compared to EGFRt control T cells. The carbon contributions of glucose and glutamine were obtained by measuring the fractions of the total carbons ($^{12}C$+$^{13}C$) of individual metabolites that were $^{13}C$ and accounting for the enrichment fractions of the respective $^{13}C$ tracers (see Supplementary Notes). Panel (c) shows the mean ± s.e.m. with n=6. Panel (d) shows the mean ± s.e.m. with n=3. Panel (e) shows the mean ± propagated error. Statistical significance in panels (c and d) was determined by two-tailed Student's *t* test in reference to the EGFRt control T cell (†) for M+0 labeling (c-d). Statistical significance in panel (e) was determined by bootstrapping (see Supplementary Notes). *$p$<0.05, **$p$<0.01, ***$p$<0.001, n.s. not statistically significant.

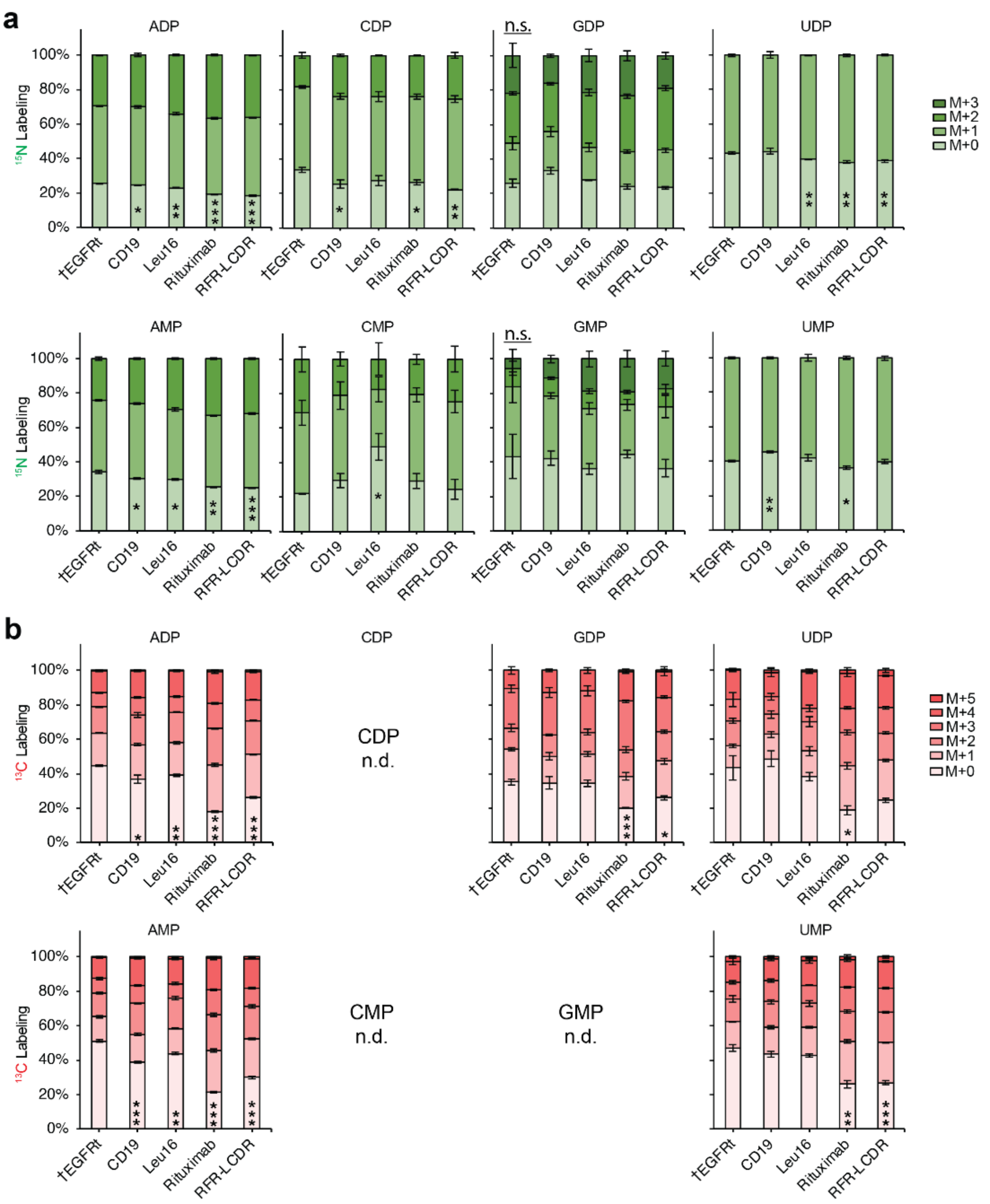


**Extended Data Figure 6.** Tracing nitrogen and carbon reveals nucleotide turnover. **(a)** EGFRt T cells and CAR-T cells from Donor 4 were cultured in media containing 50% [γ-$^{15}N$]glutamine and 50% unlabeled glutamine for 72 hours. Adenosine diphosphate and monophosphate were labeled more in all CAR-T cells than in the EGFRt control T cells. Uridine diphosphate in anti-CD20 CAR-T cells and many of the nucleotides in rituximab CAR-T cells were significantly more labeled were more labeled than those of EGFRt control T cells. **(b)** EGFRt T cells and CAR-T cells from Donor 5 were cultured in media containing [1,2-$^{13}C_2$]glucose for 48 hours. Nucleotide diphosphates and monophosphates were significantly more labeled in rituximab and RFR-LCDR CAR-T cells than in the EGFRt control T cells. The greater labeling fractions in the same (48-

and 72-hour) time periods indicated faster nucleotide turnover in rituximab and RFR-LCDR CAR-T cells. The signals of $^{13}C$-labeled CDP, CMP, and GMP were too low to be reliable (n.d.). $^{13}C$-labeling fractions were corrected for natural isotope abundance and impurities. Plots show the mean ± s.e.m. with n=3. Statistical significance was determined by two-tailed Student's *t* test in reference to the EGFRt control T cell (†) for M+0 labeling. *$p$<0.05, **$p$<0.01, ***$p$<0.001, n.s. not statistically significant.

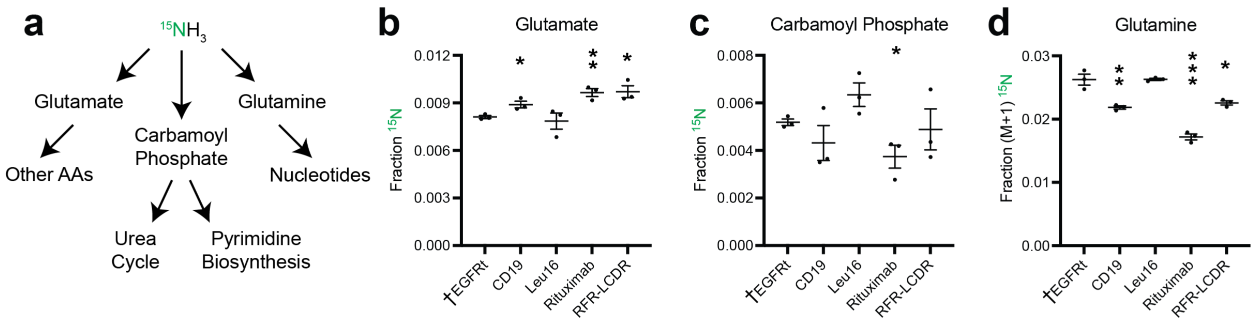


**Extended Data Figure 7.** Ammonia assimilation in CAR-T cells. **(a)** Ammonia can be incorporated into metabolism via glutamate, glutamine, and carbamoyl phosphate. **(b-c)** EGFRt T cells and CAR-T cells from Donor 4 were cultured in media containing 800μM $^{15}NH_4Cl$ for 72 hours. **(b)** The incorporation of $^{15}N$ from the labeled ammonia ($^{15}NH_3$) into glutamate was minimal in all T cells but significantly higher for anti-CD19 and rituximab-based anti-CD20 CAR-T cells compared to EGFRt control T cells. **(c)** Incorporation of $^{15}N$ from $^{15}NH_3$ into carbamoyl phosphate was minimal in all T cells but significantly lower for rituximab CAR-T cells compared to EGFRt T cells. **(d)** Incorporation of $^{15}N$ from $^{15}NH_3$ into glutamine was significantly lower for anti-CD19 and rituximab-based anti-CD20 CAR-T cells compared to EGFRt control T cells. Plots show the mean ± s.e.m. with n=3. Statistical significance was determined by two-tailed Student's *t* test in reference to the EGFRt control T cell (†). $^*p<0.05$, $^{**}p<0.01$, $^{***}p<0.001$.

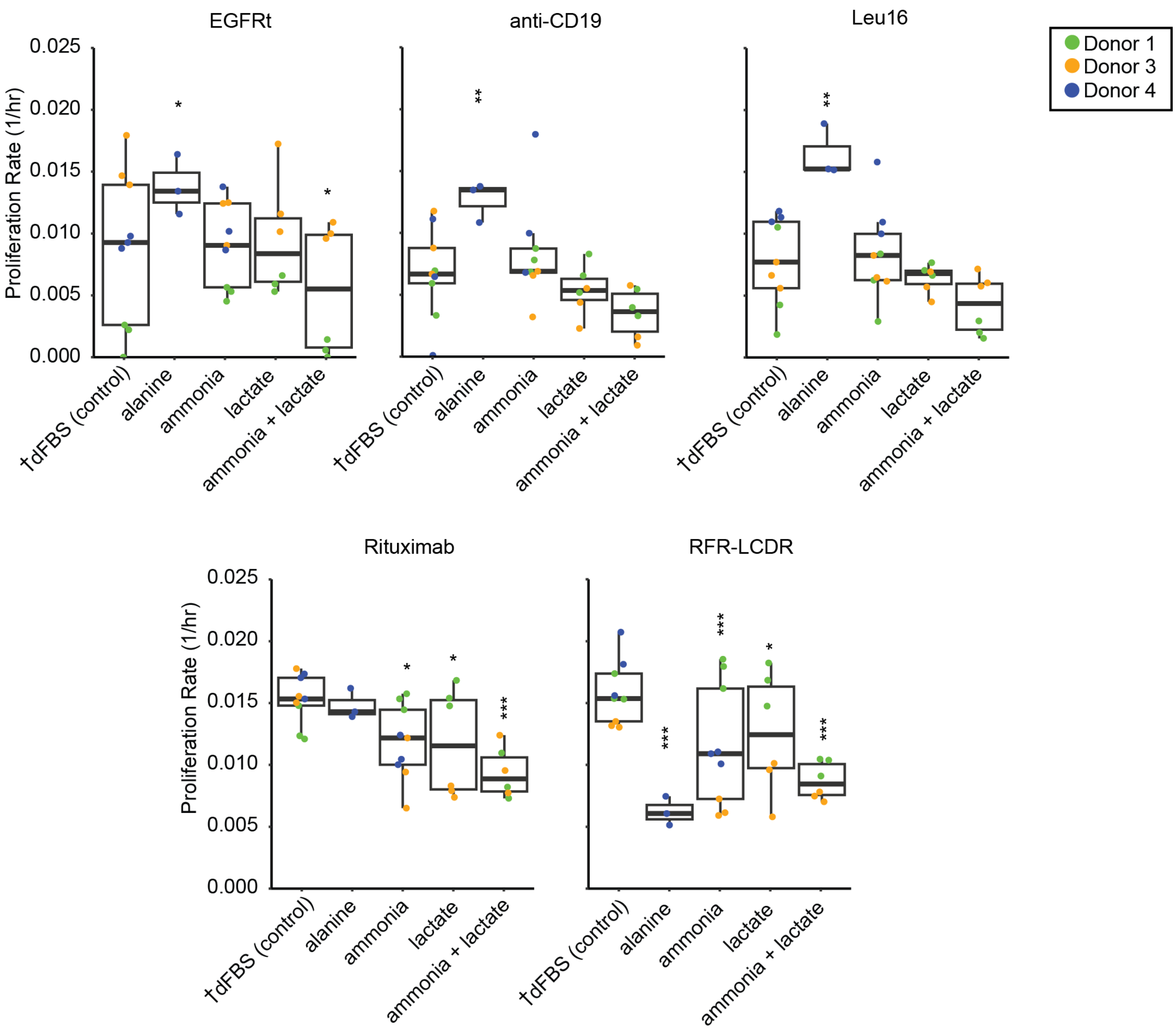


**Extended Data Figure 8.** Proliferation rates for EGFRt control and CAR-T cells grown in various media conditions containing commonly secreted metabolites. EGFRt and CAR-T cells were cultured in media containing 300 μM alanine, 800 μM ammonia, 30 mM lactate, and 800 μM ammonia + 30 mM lactate, which simulate possible product accumulation in the tumor microenvironment. CAR-T cells expressing rituximab and RFR-LCDR anti-CD20 scFvs grew slower in media conditions with ammonia and lactate compared to the base media RPMI1640 supplemented with 10% dialyzed FBS (dFBS). EGFRt T cells as well as anti-CD19 and Leu16 anti-CD20 CAR-T cells grew faster with the alanine addition. Proliferation rates were determined based on the cell number changes between day 1 and the last time point measurements for each donor's cells. Each box shows the quartiles. The whiskers extend to the minimum and maximum values that are within 1.5-fold of the interquartile range. Statistical significance was determined by a linear mixed-effects model (see Methods) in reference to the control media condition (†) for each cell type. *$p$<0.05, **$p$<0.01, ***$p$<0.001.

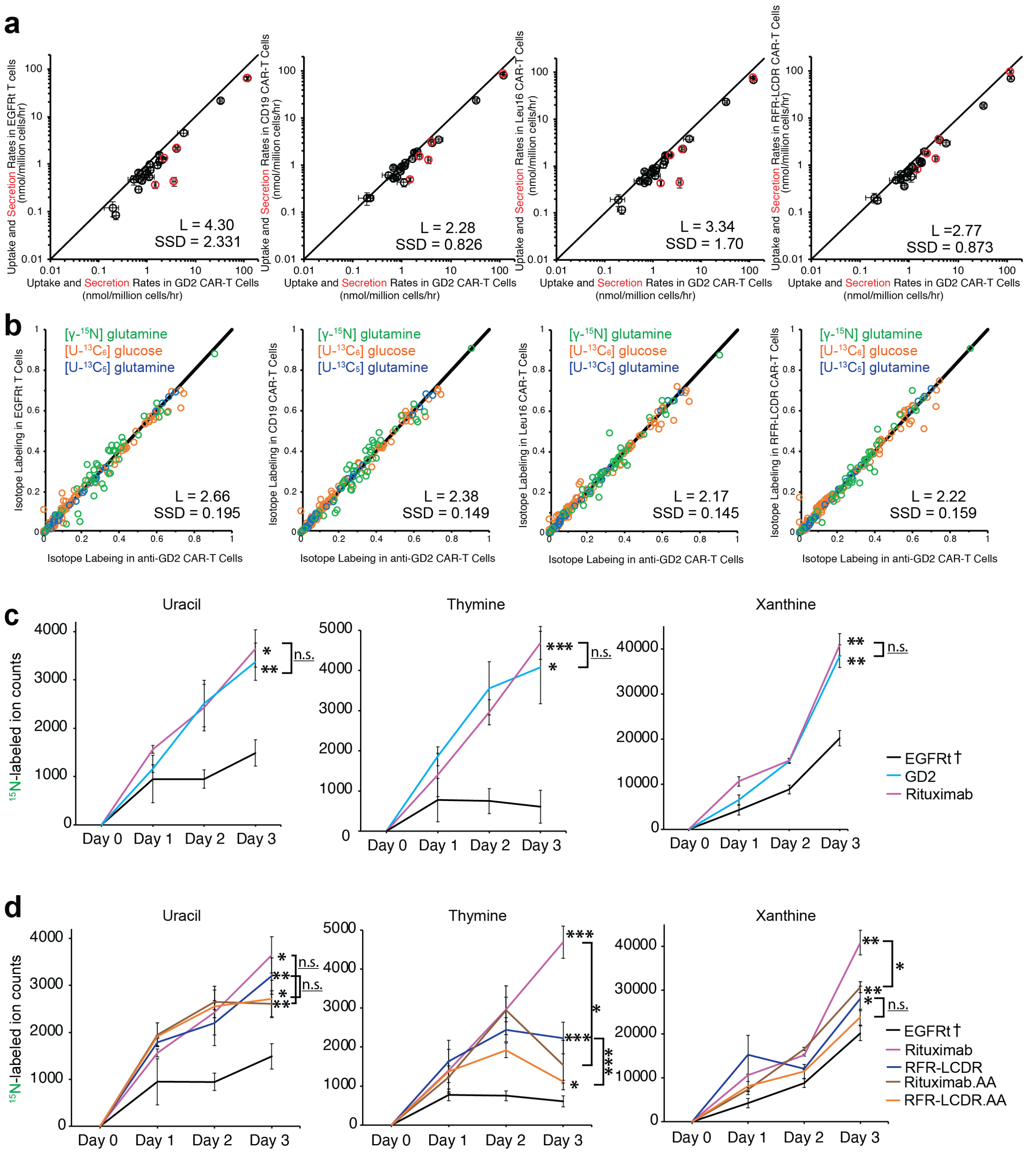


**Extended Data Figure 9.** Comparison of metabolic fluxes across CAR-T cell variants from Donors 3 and 4. **(a)** Anti-GD2 CAR-T cells were more similar to rituximab CAR-T cells and any other CAR-T cell variants in terms of nutrient uptake rates (black) and byproduct secretion rates (red) (*cf.* **Fig. 6b**). SSD is the sum of squared differences, and L is the total distance between individual points and the line of unity. Plot shows the mean ± s.e.m. with n=12 for EGFRt control and anti-CD20 CAR-T cells and n=6 for anti-GD2 CAR-T cells. **(b)** Metabolites labeling patterns in CAR-T cells, which were fed 50% [γ-$^{15}$N]glutamine, [U-$^{13}C_6$]glucose, and [U-$^{13}C_5$]glutamine, showed the greatest similarity between 14g2a-based anti-GD2 CAR-T cells and rituximab CAR-

T cells (*cf.* **Fig. 6c**). Thus, 14g2a-based anti-GD2 CAR-T cells and rituximab CAR-T cells possessed similar metabolic fluxes. **(c-d)** EGFRt T cells and CAR-T cells from Donor 4 were cultured in media containing 50% [γ-$^{15}$N]glutamine for 72 hours. $^{15}$N-labeled pyrimidine and purine nucleobases were measured from media samples collected each day. Ion counts for xanthine represent the sum of M+1 and M+2 $^{15}$N-labeled ions. **(c)** Rituximab anti-CD20 and 14g2a anti-GD2 CAR-T cells displayed similarly high accumulation of purine and pyrimidine nucleobases. **(d)** The four rituximab-based CAR-T cells had faster nucleotide degradation than EGFRt control T cells. Alanine insertions in the non-signaling intracellular portion of CAR proteins resulted in significant differences in thymine secretion. Panels (c) and (d) show the mean ± s.e.m. with n=3. Statistical significance in panels (c-d) was determined by two-tailed Student's *t* test in reference to the day-3 media sample from EGFRt control T cells (†). Further statistical tests were conducted between rituximab and anti-GD2 CAR-T cells, rituximab and rituximab.AA CAR-T cells, and RFR-LCDR and RFR-LCDR.AA CAR-T cells for day-3 nucleobase measurements. *$p<0.05$, **$p<0.01$, ***$p<0.001$, n.s. not statistically significant.

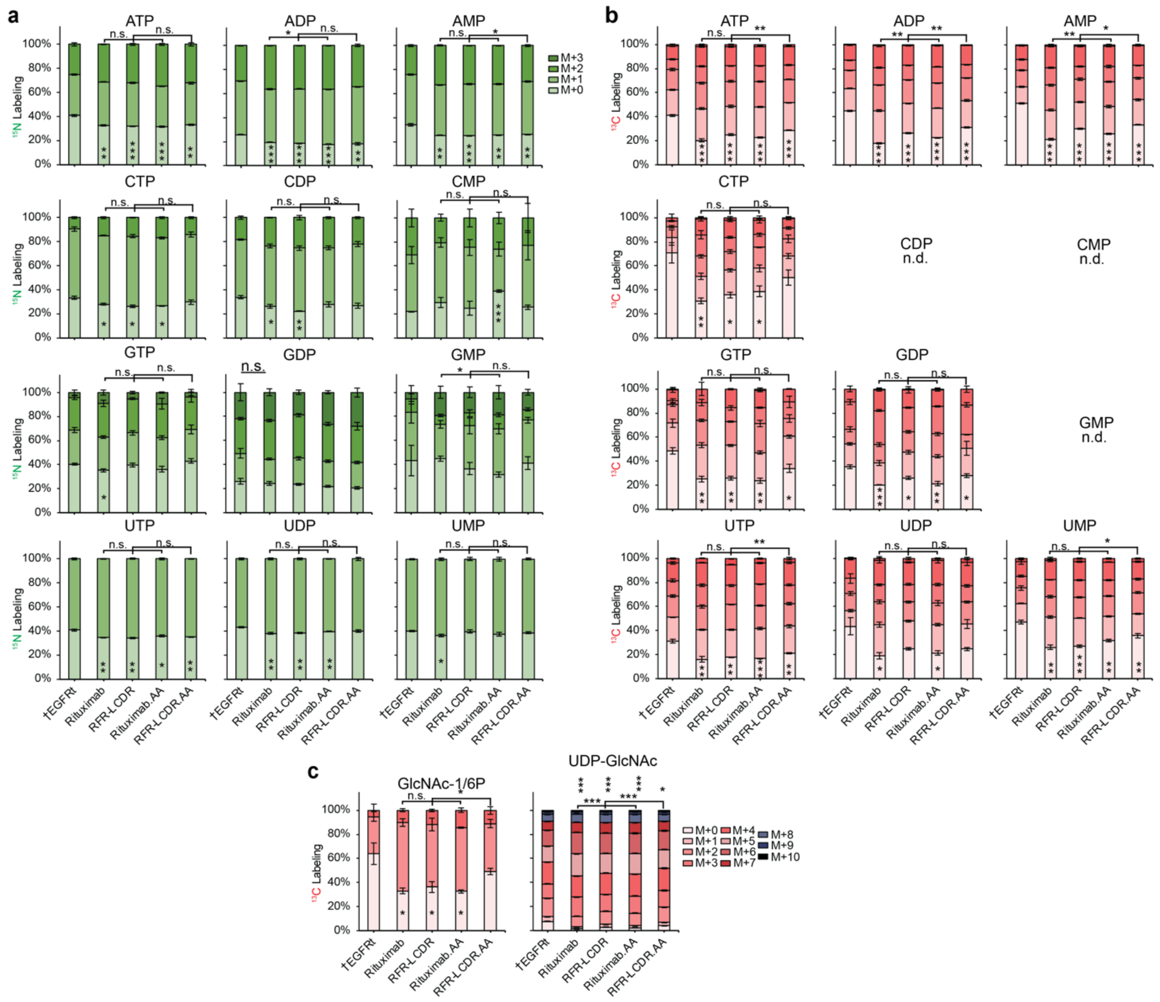

**Extended Data Figure 10.** Tracing nitrogen and carbon in nucleotide and hexosamine biosynthesis for CAR-T cells with two-alanine insertions. **(a)** EGFRt T cells and CAR-T cells from Donor 4 were cultured in media containing 50% [γ-$^{15}$N]glutamine for 72 hours. Many nucleotides were labeled more in rituximab-based CAR-T cells than in the EGFRt control T cells. Alanine insertions resulted in minimal differences. **(b-c)** EGFRt T cells and CAR-T cells from Donor 5 were cultured in media containing [1,2-$^{13}C_2$]glucose for 48 hours. **(b)** Many nucleotides were labeled more in rituximab-based CAR-T cells than in the EGFRt control T cells. Alanine insertions resulted in minimal differences. The signals of $^{13}$C-labeled CDP, CMP, and GMP were too low to be reliable (n.d.). **(c)** N-acetylglucosamine-1/6-phosphate (GlcNAc-1/6P) and UDP-N-acetylglucosamine (UDP-GlcNAc) were labeled more in rituximab-based CAR-T cells than in EGFRt control T cells. Rituximab.AA and RFR-LCDR.AA CAR-T cells with alanine insertions increased M+0 fractions of UDP-GlcNAc compared to rituximab and RFR-LCDR CAR-T cells, respectively. $^{13}$C-labeling fractions were corrected for natural isotope abundance and impurities. Panels (a-c) show the mean ± s.e.m. with n=3. Statistical significance of the observed M+0 labeling fractions was determined by two-tailed Student's *t* test in reference to the EGFRt control T cell (†) and between rituximab and rituximab.AA CAR-T cells and between RFR-LCDR and RFR-LCDR.AA CAR-T cells. *$p<0.05$, **$p<0.01$, ***$p<0.001$, n.s. not statistically significant.
