## Supplement for "Extracellular Domains of CAR Reprogram T-Cell Metabolism Without Antigen Stimulation"

Supplementary Notes 1-4

Supplementary Tables 1-14

### Supplementary Notes

#### 1. Derivation of an equation for exponential cell growth

Cell numbers $C(t)$ (in million cells) is expressed as a function of time, $t$ (in hr), and proliferation rate, $k$ (in $hr^{-1}$).

$$\frac{dC(t)}{dt} = C(t)k \tag{1}$$

$$\int \frac{dC(t)}{C(t)} = \int kdt \tag{2}$$

$$\ln C(t) = kt + constant \tag{3}$$

We used the boundary condition in which at time $t_0$, the cell number is $C_0 = C(t_0)$ to solve for the constant of integration.

$$\ln C(t) = kt + \ln C_0 - kt_0 \tag{4}$$

By rearranging **eqn. 4**, we obtained:

$$\ln\left(\frac{C(t)}{C_0}\right) = k(t - t_0) \tag{5}$$

Using logarithmic identity, we solved for $C(t)$.

$$C(t) = C_0 e^{k(t-t_0)} \tag{6}$$

Based on these equations, to obtain proliferation rate $k$ using cell counts measured at two time points $t_1$ and $t_2$, the following equation was used:

$$k = {\ln\left({C(t_2)}/{C(t_1)}\right)}/{(t_2 - t_1)} \tag{7}$$

When cell counts were measured at three or more time points, $k$ was obtained by finding the slope of the fitted line on measured $\ln C$ versus the corresponding $t$ by linear regression. If cell numbers decreased, $k$ was regarded as 0.

#### 2. Derivation of an equation for nutrient uptake and byproduct secretion rates

The level of an extracellular metabolite (nutrient or byproduct) $P(t)$ (nmol) can be expressed as a function of time, $t$ (hr), secretion or uptake rate, $R$ (nmol/million cells/hr), and cell number, $C(t)$ (million cells). R>0 for secretion and R<0 for uptake by cells. We substituted the solution from the derivation of exponential cell growth, **eqn. 6**, to yield the following differential equation:

$$\frac{dP(t)}{dt} = RC(t) = RC_0 e^{k(t-t_0)} \tag{8}$$

If $k\Delta t$ is small (<1) or if we use live cell counts instead for $C(t)$, because dead cells do not contribute to nutrient uptake or byproduct secretion, linear approximation of $C(t)$ (i.e., first-order Taylor expansion around $t_0$) can be used in the differential equation:

$$\frac{dP(t)}{dt} = R(C_0 + C_0k(t - t_0)) \tag{9}$$

$$P(t) = RC_0(t + \frac{kt^2}{2} - kt_0t) + constant \tag{10}$$

We used the boundary condition in which at time $t_0$, the production rate is $P_0 = P(t_0)$ to solve for the constant of integration.

$$P(t) = RC_0\left[(t - t_0) + \frac{k(t - t_0)^2}{2}\right] + P_0 = R(t - t_0)C_0\left[1 + \frac{k(t - t_0)}{2}\right] + P_0 \tag{11}$$

The following equation describes the average cell number $C_\mathrm{A}$ between $t_0$ and any $t$:

$$C_\mathrm{A} = \frac{C(t)+C(t_0)}{2} = \frac{2C_0+C_0k(t-t_0)}{2} = C_0\left[1+\frac{k(t-t_0)}{2}\right] \tag{12}$$

Using this equation, we obtained the expression for the uptake or secretion rate $R$ using extracellular metabolite levels $P_1 = P(t_1)$ and $P_0 = P(t_0)$ measured at two time points $t_1$ and $t_0$:

$$P_1 - P_0 = RC_\mathrm{A}(t_1 - t_0) = R\frac{C_1+C_0}{2}(t_1 - t_0) \tag{13}$$

#### 3. Calculation of uptake and secretion rates using mathematical optimization

While the uptake and secretion rate calculation between two time points is straightforward, the partial media changes that that we performed every 24 hours rendered the rate calculation more challenging because partial media change replenishes nutrients and dilutes byproducts. We also measured media samples at two or more time points. To account for partial media changes and multiple measurements, we simulated extracellular metabolite levels during the two- to three-day cell culture period. Daily levels of extracellular metabolite levels $P(t)$ were calculated using **eqn. 13** and the volumes of partial media changes at the following points:

*P0: P(t=0 hrs, i.e., fresh media),*
*P1: P(t=24 hrs before media change),*
*P1s: P(t=24 hrs after media change),*
*P2: P(t=48 hrs before media change),*
*P2s: P(t= 48 hrs after media change), and*
*P3: P(t=72 hrs).*

For each nutrient and byproduct, we solved for the uptake or secretion rate, $R$, by simulating the values of these $P$ variables and finding $R$ that resulted in the best agreement with the measured values of $P1$, $P2$, and $P3$, which we obtained by LC-MS. Cell count was taken to be the average cell number between the previous and current days of media collection.

Using **eqn. 13**,

$$P1 = P0 + R\frac{C_1 + C_0}{2}(t_1 - t_0) \tag{14}$$

When the consumed media is partially removed from the culture at 24 hours, the extracellular metabolite level (in nmol) decreases by $\frac{Vr_1}{V_1}P1$, where $\frac{Vr_1}{V_1}$ is the fraction of volume removed. Subsequently, when fresh media is added to the culture, the amount of the extracellular nutrient level increases by $Va_1M_f$, where $Va_1$ is the volume of fresh media added (L) and $M_f$ is the concentration (in nM) of the metabolite in fresh media.

$$P1s = P1 - \frac{Vr_1}{V_1} \cdot P1 + Va_1 \cdot M_f \tag{15}$$

The equations for 48- and 72-hour timepoints, with a partial media change at 48 hours were derived similarly:

$$P2 = P1s + R\frac{C_2 + C_1}{2}(t_2 - t_1) \tag{16}$$

$$P2s = P2 - \frac{Vr_2}{V_2} \cdot P2 + Va_2 \cdot M_f \tag{17}$$

$$P3 = P2s + R\frac{C_3 + C_2}{2}(t_3 - t_2) \tag{18}$$

These equations were used to simulate the values of $P$ given $R$. To find $R$ from our experimental results, we started the simulation from the measured $P1 = P1_{obs}$ at $t_1$=24 hours and used mathematical optimization to find the best fit $R$ that minimized the sum of squared residuals between the simulated and measured $P2$ and $P3$ values:

$$\min_R \sum (P2_{obs} - P2(R))^2 + (P3_{obs} - P3\,(R))^2 \tag{19}$$

where $P1_{obs}$, $P2_{obs}$, and $P3_{obs}$ are the measured metabolite levels, and $P2(R)$, and $P3(R)$ are the simulated levels by solving **eqns. 15-18**. For essential and conditionally essential amino acids, $R$ was constrained to be less than or equal to 0. To obtain the global minimum, this process was iterated 150 times with random initial values of R. With this approach, we obtained the uptake and secretion rates for glucose, lactate, pyruvate, amino acids, and ammonia.

### 4. Calculation of glutamine and glucose contribution to the TCA cycle metabolites

The contributions of glucose and glutamine to the TCA cycle metabolites (citrate, α-ketoglutarate, fumarate, and malate) were obtained by finding the fractions of the total carbons ($^{12}C$+$^{13}C$) of individual metabolites that were $^{13}C$ and accounting for the enrichment fractions of the respective $^{13}C$ tracers (i.e., 100% for [U-$^{13}C_6$]glucose and 50% for [U-$^{13}C_5$]glutamine).

$$Met_{13C} = \frac{\sum_{n=0}^{N} n \cdot Fraction_{M+n}}{N} \tag{20}$$

where $Met_{13C}$ is the fraction of all carbons that are $^{13}C$ in the metabolite $Met$, $N$ is the number of carbons in each $Met$ molecule, and $Fraction_{M+n}$ is the measured fraction of M+n mass isotopomer.

Using this formula, we obtained the mean $\overline{Met_{13C,glc}}$ and $\overline{Met_{13C,gln}}$ as well as their standard errors of the mean (s.e.m.) $s_{Glc}$ and $s_{Gln}$ from [U-$^{13}C_6$]glucose and 50% [U-$^{13}C_5$]glutamine, respectively. The ratio of glutamine-to-glucose contribution $(r)$ and its uncertainty $(s)$ were computed using error propagation.

$$r = \frac{\overline{Met_{13C,gln}}}{\overline{Met_{13C,glc}}} \;;\; s = |r|\sqrt{\left(\frac{s_{Glc}}{\overline{Met_{13C,glc}}}\right)^2 + \left(\frac{s_{Gln}}{\overline{Met_{13C,gln}}}\right)^2} \tag{21}$$

Statistical significance for the glutamine-to-glucose contribution ratios between EGFRt T cells and CAR-T cells was determined by bootstrapping. We bootstrapped the distribution of $r_{EGFRt} - r_{CAR}$, the difference between two contribution ratios from EGFRt T cell variant and one of the CAR-T cell variants (CD19, Leu16, Rituximab, and RFR-LCDR), assuming they came from the same population. For each T-cell variant (EGFRt, CD19, Leu16, Rituximab, and RFR-LCDR), we had six measurements from [U-$^{13}C_6$]glucose labeling and three for 50% [U-$^{13}C_5$]glutamine labeling. For each pair of the EGFRt T cell variant and one CAR-T cell variant, we pooled together the 12 measurements of glucose contribution $(Met_{13C,glc})$ to $Met$ and six measurements of glutamine contribution $(Met_{13C,gln})$ to $Met$. From the pooled glucose contribution sample, six were randomly chosen with replacement for EGFRt and another six were randomly chosen with replacement for the CAR-T cell variant (six being the number of replicate measurements from [U-$^{13}C_6$]glucose tracing). From the pooled glutamine contribution sample, three were randomly chosen with replacement for EGFRt and another three for the CAR-T cell variant (three being the number of replicate measurements from 50% [U-$^{13}C_5$]glutamine tracing). The means of the resampled glucose and glutamine contributions ($\overline{Met_{13C,glc}}$ and $\overline{Met_{13C,gln}}$) and their ratio $(r)$ were obtained for the EGFRt T cell variant and for the CAR-T cell variant. The difference between the two ratios was stored. We iterated this process 100,000 times to generate the bootstrapped null distribution of $r_{EGFRt} - r_{CAR}$. We evaluated how likely it is to observe from this distribution the experimentally measured difference between the glutamine-to-glucose contribution ratios of the EGFRt T cell variant and the CAR-T cell variant. We obtained the p value by finding the probability of observing the absolute differences in the bootstrapped distribution at least as large as the experimentally observed absolute difference. This procedure was employed to find significant differences in the glutamine-to-glucose contribution ratios for citrate, α-ketoglutarate, fumarate, and malate.

**Supplementary Table 1.** CAR-T Cell Proliferation Rates

***CAR-T Cell Proliferation Rates (1/hr)***

| Media Condition | Donor | EGFRt | s.e.m. | CD19 | s.e.m. | Leu16 | s.e.m. | Rituximab | s.e.m. | RFR-LCDR | s.e.m. | Rituximab.AA | s.e.m. | RFR-LCDR.AA | s.e.m. | GD2 | s.e.m. |
|---|---|---|---|---|---|---|---|---|---|---|---|---|---|---|---|---|---|
| RPMI +10% dFBS | 1 | 0.0016 | 0.0008 | 0.0053 | 0.0011 | 0.0056 | 0.0026 | 0.0131 | 0.0009 | 0.0160 | 0.0007 | - | - | - | - | - | - |
| | 2 | 0.0010 | 0.0005 | 0.0007 | 0.0004 | 0.0036 | 0.0004 | 0.0069 | 0.0007 | 0.0056 | 0.0019 | 0.0041 | 0.0013 | 0.0053 | 0.0008 | - | - |
| | 3 | 0.0154 | 0.0012 | 0.0089 | 0.0015 | 0.0067 | 0.0006 | 0.0161 | 0.0008 | 0.0133 | 0.0001 | - | - | - | - | 0.0083 | 0.0004 |
| | 4 | 0.0092 | 0.0003 | 0.0058 | 0.0032 | 0.0114 | 0.0002 | 0.0165 | 0.0006 | 0.0181 | 0.0015 | 0.0169 | 0.0011 | 0.0148 | 0.0005 | 0.0076 | 0.0023 |
| | 5 | 0.0001 | 0.0001 | 0.0040 | 0.0005 | 0.0016 | 0.0006 | 0.0069 | 0.0008 | 0.0029 | 0.0004 | 0.0054 | 0.0005 | 0.0013 | 0.0007 | - | - |
| RPMI +10% dFBS + 300 μM alanine | 4 | 0.0137 | 0.0014 | 0.0125 | 0.0009 | 0.0164 | 0.0012 | 0.0148 | 0.0007 | 0.0063 | 0.0007 | 0.0094 | 0.0022 | 0.0115 | 0.0004 | 0.0154 | 0.0002 |
| RPMI +10% dFBS + 800 μM ammonia | 1 | 0.0051 | 0.0003 | 0.0077 | 0.0005 | 0.0059 | 0.0016 | 0.0151 | 0.0004 | 0.0175 | 0.0007 | - | - | - | - | - | - |
| | 3 | 0.0113 | 0.0011 | 0.0054 | 0.0012 | 0.0070 | 0.0006 | 0.0094 | 0.0016 | 0.0065 | 0.0004 | - | - | - | - | 0.0068 | 0.0004 |
| | 4 | 0.0108 | 0.0015 | 0.0114 | 0.0033 | 0.0122 | 0.0018 | 0.0110 | 0.0007 | 0.0107 | 0.0003 | 0.0096 | 0.0025 | 0.0102 | 0.0034 | 0.0086 | 0.0003 |
| RPMI +10% dFBS + 30 mM lactate | 1 | 0.0059 | 0.0004 | 0.0065 | 0.0009 | 0.0071 | 0.0003 | 0.0156 | 0.0006 | 0.0166 | 0.0010 | - | - | - | - | - | - |
| | 3 | 0.0129 | 0.0022 | 0.0039 | 0.0009 | 0.0057 | 0.0007 | 0.0079 | 0.0003 | 0.0085 | 0.0014 | - | - | - | - | 0.0069 | 0.0010 |
| RPMI +10% dFBS + 800 μM ammonia + 30 mM lactate | 1 | 0.0041 | 0.0006 | 0.0006 | 0.0005 | 0.0022 | 0.0004 | 0.0088 | 0.0011 | 0.0100 | 0.0004 | - | - | - | - | - | - |
| | 3 | 0.0026 | 0.0015 | 0.0101 | 0.0004 | 0.0063 | 0.0004 | 0.0099 | 0.0014 | 0.0075 | 0.0002 | - | - | - | - | 0.0085 | 0.0011 |

Values represent the mean and standard error (n=3).

**Supplementary Table 2.** CAR-T Cell Metabolite Levels (Relative to Those of EGFRt T cells)

***CAR-T Cell Metabolite Levels (Levels Relative to EGFRt)***

| Metabolite | Donor | EGFRt | s.e.m. | CD19 | s.e.m. | Leu16 | s.e.m. | Rituximab | s.e.m. | RFR-LCDR | s.e.m. | Rituximab.AA | s.e.m. | RFR-LCDR.AA | s.e.m. | GD2 | s.e.m. |
|---|---|---|---|---|---|---|---|---|---|---|---|---|---|---|---|---|---|
| Hexose Phosphate | 1 | 1.00 | 0.05 | 1.34 | 0.03 | 0.80 | 0.07 | 0.89 | 0.06 | 0.87 | 0.04 | - | - | - | - | - | - |
| | 3 | 1.00 | 0.03 | 0.65 | 0.04 | 0.66 | 0.03 | 0.89 | 0.10 | 0.77 | 0.04 | - | - | - | - | 0.76 | 0.01 |
| | 4 | 1.00 | 0.04 | 1.10 | 0.02 | 1.09 | 0.04 | 1.01 | 0.00 | 1.04 | 0.03 | 1.10 | 0.04 | 1.11 | 0.04 | 0.92 | 0.01 |
| | 5 | 1.00 | 0.05 | 1.18 | 0.00 | 0.99 | 0.16 | 1.64 | 0.31 | 1.61 | 0.07 | 1.59 | 0.09 | 1.49 | 0.17 | - | - |
| | 6 | 1.00 | 0.11 | 0.88 | 0.11 | - | - | 1.04 | 0.16 | - | - | - | - | - | - | - | - |
| Fructose-1,6-bisphosphate | 1 | 1.00 | 0.04 | 1.16 | 0.05 | 1.28 | 0.30 | 1.39 | 0.18 | 1.46 | 0.08 | - | - | - | - | - | - |
| | 3 | 1.00 | 0.06 | 1.21 | 0.04 | 1.15 | 0.09 | 1.46 | 0.11 | 1.45 | 0.18 | - | - | - | - | 1.09 | 0.12 |
| | 4 | 1.00 | 0.14 | 1.01 | 0.09 | 0.73 | 0.06 | 0.61 | 0.07 | 1.17 | 0.07 | 0.80 | 0.06 | 0.88 | 0.08 | 0.69 | 0.05 |
| | 5 | 1.00 | 0.15 | 2.30 | 0.14 | 1.84 | 0.03 | 2.93 | 0.46 | 2.62 | 0.14 | 2.96 | 0.06 | 2.83 | 0.19 | - | - |
| | 6 | 1.00 | 0.09 | 0.81 | 0.07 | - | - | 1.71 | 0.07 | - | - | - | - | - | - | - | - |
| Dihydroxy-acetone-phosphate | 1 | 1.00 | 0.05 | 2.05 | 0.10 | 1.34 | 0.13 | 1.88 | 0.21 | 2.24 | 0.12 | - | - | - | - | - | - |
| | 3 | 1.00 | 0.26 | 1.77 | 0.07 | 1.19 | 0.03 | 1.05 | 0.39 | 1.06 | 0.11 | - | - | - | - | 1.42 | 0.15 |
| | 4 | 1.00 | 0.12 | 1.31 | 0.06 | 1.16 | 0.10 | 1.40 | 0.05 | 1.27 | 0.12 | 1.19 | 0.11 | 0.90 | 0.04 | 1.48 | 0.00 |
| | 5 | 1.00 | 0.21 | 1.24 | 0.32 | 0.58 | 0.12 | 0.82 | 0.08 | 1.00 | 0.31 | 1.41 | 0.19 | 1.47 | 0.28 | - | - |
| | 6 | 1.00 | 0.07 | 0.90 | 0.05 | - | - | 0.53 | 0.10 | - | - | - | - | - | - | - | - |
| 2,3-Bisphosphoglycerate | 1 | - | - | - | - | - | - | - | - | - | - | - | - | - | - | - | - |
| | 3 | 1.00 | 0.25 | 1.14 | 0.07 | 1.32 | 0.20 | 0.53 | 0.12 | 1.07 | 0.16 | - | - | - | - | 1.17 | 0.13 |
| | 4 | 1.00 | 0.04 | 0.97 | 0.09 | 0.75 | 0.13 | 0.76 | 0.16 | 1.01 | 0.06 | 1.03 | 0.05 | 1.36 | 0.12 | 0.81 | 0.15 |
| | 5 | - | - | - | - | - | - | - | - | - | - | - | - | - | - | - | - |
| | 6 | - | - | - | - | - | - | - | - | - | - | - | - | - | - | - | - |
| 3-Phosphoglycerate | 1 | 1.00 | 0.14 | 1.77 | 0.20 | 2.18 | 0.36 | 1.01 | 0.12 | 0.94 | 0.07 | - | - | - | - | - | - |
| | 3 | 1.00 | 0.18 | 0.59 | 0.06 | 0.66 | 0.03 | 1.47 | 0.53 | 0.60 | 0.06 | - | - | - | - | 0.60 | 0.06 |
| | 4 | 1.00 | 0.30 | 0.88 | 0.11 | 0.71 | 0.12 | 0.58 | 0.11 | 1.09 | 0.16 | 0.63 | 0.05 | 0.87 | 0.06 | 0.76 | 0.07 |
| | 5 | 1.00 | 0.58 | 1.68 | 0.55 | 1.06 | 0.21 | 2.12 | 0.38 | 2.22 | 0.59 | 1.83 | 0.52 | 3.22 | 0.76 | - | - |
| | 6 | 1.00 | 0.14 | 0.79 | 0.06 | - | - | 0.87 | 0.07 | - | - | - | - | - | - | - | - |
| Phosphoenolpyruvate | 1 | 1.00 | 0.13 | 1.10 | 0.07 | 2.43 | 0.39 | 1.09 | 0.18 | 1.49 | 0.05 | - | - | - | - | - | - |
| | 3 | 1.00 | 0.32 | 1.86 | 0.15 | 1.35 | 0.13 | 0.52 | 0.22 | 0.98 | 0.12 | - | - | - | - | 1.00 | 0.14 |
| | 4 | 1.00 | 0.27 | 1.00 | 0.12 | 0.70 | 0.14 | 0.75 | 0.11 | 1.14 | 0.11 | 0.83 | 0.07 | 1.17 | 0.22 | 0.81 | 0.11 |
| | 5 | 1.00 | 0.09 | 1.24 | 0.17 | 0.96 | 0.16 | 1.36 | 0.11 | 1.85 | 0.19 | 2.12 | 0.40 | 1.95 | 0.12 | - | - |
| | 6 | 1.00 | 0.11 | 0.70 | 0.07 | - | - | 0.54 | 0.05 | - | - | - | - | - | - | - | - |
| Pyruvate | 1 | 1.00 | 0.05 | 1.14 | 0.06 | 0.90 | 0.06 | 1.79 | 0.11 | 1.61 | 0.03 | - | - | - | - | - | - |
| | 3 | 1.00 | 0.14 | 0.83 | 0.08 | 0.96 | 0.12 | 0.74 | 0.05 | 1.03 | 0.09 | - | - | - | - | 0.91 | 0.08 |
| | 4 | 1.00 | 0.21 | 0.66 | 0.07 | 0.91 | 0.14 | 0.83 | 0.19 | 0.90 | 0.10 | 1.39 | 0.14 | 1.18 | 0.03 | 0.49 | 0.01 |
| | 5 | 1.00 | 0.25 | 0.78 | 0.19 | 0.79 | 0.08 | 1.46 | 0.45 | 1.07 | 0.14 | 1.64 | 0.56 | 1.35 | 0.44 | - | - |
| | 6 | 1.00 | 0.05 | 0.85 | 0.04 | - | - | 0.74 | 0.13 | - | - | - | - | - | - | - | - |
| Citrate | 1 | 1.00 | 0.06 | 1.24 | 0.06 | 1.40 | 0.15 | 1.02 | 0.03 | 0.95 | 0.05 | - | - | - | - | - | - |
| | 3 | 1.00 | 0.22 | 1.21 | 0.05 | 0.86 | 0.11 | 2.49 | 0.45 | 1.53 | 0.26 | - | - | - | - | 1.18 | 0.26 |
| | 4 | 1.00 | 0.03 | 0.98 | 0.09 | 1.09 | 0.07 | 1.17 | 0.04 | 1.27 | 0.10 | 1.46 | 0.05 | 1.35 | 0.09 | 0.85 | 0.02 |
| | 5 | 1.00 | 0.02 | 1.03 | 0.07 | 0.95 | 0.09 | 1.53 | 0.23 | 0.99 | 0.04 | 1.47 | 0.11 | 1.23 | 0.09 | - | - |
| | 6 | 1.00 | 0.10 | 1.02 | 0.09 | - | - | 1.33 | 0.08 | - | - | - | - | - | - | - | - |
| α-ketoglutarate | 1 | 1.00 | 0.04 | 1.12 | 0.09 | 1.22 | 0.16 | 0.94 | 0.13 | 1.05 | 0.08 | - | - | - | - | - | - |
| | 3 | 1.00 | 0.31 | 0.85 | 0.08 | 0.84 | 0.07 | 1.39 | 0.20 | 1.01 | 0.10 | - | - | - | - | 0.97 | 0.07 |
| | 4 | 1.00 | 0.01 | 1.15 | 0.07 | 1.14 | 0.07 | 1.24 | 0.11 | 1.04 | 0.04 | 1.22 | 0.06 | 1.00 | 0.00 | 1.25 | 0.08 |
| | 5 | 1.00 | 0.07 | 0.70 | 0.07 | 0.71 | 0.05 | 1.20 | 0.31 | 0.80 | 0.04 | 1.09 | 0.23 | 0.74 | 0.14 | - | - |
| | 6 | 1.00 | 0.09 | 1.00 | 0.10 | - | - | 1.33 | 0.18 | - | - | - | - | - | - | - | - |
| Succinate | 1 | 1.00 | 0.06 | 1.06 | 0.04 | 1.24 | 0.00 | 1.03 | 0.06 | 0.96 | 0.10 | - | - | - | - | - | - |
| | 3 | 1.00 | 0.24 | 0.85 | 0.04 | 0.96 | 0.05 | 1.07 | 0.08 | 1.21 | 0.11 | - | - | - | - | 1.01 | 0.05 |
| | 4 | 1.00 | 0.14 | 0.97 | 0.12 | 0.97 | 0.10 | 0.95 | 0.06 | 0.97 | 0.02 | 1.05 | 0.04 | 1.07 | 0.09 | 0.97 | 0.04 |
| | 5 | 1.00 | 0.07 | 1.67 | 0.56 | 0.94 | 0.03 | 1.00 | 0.03 | 0.82 | 0.03 | 1.00 | 0.12 | 0.94 | 0.08 | - | - |
| | 6 | 1.00 | 0.08 | 0.88 | 0.06 | - | - | 1.03 | 0.16 | - | - | - | - | - | - | - | - |
| Fumarate | 1 | 1.00 | 0.02 | 1.25 | 0.03 | 1.10 | 0.04 | 1.36 | 0.13 | 1.42 | 0.04 | - | - | - | - | - | - |
| | 3 | 1.00 | 0.05 | 0.99 | 0.00 | 0.91 | 0.05 | 1.20 | 0.08 | 0.97 | 0.05 | - | - | - | - | 1.10 | 0.03 |
| | 4 | 1.00 | 0.15 | 1.01 | 0.07 | 0.95 | 0.03 | 1.25 | 0.06 | 1.11 | 0.03 | 1.37 | 0.04 | 1.43 | 0.04 | 1.25 | 0.06 |
| | 5 | 1.00 | 0.42 | 0.76 | 0.15 | 0.27 | 0.04 | 0.43 | 0.08 | 1.10 | 0.07 | 0.35 | 0.02 | 0.35 | 0.11 | - | - |
| | 6 | 1.00 | 0.04 | 0.82 | 0.07 | - | - | 1.20 | 0.03 | - | - | - | - | - | - | - | - |
| Malate | 1 | 1.00 | 0.03 | 1.19 | 0.03 | 1.36 | 0.14 | 1.04 | 0.01 | 0.96 | 0.06 | - | - | - | - | - | - |
| | 3 | 1.00 | 0.07 | 0.89 | 0.02 | 0.79 | 0.07 | 1.00 | 0.07 | 0.82 | 0.05 | - | - | - | - | 0.91 | 0.02 |
| | 4 | 1.00 | 0.02 | 1.10 | 0.05 | 0.94 | 0.12 | 1.20 | 0.06 | 1.09 | 0.03 | 1.05 | 0.02 | 1.17 | 0.04 | 1.20 | 0.02 |
| | 5 | 1.00 | 0.11 | 1.53 | 0.31 | 1.57 | 0.09 | 1.52 | 0.10 | 1.00 | 0.05 | 1.47 | 0.17 | 1.09 | 0.10 | - | - |
| | 6 | 1.00 | 0.06 | 0.95 | 0.06 | - | - | 1.65 | 0.12 | - | - | - | - | - | - | - | - |
| 6-Phosphogluconate | 1 | 1.00 | 0.07 | 1.30 | 0.15 | 0.99 | 0.09 | 0.90 | 0.08 | 1.05 | 0.04 | - | - | - | - | - | - |
| | 3 | 1.00 | 0.09 | 0.76 | 0.03 | 0.58 | 0.02 | 1.06 | 0.25 | 0.67 | 0.05 | - | - | - | - | 0.67 | 0.07 |
| | 4 | - | - | - | - | - | - | - | - | - | - | - | - | - | - | - | - |
| | 5 | - | - | - | - | - | - | - | - | - | - | - | - | - | - | - | - |
| | 6 | 1.00 | 0.12 | 1.06 | 0.16 | - | - | 0.74 | 0.06 | - | - | - | - | - | - | - | - |
| Pentose phosphate | 1 | 1.00 | 0.13 | 1.23 | 0.12 | 0.84 | 0.11 | 1.01 | 0.12 | 0.89 | 0.10 | - | - | - | - | - | - |
| | 3 | 1.00 | 0.32 | 0.83 | 0.15 | 1.10 | 0.14 | 0.86 | 0.12 | 0.65 | 0.09 | - | - | - | - | 0.88 | 0.14 |
| | 4 | 1.00 | 0.29 | 0.73 | 0.08 | 0.89 | 0.12 | 0.56 | 0.13 | 0.64 | 0.00 | 0.75 | 0.06 | 0.75 | 0.08 | 0.61 | 0.02 |
| | 5 | 1.00 | 0.18 | 1.75 | 0.22 | 1.33 | 0.05 | 2.34 | 0.54 | 1.78 | 0.02 | 2.37 | 0.55 | 1.75 | 0.21 | - | - |
| | 6 | 1.00 | 0.07 | 1.01 | 0.10 | - | - | 0.83 | 0.10 | - | - | - | - | - | - | - | - |
| Sedoheptulose-7-phosphate | 1 | 1.00 | 0.04 | 1.59 | 0.09 | 0.77 | 0.03 | 0.86 | 0.06 | 0.93 | 0.05 | - | - | - | - | - | - |
| | 3 | 1.00 | 0.21 | 0.34 | 0.03 | 0.46 | 0.09 | 0.67 | 0.12 | 0.42 | 0.01 | - | - | - | - | 0.62 | 0.11 |
| | 4 | 1.00 | 0.11 | 1.33 | 0.07 | 1.04 | 0.07 | 1.18 | 0.05 | 0.85 | 0.08 | 1.05 | 0.06 | 0.77 | 0.07 | 1.34 | 0.03 |
| | 5 | 1.00 | 0.26 | 2.68 | 0.58 | 1.66 | 0.30 | 6.09 | 1.52 | 2.99 | 0.38 | 5.53 | 1.35 | 2.48 | 0.46 | - | - |
| | 6 | 1.00 | 0.15 | 0.91 | 0.13 | - | - | 1.11 | 0.17 | - | - | - | - | - | - | - | - |
| NADP+ | 1 | - | - | - | - | - | - | - | - | - | - | - | - | - | - | - | - |
| | 3 | - | - | - | - | - | - | - | - | - | - | - | - | - | - | - | - |
| | 4 | 1.00 | 0.12 | 1.16 | 0.06 | 1.27 | 0.14 | 1.08 | 0.12 | 1.41 | 0.08 | 1.15 | 0.07 | 1.08 | 0.23 | 0.93 | 0.13 |
| | 5 | 1.00 | 0.15 | 0.65 | 0.10 | 1.40 | 0.51 | 1.19 | 0.25 | 0.52 | 0.31 | 0.72 | 0.42 | 0.59 | 0.39 | - | - |
| | 6 | 1.00 | 0.22 | 0.93 | 0.08 | - | - | 0.47 | 0.13 | - | - | - | - | - | - | - | - |
| NADH | 1 | - | - | - | - | - | - | - | - | - | - | - | - | - | - | - | - |
| | 3 | 1.00 | 0.07 | 1.28 | 0.02 | 0.90 | 0.06 | 1.33 | 0.12 | 0.82 | 0.04 | - | - | - | - | 1.37 | 0.13 |
| | 4 | - | - | - | - | - | - | - | - | - | - | - | - | - | - | - | - |
| | 5 | 1.00 | 0.25 | 0.66 | 0.23 | 0.76 | 0.03 | 0.38 | 0.06 | 0.79 | 0.05 | 0.86 | 0.25 | 1.22 | 0.25 | - | - |
| | 6 | - | - | - | - | - | - | - | - | - | - | - | - | - | - | - | - |
| NAD+ | 1 | 1.00 | 0.05 | 0.88 | 0.09 | 0.40 | 0.05 | 0.50 | 0.05 | 0.52 | 0.05 | - | - | - | - | - | - |
| | 3 | 1.00 | 0.16 | 1.12 | 0.09 | 0.90 | 0.07 | 0.94 | 0.03 | 0.89 | 0.16 | - | - | - | - | 1.68 | 0.10 |
| | 4 | - | - | - | - | - | - | - | - | - | - | - | - | - | - | - | - |
| | 5 | 1.00 | 0.03 | 0.61 | 0.04 | 0.70 | 0.07 | 0.58 | 0.08 | 0.73 | 0.02 | 0.76 | 0.09 | 1.01 | 0.10 | - | - |
| | 6 | 1.00 | 0.07 | 1.02 | 0.10 | - | - | 0.53 | 0.14 | - | - | - | - | - | - | - | - |
| Glutathione | 1 | 1.00 | 0.10 | 0.90 | 0.05 | 0.40 | 0.08 | 1.84 | 0.23 | 1.73 | 0.07 | - | - | - | - | - | - |
| | 3 | - | - | - | - | - | - | - | - | - | - | - | - | - | - | - | - |
| | 4 | 1.00 | 0.23 | 0.63 | 0.19 | 1.20 | 0.36 | 1.33 | 0.38 | 0.49 | 0.06 | 1.51 | 0.32 | 1.02 | 0.14 | 0.81 | 0.11 |
| | 5 | 1.00 | 0.17 | 2.93 | 0.80 | 1.84 | 0.14 | 0.84 | 0.32 | 2.11 | 0.33 | 1.52 | 0.21 | 3.61 | 0.78 | - | - |
| | 6 | - | - | - | - | - | - | - | - | - | - | - | - | - | - | - | - |
| Glutathione disulfide | 1 | - | - | - | - | - | - | - | - | - | - | - | - | - | - | - | - |
| | 3 | 1.00 | 0.26 | 1.19 | 0.06 | 0.89 | 0.02 | 1.16 | 0.02 | 0.65 | 0.06 | - | - | - | - | 1.24 | 0.10 |
| | 4 | 1.00 | 0.29 | 0.96 | 0.17 | 0.90 | 0.01 | 0.78 | 0.06 | 0.94 | 0.13 | 1.13 | 0.08 | 1.28 | 0.15 | 0.68 | 0.07 |
| | 5 | 1.00 | 0.20 | 1.36 | 0.19 | 1.07 | 0.00 | 0.61 | 0.21 | 1.34 | 0.25 | 0.89 | 0.06 | 1.87 | 0.08 | - | - |
| | 6 | 1.00 | 0.08 | 0.96 | 0.11 | - | - | 0.58 | 0.17 | - | - | - | - | - | - | - | - |
| ATP | 1 | 1.00 | 0.04 | 0.90 | 0.09 | 1.03 | 0.02 | 0.35 | 0.04 | 0.44 | 0.01 | - | - | - | - | - | - |
| | 3 | 1.00 | 0.16 | 1.09 | 0.25 | 1.01 | 0.07 | 1.08 | 0.15 | 1.04 | 0.08 | - | - | - | - | 1.55 | 0.05 |
| | 4 | 1.00 | 0.32 | 0.77 | 0.10 | 0.81 | 0.07 | 0.49 | 0.12 | 0.98 | 0.07 | 0.59 | 0.03 | 0.63 | 0.17 | 0.43 | 0.02 |
| | 5 | 1.00 | 0.11 | 0.67 | 0.06 | 0.72 | 0.04 | 0.58 | 0.08 | 0.77 | 0.02 | 0.78 | 0.09 | 0.86 | 0.05 | - | - |
| | 6 | 1.00 | 0.19 | 0.80 | 0.18 | - | - | 0.24 | 0.09 | - | - | - | - | - | - | - | - |
| ADP | 1 | 1.00 | 0.02 | 1.08 | 0.07 | 1.50 | 0.20 | 1.26 | 0.13 | 1.07 | 0.15 | - | - | - | - | - | - |
| | 3 | 1.00 | 0.24 | 0.71 | 0.11 | 0.76 | 0.12 | 0.77 | 0.07 | 0.68 | 0.05 | - | - | - | - | 0.66 | 0.07 |
| | 4 | 1.00 | 0.13 | 1.13 | 0.12 | 0.97 | 0.06 | 0.96 | 0.05 | 1.52 | 0.10 | 1.40 | 0.03 | 1.30 | 0.16 | 0.87 | 0.00 |
| | 5 | 1.00 | 0.02 | 0.70 | 0.02 | 0.79 | 0.03 | 0.75 | 0.12 | 1.18 | 0.09 | 1.16 | 0.05 | 1.19 | 0.14 | - | - |
| | 6 | 1.00 | 0.04 | 0.80 | 0.04 | - | - | 0.61 | 0.12 | - | - | - | - | - | - | - | - |
| AMP | 1 | 1.00 | 0.07 | 1.04 | 0.09 | 0.86 | 0.12 | 0.75 | 0.10 | 0.57 | 0.08 | - | - | - | - | - | - |
| | 3 | 1.00 | 0.09 | 0.86 | 0.07 | 1.10 | 0.16 | 1.07 | 0.08 | 1.16 | 0.07 | - | - | - | - | 0.93 | 0.03 |
| | 4 | 1.00 | 0.14 | 1.39 | 0.34 | 1.30 | 0.21 | 1.29 | 0.10 | 1.42 | 0.21 | 1.31 | 0.08 | 0.97 | 0.05 | 2.05 | 0.34 |
| | 5 | 1.00 | 0.09 | 1.62 | 0.37 | 0.92 | 0.01 | 1.33 | 0.20 | 1.96 | 0.16 | 1.94 | 0.05 | 2.12 | 0.15 | - | - |
| | 6 | 1.00 | 0.13 | 0.87 | 0.18 | - | - | 1.83 | 0.29 | - | - | - | - | - | - | - | - |
| UTP | 1 | 1.00 | 0.02 | 1.18 | 0.07 | 1.05 | 0.02 | 0.40 | 0.05 | 0.50 | 0.02 | - | - | - | - | - | - |
| | 3 | 1.00 | 0.44 | 1.91 | 0.09 | 1.34 | 0.02 | 0.78 | 0.42 | 1.17 | 0.21 | - | - | - | - | 2.06 | 0.04 |
| | 4 | 1.00 | 0.31 | 0.81 | 0.09 | 0.73 | 0.08 | 0.52 | 0.12 | 1.03 | 0.04 | 0.59 | 0.01 | 0.64 | 0.18 | 0.51 | 0.02 |
| | 5 | 1.00 | 0.14 | 1.03 | 0.06 | 0.71 | 0.07 | 1.07 | 0.20 | 1.48 | 0.03 | 1.53 | 0.11 | 1.20 | 0.05 | - | - |
| | 6 | 1.00 | 0.19 | 0.75 | 0.15 | - | - | 0.61 | 0.21 | - | - | - | - | - | - | - | - |
| UDP | 1 | 1.00 | 0.02 | 1.58 | 0.16 | 1.87 | 0.26 | 1.35 | 0.12 | 1.29 | 0.18 | - | - | - | - | - | - |
| | 3 | 1.00 | 0.20 | 0.40 | 0.02 | 0.44 | 0.08 | 0.72 | 0.18 | 0.57 | 0.12 | - | - | - | - | 0.33 | 0.02 |
| | 4 | 1.00 | 0.15 | 1.18 | 0.17 | 1.01 | 0.12 | 1.10 | 0.06 | 1.43 | 0.14 | 1.31 | 0.12 | 1.20 | 0.04 | 1.22 | 0.11 |
| | 5 | 1.00 | 0.14 | 1.31 | 0.29 | 0.97 | 0.02 | 1.76 | 0.30 | 1.99 | 0.03 | 1.84 | 0.08 | 1.63 | 0.03 | - | - |
| | 6 | 1.00 | 0.05 | 0.85 | 0.06 | - | - | 1.38 | 0.22 | - | - | - | - | - | - | - | - |
| UMP | 1 | 1.00 | 0.01 | 1.82 | 0.34 | 3.08 | 0.35 | 2.39 | 0.57 | 2.22 | 0.36 | - | - | - | - | - | - |
| | 3 | 1.00 | 0.07 | 0.82 | 0.05 | 1.04 | 0.14 | 0.84 | 0.06 | 1.20 | 0.20 | - | - | - | - | 0.61 | 0.09 |
| | 4 | 1.00 | 0.35 | 1.31 | 0.40 | 0.91 | 0.18 | 0.79 | 0.09 | 1.82 | 0.27 | 1.00 | 0.16 | 1.02 | 0.22 | 1.28 | 0.12 |
| | 5 | 1.00 | 0.14 | 1.11 | 0.14 | 0.97 | 0.05 | 1.32 | 0.15 | 2.05 | 0.22 | 1.90 | 0.12 | 1.87 | 0.27 | - | - |
| | 6 | 1.00 | 0.06 | 0.69 | 0.06 | - | - | 1.93 | 0.39 | - | - | - | - | - | - | - | - |

Values represent the mean and standard error (n=3).

**Supplementary Table 3.** Amino Acid Sequences of ScFvs

***CAR-T Cell ScFv Amino Acid Sequence***

| ScFv | Region | Sequence |
|---|---|---|
| FMC63-based anti-CD19 | $V_H$ | EVKLQESGPGLVAPSQSLSVTCTVSGVSLPDYGVSWIRQPPRKGLEWLGVIWGSETTYYNSALKSRLTIIKDNSKSQVFLKMNSLQTDDTAIYYCAKHYYYGGSYAMDYWGQGTSVTVSS |
| Leu16 anti-CD20 | $V_H$ | EVQLQQSGAELVKPGASVKMSCKASGYTFTSYNMHWVKQTPGQGLEWIGAIYPGNGDTSYNQKFKGKATLTADKSSSTAYMQLSSLTSEDSADYYCARSNYYGSSYWFFDVWGAGTTVTVSS |
| Rituximab anti-CD20 | $V_H$ | QVQLQQPGAELVKPGASVKMSCKASGYTFTSYNMHWVKQTPGRGLEWIGAIYPGNGDTSYNQKFKGKATLTADKSSSTAYMQLSSLTSEDSAVYYCARSTYYGGDWYFNVWGAGTTVTVSS |
| RFR-LCDR anti-CD20 | $V_H$ | QVQLQQPGAELVKPGASVKMSCKASGYTFTSYNMHWVKQTPGRGLEWIGAIYPGNGDTSYNQKFKGKATLTADKSSSTAYMQLSSLTSEDSAVYYCARSNYYGSSYWFFDVWGAGTTVTVSS |
| 14g2a-based anti-GD2 | $V_H$ | EVQLLQSGPELEKPGASVMISCKASGSSFTGYNMNWVRQNIGKSLEWIGAIDPYYGGTSYNQKFKGRATLTVDKSSSTAYMHLKSLTSEDSAVYYCVSGMEYWGQGTSVTVSS |
| FMC63-based anti-CD19 | $V_L$ | DIQMTQTTSSLSASLGDRVTISCRASQDISKYLNWYQQKPDGTVKLLIYHTSRLHSGVPSRFSGSGSGTDYSLTISNLEQEDIATYFCQQGNTLPYTFGGGTKLEIT |
| Leu16 anti-CD20 | $V_L$ | DIVLTQSPAILSASPGEKVTMTCRASSSVNYMDWYQKKPGSSPKPWIYATSNLASGVPARFSGSGSGTSYSLTISRVEAEDAATYYCQQWSFNPPTFGGGTKLEIK |
| Rituximab anti-CD20 | $V_L$ | QIVLSQSPAILSASPGEKVTMTCRASSSVSYIHWFQQKPGSSPKPWIYATSNLASGVPVRFSGSGSGTSYSLTISRVEAEDAATYYCQQWTSNPPTFGGGTKLEIK |
| RFR-LCDR anti-CD20 | $V_L$ | QIVLSQSPAILSASPGEKVTMTCRASSSVNYMDWFQQKPGSSPKPWIYATSNLASGVPVRFSGSGSGTSYSLTISRVEAEDAATYYCQQWSFNPPTFGGGTKLEIK |
| 14g2a-based anti-GD2 | $V_L$ | DVVMTQTPLSLPVSLGDQASISCRSSQSLVHRNGNTYLHWYLQKPGQSPKLLIHKVSNRFSGVPDRFSGSGSGTDFTLKISRVEAEDLGVYFCSQSTHVPPLTFGAGTKLELKRA |
| FMC63-based anti-CD19 | Whole | DIQMTQTTSSLSASLGDRVTISCRASQDISKYLNWYQQKPDGTVKLLIYHTSRLHSGVPSRFSGSGSGTDYSLTISNLEQEDIATYFCQQGNTLPYTFGGGTKLEITGSTSGSGKPGSGEGSTKGEVKLQESGPGLVAPSQSLSVTCTVSGVSLPDYGVSWIRQPPRKGLEWLGVIWGSETTYYNSALKSRLTIIKDNSKSQVFLKMNSLQTDDTAIYYCAKHYYYGGSYAMDYWGQGTSVTVSS |
| Leu16 anti-CD20 | Whole | DIVLTQSPAILSASPGEKVTMTCRASSSVNYMDWYQKKPGSSPKPWIYATSNLASGVPARFSGSGSGTSYSLTISRVEAEDAATYYCQQWSFNPPTFGGGTKLEIKGSTSGGGSGGGSGGGGSSEVQLQQSGAELVKPGASVKMSCKASGYTFTSYNMHWVKQTPGQGLEWIGAIYPGNGDTSYNQKFKGKATLTADKSSSTAYMQLSSLTSEDSADYYCARSNYYGSSYWFFDVWGAGTTVTVSS |
| Rituximab anti-CD20 | Whole | QIVLSQSPAILSASPGEKVTMTCRASSSVSYIHWFQQKPGSSPKPWIYATSNLASGVPVRFSGSGSGTSYSLTISRVEAEDAATYYCQQWTSNPPTFGGGTKLEIKGSTSGGGSGGGSGGGGSSQVQLQQPGAELVKPGASVKMSCKASGYTFTSYNMHWVKQTPGRGLEWIGAIYPGNGDTSYNQKFKGKATLTADKSSSTAYMQLSSLTSEDSAVYYCARSTYYGGDWYFNVWGAGTTVTVSS |
| RFR-LCDR anti-CD20 | Whole | QIVLSQSPAILSASPGEKVTMTCRASSSVNYMDWFQQKPGSSPKPWIYATSNLASGVPVRFSGSGSGTSYSLTISRVEAEDAATYYCQQWSFNPPTFGGGTKLEIKGSTSGGGSGGGSGGGGSSQVQLQQPGAELVKPGASVKMSCKASGYTFTSYNMHWVKQTPGRGLEWIGAIYPGNGDTSYNQKFKGKATLTADKSSSTAYMQLSSLTSEDSAVYYCARSNYYGSSYWFFDVWGAGTTVTVSS |
| 14g2a-based anti-GD2 | Whole | DVVMTQTPLSLPVSLGDQASISCRSSQSLVHRNGNTYLHWYLQKPGQSPKLLIHKVSNRFSGVPDRFSGSGSGTDFTLKISRVEAEDLGVYFCSQSTHVPPLTFGAGTKLELKRAGSTSGSGKPGSGEGSTKGEVQLLQSGPELEKPGASVMISCKASGSSFTGYNMNWVRQNIGKSLEWIGAIDPYYGGTSYNQKFKGRATLTVDKSSSTAYMHLKSLTSEDSAVYYCVSGMEYWGQGTSVTVSS |

**Supplementary Table 4.** Percent Identity Matrix for ScFvs

| ***CAR-T Cell ScFv Sequence Homology Matrix*** | | | | | | |
|---|---|---|---|---|---|---|
| **ScFv** | **Region** | **CD19** | **Leu16** | **Rituximab** | **RFR-LCDR** | **GD2** |
| FMC63-based anti-CD19 | $V_H$ | 100.00 | 45.00 | 42.50 | 43.33 | 45.54 |
| Leu16 anti-CD20 | $V_H$ | 45.00 | 100.00 | 92.56 | 96.72 | 72.57 |
| Rituximab anti-CD20 | $V_H$ | 42.50 | 92.56 | 100.00 | 95.87 | 71.68 |
| RFR-LCDR anti-CD20 | $V_H$ | 43.33 | 96.72 | 95.87 | 100.00 | 71.68 |
| 14g2a-based anti-GD2 | $V_H$ | 45.54 | 72.57 | 71.68 | 71.68 | 100.00 |
| FMC63-based anti-CD19 | $V_L$ | 100.00 | 62.26 | 60.38 | 60.38 | 57.94 |
| Leu16 anti-CD20 | $V_L$ | 62.26 | 100.00 | 90.57 | 95.28 | 56.60 |
| Rituximab anti-CD20 | $V_L$ | 60.38 | 90.57 | 100.00 | 95.28 | 56.60 |
| RFR-LCDR anti-CD20 | $V_L$ | 60.38 | 95.28 | 95.28 | 100.00 | 54.72 |
| 14g2a-based anti-GD2 | $V_L$ | 57.94 | 56.60 | 56.60 | 54.72 | 100.00 |
| FMC63-based anti-CD19 | Mean of $V_H$ & $V_L$ | 100.00 | 53.63 | 51.44 | 51.86 | 51.74 |
| Leu16 anti-CD20 | Mean of $V_H$ & $V_L$ | 53.63 | 100.00 | 91.57 | 96.00 | 64.59 |
| Rituximab anti-CD20 | Mean of $V_H$ & $V_L$ | 51.44 | 91.57 | 100.00 | 95.58 | 64.14 |
| RFR-LCDR anti-CD20 | Mean of $V_H$ & $V_L$ | 51.86 | 96.00 | 95.58 | 100.00 | 63.20 |
| 14g2a-based anti-GD2 | Mean of $V_H$ & $V_L$ | 51.74 | 64.59 | 64.14 | 63.20 | 100.00 |
| FMC63-based anti-CD19 | Whole | 100.00 | 52.46 | 50.41 | 50.82 | 55.27 |
| Leu16 anti-CD20 | Whole | 52.46 | 100.00 | 92.24 | 96.34 | 63.29 |
| Rituximab anti-CD20 | Whole | 50.41 | 92.24 | 100.00 | 95.92 | 62.87 |
| RFR-LCDR anti-CD20 | Whole | 50.82 | 96.34 | 95.92 | 100.00 | 62.03 |
| 14g2a-based anti-GD2 | Whole | 55.27 | 63.29 | 62.87 | 62.03 | 100.00 |

**Supplementary Table 5.** CAR-T Cell Uptake and Secretion Rates (nmol/million cells/hr)

| *CAR-T Cell Uptake and Secretion Rates (nmol/million cells/hr)* | | | | | | | | | | | | | | | | | |
|---|---|---|---|---|---|---|---|---|---|---|---|---|---|---|---|---|---|
| **Metabolite** | **Donor** | **EGFRt** | **s.e.m.** | **CD19** | **s.e.m.** | **Leu16** | **s.e.m.** | **Rituximab** | **s.e.m.** | **RFR-LCDR** | **s.e.m.** | **Rituximab.AA** | **s.e.m.** | **RFR-LCDR.AA** | **s.e.m.** | **GD2** | **s.e.m.** |
| Alanine (secretion) | 1 | 0.68 | 0.06 | 0.67 | 0.06 | 0.71 | 0.06 | 0.80 | 0.06 | 0.57 | 0.05 | - | - | - | - | - | - |
| | 2 | 0.24 | 0.01 | 0.39 | 0.06 | 0.36 | 0.02 | 4.44 | 0.15 | 1.09 | 0.15 | 4.54 | 0.15 | 1.64 | 0.09 | - | - |
| | 3 | 0.18 | 0.01 | 0.47 | 0.06 | 0.26 | 0.01 | 0.93 | 0.03 | 1.10 | 0.04 | - | - | - | - | 1.49 | 0.03 |
| | 4 | 0.38 | 0.02 | 0.46 | 0.06 | 0.39 | 0.02 | 1.19 | 0.05 | 0.49 | 0.04 | 0.73 | 0.05 | 0.43 | 0.01 | 1.45 | 0.10 |
| Serine | 1 | 1.66 | 0.15 | 2.26 | 0.21 | 1.98 | 0.30 | 2.82 | 0.12 | 2.86 | 0.17 | - | - | - | - | - | - |
| | 2 | 1.57 | 0.08 | 2.32 | 0.24 | 1.80 | 0.13 | 4.47 | 0.18 | 1.85 | 0.25 | 4.62 | 0.36 | 2.81 | 0.15 | - | - |
| | 3 | 0.57 | 0.12 | 1.87 | 0.31 | 1.47 | 0.01 | 1.48 | 0.14 | 1.30 | 0.10 | - | - | - | - | 1.33 | 0.24 |
| | 4 | 1.20 | 0.01 | 1.51 | 0.17 | 1.62 | 0.10 | 2.48 | 0.12 | 1.78 | 0.07 | 2.04 | 0.06 | 1.40 | 0.16 | 2.78 | 0.13 |
| Glycine | 1 | 0.17 | 0.09 | 0.32 | 0.16 | 0.32 | 0.09 | 0.14 | 0.10 | 0.26 | 0.06 | - | - | - | - | - | - |
| | 2 | 0.09 | 0.04 | 0.07 | 0.05 | 0.03 | 0.03 | 0.00 | 0.00 | 0.25 | 0.08 | 0.38 | 0.04 | 0.00 | 0.00 | - | - |
| | 3 | 0.23 | 0.11 | 0.29 | 0.14 | 0.39 | 0.08 | 0.00 | 0.00 | 0.31 | 0.11 | - | - | - | - | 0.09 | 0.09 |
| | 4 | 0.00 | 0.00 | 0.12 | 0.10 | 0.02 | 0.02 | 0.05 | 0.03 | 0.00 | 0.00 | 0.00 | 0.00 | 0.00 | 0.00 | 0.30 | 0.04 |
| Methionine | 1 | 0.05 | 0.03 | 0.44 | 0.03 | 0.30 | 0.08 | 0.69 | 0.04 | 0.71 | 0.03 | - | - | - | - | - | - |
| | 2 | 0.45 | 0.02 | 0.60 | 0.12 | 0.48 | 0.02 | 1.27 | 0.03 | 0.54 | 0.09 | 1.50 | 0.07 | 0.86 | 0.02 | - | - |
| | 3 | 0.31 | 0.02 | 0.62 | 0.03 | 0.49 | 0.02 | 0.47 | 0.00 | 0.38 | 0.02 | - | - | - | - | 0.49 | 0.06 |
| | 4 | 0.37 | 0.02 | 0.54 | 0.06 | 0.45 | 0.05 | 0.78 | 0.06 | 0.48 | 0.02 | 0.59 | 0.02 | 0.37 | 0.04 | 0.85 | 0.07 |
| Cystine | 1 | 1.08 | 0.13 | 1.06 | 0.06 | 0.65 | 0.27 | 0.79 | 0.16 | 0.94 | 0.09 | - | - | - | - | - | - |
| | 2 | 0.82 | 0.07 | 0.77 | 0.11 | 1.09 | 0.08 | 1.30 | 0.13 | 0.78 | 0.11 | 1.77 | 0.19 | 1.00 | 0.13 | - | - |
| | 3 | 0.43 | 0.01 | 0.66 | 0.02 | 0.72 | 0.14 | 0.65 | 0.06 | 0.91 | 0.01 | - | - | - | - | 0.67 | 0.06 |
| | 4 | 0.51 | 0.05 | 0.54 | 0.11 | 0.63 | 0.05 | 0.62 | 0.11 | 0.16 | 0.03 | 0.49 | 0.09 | 0.31 | 0.06 | 1.21 | 0.05 |
| Leucine | 1 | 0.41 | 0.05 | 1.03 | 0.10 | 0.08 | 0.04 | 1.28 | 0.10 | 1.32 | 0.11 | - | - | - | - | - | - |
| | 2 | 1.57 | 0.12 | 1.96 | 0.19 | 1.79 | 0.22 | 3.55 | 0.22 | 1.71 | 0.17 | 4.35 | 0.28 | 2.57 | 0.14 | - | - |
| | 3 | 0.89 | 0.08 | 1.47 | 0.14 | 1.34 | 0.15 | 1.26 | 0.13 | 0.82 | 0.04 | - | - | - | - | 1.14 | 0.02 |
| | 4 | 0.95 | 0.12 | 1.17 | 0.10 | 1.17 | 0.22 | 1.81 | 0.13 | 1.12 | 0.07 | 1.22 | 0.07 | 0.59 | 0.04 | 2.16 | 0.14 |
| Isoleucine | 1 | 0.80 | 0.37 | 1.27 | 0.12 | 0.56 | 0.13 | 1.24 | 0.07 | 1.42 | 0.11 | - | - | - | - | - | - |
| | 2 | 1.60 | 0.02 | 2.59 | 0.27 | 1.70 | 0.16 | 3.43 | 0.12 | 1.70 | 0.35 | 4.45 | 0.30 | 2.61 | 0.07 | - | - |
| | 3 | 1.09 | 0.10 | 2.14 | 0.21 | 1.64 | 0.08 | 1.44 | 0.18 | 0.97 | 0.03 | - | - | - | - | 1.33 | 0.03 |
| | 4 | 1.06 | 0.05 | 1.42 | 0.22 | 1.16 | 0.20 | 1.81 | 0.14 | 0.69 | 0.02 | 1.03 | 0.05 | 0.44 | 0.10 | 2.21 | 0.13 |
| Valine | 1 | 1.15 | 0.10 | 1.25 | 0.12 | 1.10 | 0.10 | 1.19 | 0.07 | 1.16 | 0.06 | - | - | - | - | - | - |
| | 2 | 0.79 | 0.06 | 1.27 | 0.08 | 0.81 | 0.05 | 2.01 | 0.14 | 0.92 | 0.18 | 2.33 | 0.16 | 1.37 | 0.10 | - | - |
| | 3 | 0.38 | 0.00 | 1.02 | 0.10 | 0.93 | 0.09 | 0.87 | 0.05 | 0.66 | 0.11 | - | - | - | - | 0.81 | 0.04 |
| | 4 | 0.69 | 0.01 | 0.96 | 0.11 | 0.82 | 0.09 | 1.28 | 0.09 | 0.83 | 0.03 | 0.93 | 0.02 | 0.64 | 0.04 | 1.56 | 0.06 |
| Phenylalanine | 1 | 1.06 | 0.16 | 0.96 | 0.07 | 0.71 | 0.01 | 0.79 | 0.05 | 0.71 | 0.04 | - | - | - | - | - | - |
| | 2 | 0.41 | 0.06 | 0.62 | 0.08 | 0.36 | 0.04 | 1.01 | 0.06 | 0.44 | 0.08 | 1.23 | 0.09 | 0.69 | 0.02 | - | - |
| | 3 | 0.17 | 0.01 | 0.46 | 0.06 | 0.34 | 0.06 | 0.29 | 0.02 | 0.31 | 0.07 | - | - | - | - | 0.25 | 0.03 |
| | 4 | 0.25 | 0.02 | 0.45 | 0.04 | 0.52 | 0.02 | 0.65 | 0.02 | 0.46 | 0.03 | 0.51 | 0.04 | 0.31 | 0.04 | 0.82 | 0.02 |
| Tyrosine | 1 | 0.84 | 0.07 | 0.76 | 0.09 | 0.66 | 0.06 | 0.98 | 0.08 | 0.90 | 0.05 | - | - | - | - | - | - |
| | 2 | 0.43 | 0.07 | 0.69 | 0.10 | 0.44 | 0.04 | 0.94 | 0.05 | 0.42 | 0.07 | 1.24 | 0.09 | 0.66 | 0.04 | - | - |
| | 3 | 0.13 | 0.11 | 0.13 | 0.01 | 0.45 | 0.06 | 0.54 | 0.02 | 0.69 | 0.02 | - | - | - | - | 0.32 | 0.05 |
| | 4 | 0.55 | 0.03 | 0.52 | 0.02 | 0.59 | 0.03 | 0.93 | 0.06 | 0.58 | 0.03 | 0.63 | 0.02 | 0.43 | 0.05 | 1.04 | 0.06 |
| Tryptophan | 1 | 0.00 | 0.00 | 0.30 | 0.08 | 0.00 | 0.00 | 0.24 | 0.00 | 0.30 | 0.01 | - | - | - | - | - | - |
| | 2 | 0.12 | 0.01 | 0.16 | 0.02 | 0.13 | 0.02 | 0.45 | 0.01 | 0.17 | 0.03 | 0.49 | 0.02 | 0.28 | 0.01 | - | - |
| | 3 | 0.12 | 0.02 | 0.20 | 0.02 | 0.19 | 0.01 | 0.18 | 0.02 | 0.12 | 0.02 | - | - | - | - | 0.15 | 0.02 |
| | 4 | 0.09 | 0.02 | 0.15 | 0.01 | 0.14 | 0.02 | 0.24 | 0.01 | 0.13 | 0.01 | 0.16 | 0.01 | 0.09 | 0.01 | 0.30 | 0.05 |
| Glutamine | 1 | 32.55 | 3.47 | 32.88 | 1.93 | 27.63 | 0.90 | 29.25 | 1.92 | 25.30 | 1.91 | - | - | - | - | - | - |
| | 2 | 10.38 | 0.70 | 15.14 | 1.31 | 13.51 | 1.46 | 33.18 | 1.77 | 13.00 | 1.94 | 37.71 | 1.03 | 21.20 | 0.06 | - | - |
| | 3 | 20.31 | 0.73 | 22.28 | 0.18 | 30.62 | 0.46 | 24.77 | 0.16 | 16.26 | 0.77 | - | - | - | - | 28.28 | 2.68 |
| | 4 | 21.61 | 0.72 | 24.86 | 1.11 | 20.88 | 2.25 | 28.83 | 1.46 | 18.99 | 1.65 | 21.21 | 0.84 | 13.44 | 0.47 | 37.55 | 0.18 |
| Glutamate (secretion) | 1 | 0.59 | 0.10 | 0.44 | 0.13 | 0.59 | 0.12 | 0.86 | 0.08 | 0.68 | 0.08 | - | - | - | - | - | - |
| | 2 | 0.07 | 0.03 | 1.79 | 0.57 | 0.36 | 0.04 | 3.92 | 0.37 | 1.15 | 0.23 | 4.48 | 0.20 | 1.68 | 0.18 | - | - |
| | 3 | 0.27 | 0.01 | 1.18 | 0.01 | 0.00 | 0.00 | 3.81 | 0.08 | 1.76 | 0.17 | - | - | - | - | 3.17 | 0.19 |
| | 4 | 0.82 | 0.03 | 1.71 | 0.15 | 0.84 | 0.18 | 3.10 | 0.19 | 1.91 | 0.15 | 2.42 | 0.16 | 1.78 | 0.08 | 3.99 | 0.39 |
| Asparagine | 1 | 2.09 | 0.17 | 1.89 | 0.19 | 1.68 | 0.26 | 1.10 | 0.18 | 1.47 | 0.07 | - | - | - | - | - | - |
| | 2 | 1.83 | 0.38 | 2.55 | 0.11 | 1.38 | 0.09 | 0.19 | 0.07 | 1.12 | 0.17 | 1.28 | 0.29 | 1.43 | 0.13 | - | - |
| | 3 | 1.05 | 0.07 | 1.68 | 0.17 | 2.35 | 0.13 | 1.82 | 0.09 | 1.07 | 0.16 | - | - | - | - | 1.80 | 0.16 |
| | 4 | 1.31 | 0.19 | 1.33 | 0.19 | 1.26 | 0.18 | 1.44 | 0.04 | 0.81 | 0.05 | 0.76 | 0.02 | 0.57 | 0.03 | 1.80 | 0.14 |
| Aspartate | 1 | 1.08 | 0.08 | 1.07 | 0.12 | 0.77 | 0.05 | 0.76 | 0.07 | 0.58 | 0.03 | - | - | - | - | - | - |
| | 2 | 0.76 | 0.06 | 1.29 | 0.08 | 0.76 | 0.05 | 1.28 | 0.08 | 0.72 | 0.12 | 1.64 | 0.13 | 0.91 | 0.08 | - | - |
| | 3 | 0.56 | 0.02 | 0.79 | 0.04 | 0.97 | 0.05 | 0.34 | 0.07 | 0.24 | 0.08 | - | - | - | - | 0.59 | 0.01 |
| | 4 | 0.21 | 0.05 | 0.47 | 0.05 | 0.54 | 0.06 | 0.83 | 0.11 | 0.45 | 0.01 | 0.30 | 0.01 | 0.23 | 0.04 | 0.73 | 0.05 |
| Arginine | 1 | 7.46 | 0.67 | 4.31 | 0.29 | 4.65 | 1.09 | 4.16 | 0.70 | 2.96 | 0.22 | - | - | - | - | - | - |
| | 2 | 4.87 | 0.74 | 4.47 | 0.59 | 4.11 | 0.46 | 7.86 | 0.34 | 3.93 | 0.62 | 8.91 | 0.43 | 6.13 | 0.32 | - | - |
| | 3 | 2.16 | 0.17 | 1.93 | 0.06 | 3.26 | 0.36 | 2.66 | 0.13 | 1.79 | 0.26 | - | - | - | - | 2.79 | 0.35 |
| | 4 | 3.50 | 0.70 | 3.33 | 0.04 | 3.20 | 0.75 | 4.84 | 0.46 | 3.20 | 0.35 | 3.86 | 0.34 | 1.60 | 0.02 | 8.52 | 0.99 |
| Proline | 1 | 0.72 | 0.08 | 0.92 | 0.09 | 0.68 | 0.16 | 0.80 | 0.04 | 0.79 | 0.02 | - | - | - | - | - | - |
| | 2 | 0.61 | 0.03 | 1.00 | 0.11 | 0.64 | 0.05 | 1.66 | 0.06 | 0.71 | 0.14 | 2.16 | 0.12 | 1.15 | 0.07 | - | - |
| | 3 | 0.42 | 0.02 | 0.75 | 0.08 | 0.74 | 0.05 | 0.53 | 0.05 | 0.24 | 0.01 | - | - | - | - | 0.56 | 0.04 |
| | 4 | 0.62 | 0.01 | 0.73 | 0.05 | 0.63 | 0.11 | 0.88 | 0.10 | 0.55 | 0.04 | 0.64 | 0.04 | 0.37 | 0.07 | 1.23 | 0.06 |
| Histidine | 1 | 0.44 | 0.05 | 0.43 | 0.05 | 0.45 | 0.09 | 0.66 | 0.04 | 0.63 | 0.05 | - | - | - | - | - | - |
| | 2 | 0.51 | 0.05 | 0.72 | 0.11 | 0.40 | 0.08 | 0.70 | 0.05 | 0.25 | 0.02 | 0.94 | 0.04 | 0.62 | 0.02 | - | - |
| | 3 | 0.54 | 0.05 | 0.57 | 0.10 | 0.70 | 0.10 | 0.46 | 0.00 | 0.14 | 0.07 | - | - | - | - | 0.43 | 0.08 |
| | 4 | 0.65 | 0.16 | 0.02 | 0.01 | 0.93 | 0.05 | 1.94 | 0.05 | 1.27 | 0.03 | 1.45 | 0.15 | 1.46 | 0.18 | 1.78 | 0.16 |
| Threonine | 1 | 0.82 | 0.05 | 1.37 | 0.13 | 1.14 | 0.07 | 1.25 | 0.04 | 1.06 | 0.06 | - | - | - | - | - | - |
| | 2 | 0.70 | 0.06 | 1.11 | 0.04 | 0.84 | 0.13 | 1.90 | 0.00 | 0.82 | 0.14 | 2.20 | 0.12 | 1.43 | 0.07 | - | - |
| | 3 | 0.61 | 0.04 | 0.65 | 0.01 | 0.82 | 0.18 | 0.57 | 0.07 | 0.27 | 0.11 | - | - | - | - | 0.91 | 0.10 |
| | 4 | 0.39 | 0.04 | 0.64 | 0.05 | 0.70 | 0.06 | 1.30 | 0.07 | 0.81 | 0.03 | 0.90 | 0.04 | 0.49 | 0.05 | 1.09 | 0.13 |
| Lysine | 1 | 1.62 | 0.35 | 1.00 | 0.22 | 0.61 | 0.10 | 1.19 | 0.20 | 1.25 | 0.16 | - | - | - | - | - | - |
| | 2 | 0.92 | 0.11 | 0.93 | 0.22 | 1.04 | 0.07 | 2.43 | 0.13 | 0.90 | 0.15 | 2.71 | 0.05 | 1.56 | 0.02 | - | - |
| | 3 | 0.84 | 0.12 | 1.43 | 0.09 | 0.97 | 0.07 | 0.55 | 0.15 | 1.39 | 0.11 | - | - | - | - | 1.12 | 0.29 |
| | 4 | 0.58 | 0.06 | 0.61 | 0.13 | 0.52 | 0.09 | 1.00 | 0.04 | 0.31 | 0.02 | 0.67 | 0.02 | 0.37 | 0.07 | 1.18 | 0.10 |
| Hydroxyproline | 1 | 0.09 | 0.03 | 0.43 | 0.12 | 0.03 | 0.03 | 0.17 | 0.04 | 0.19 | 0.03 | - | - | - | - | - | - |
| | 2 | 0.59 | 0.02 | 0.86 | 0.03 | 0.70 | 0.08 | 0.79 | 0.11 | 0.54 | 0.05 | 1.67 | 0.10 | 0.85 | 0.08 | - | - |
| | 3 | 0.59 | 0.05 | 0.54 | 0.14 | 0.60 | 0.07 | 0.50 | 0.07 | 0.36 | 0.05 | - | - | - | - | 0.66 | 0.06 |
| | 4 | 0.54 | 0.03 | 0.45 | 0.09 | 0.57 | 0.06 | 0.82 | 0.07 | 0.37 | 0.04 | 0.33 | 0.06 | 0.19 | 0.02 | 0.93 | 0.04 |
| Myo-inositol | 1 | 0.05 | 0.21 | 0.23 | 0.03 | 0.25 | 0.09 | 0.23 | 0.04 | 0.45 | 0.03 | - | - | - | - | - | - |
| | 2 | 0.64 | 0.14 | 1.27 | 0.16 | 0.61 | 0.09 | 0.85 | 0.08 | 0.54 | 0.08 | 1.44 | 0.05 | 0.85 | 0.06 | - | - |
| | 3 | 0.46 | 0.05 | 0.56 | 0.04 | 0.51 | 0.01 | 0.47 | 0.02 | 0.22 | 0.01 | - | - | - | - | 0.45 | 0.02 |
| | 4 | 0.58 | 0.03 | 0.61 | 0.04 | 0.48 | 0.04 | 1.31 | 0.10 | 0.42 | 0.06 | 0.56 | 0.04 | 0.58 | 0.02 | 1.41 | 0.16 |
| Glucose | 1 | 74.35 | 7.91 | 87.18 | 7.72 | 67.54 | 6.79 | 94.38 | 5.56 | 83.92 | 2.82 | - | - | - | - | - | - |
| | 2 | 49.86 | 7.94 | 93.11 | 13.33 | 65.08 | 5.61 | 181.57 | 5.84 | 67.07 | 11.32 | 206.32 | 5.46 | 110.15 | 3.61 | - | - |
| | 3 | 80.06 | 1.19 | 87.38 | 3.36 | 87.10 | 1.88 | 123.66 | 3.82 | 78.85 | 3.06 | - | - | - | - | 128.75 | 0.58 |
| | 4 | 49.17 | 3.17 | 56.82 | 1.38 | 60.58 | 3.43 | 93.14 | 5.81 | 56.07 | 1.11 | 72.11 | 3.05 | 51.13 | 3.24 | 109.54 | 5.01 |
| Lactate | 1 | 81.37 | 2.38 | 94.01 | 3.61 | 91.40 | 6.36 | 117.20 | 7.83 | 112.60 | 7.85 | - | - | - | - | - | - |
| | 2 | 45.36 | 2.34 | 89.78 | 7.99 | 55.56 | 1.56 | 251.06 | 12.09 | 87.79 | 10.56 | 260.59 | 3.61 | 142.97 | 4.37 | - | - |
| | 3 | 68.21 | 2.72 | 102.59 | 2.84 | 89.12 | 1.67 | 110.45 | 2.70 | 110.46 | 2.13 | - | - | - | - | 117.70 | 1.45 |
| | 4 | 56.72 | 0.78 | 65.61 | 5.64 | 71.80 | 5.82 | 112.78 | 5.51 | 73.04 | 4.95 | 92.29 | 1.31 | 60.82 | 2.34 | 119.03 | 9.56 |
| Pyruvate | 1 | 1.41 | 0.15 | 1.33 | 0.20 | 1.73 | 0.05 | 1.54 | 0.18 | 1.19 | 0.11 | - | - | - | - | - | - |
| | 2 | 1.47 | 0.23 | 1.77 | 0.19 | 1.74 | 0.23 | 1.61 | 0.18 | 1.96 | 0.34 | 1.87 | 0.13 | 3.10 | 0.32 | - | - |
| | 3 | 1.11 | 0.05 | 1.98 | 0.15 | 1.71 | 0.05 | 2.03 | 0.08 | 2.20 | 0.10 | - | - | - | - | 1.88 | 0.07 |
| | 4 | 1.34 | 0.07 | 1.00 | 0.19 | 1.55 | 0.09 | 2.09 | 0.12 | 1.79 | 0.11 | 2.02 | 0.18 | 0.84 | 0.12 | 2.74 | 0.15 |
| Ammonia (secretion) | 1 | 2.04 | 0.36 | 2.56 | 0.02 | 2.47 | 0.16 | 5.04 | 0.21 | 3.54 | 0.32 | - | - | - | - | - | - |
| | 2 | 2.59 | 0.12 | 4.65 | 0.70 | 2.87 | 0.39 | 6.62 | 0.16 | 3.28 | 0.39 | 7.04 | 0.59 | 4.03 | 0.91 | - | - |
| | 3 | 2.08 | 0.05 | 3.30 | 0.18 | 2.79 | 0.10 | 4.20 | 0.10 | 5.83 | 0.43 | - | - | - | - | 4.54 | 0.12 |
| | 4 | 1.90 | 0.35 | 1.76 | 0.20 | 0.99 | 0.14 | 2.68 | 0.37 | 1.56 | 0.14 | 1.70 | 0.39 | 1.70 | 0.11 | 3.67 | 0.39 |

Values represent the mean and standard error (n=3).

**Supplementary Table 6.** Labeling Patterns of the TCA Cycle Intermediates with [U-$^{13}C_6$]Glucose

***Donors 1 and 4- TCA cycle labeling with [U-$^{13}C_6$] glucose***

| Metabolite | Labeling | EGFRt | s.e.m. | CD19 | s.e.m. | Leu16 | s.e.m. | Rituximab | s.e.m. | RFR-LCDR | s.e.m. | Rituximab.AA | s.e.m. | RFR-LCDR.AA | s.e.m. | GD2 | s.e.m. |
|---|---|---|---|---|---|---|---|---|---|---|---|---|---|---|---|---|---|
| Citrate | M+0 | 0.214 | 0.009 | 0.229 | 0.020 | 0.216 | 0.016 | 0.192 | 0.012 | 0.164 | 0.004 | 0.154 | 0.002 | 0.147 | 0.003 | 0.199 | 0.004 |
| | M+1 | 0.005 | 0.001 | 0.004 | 0.001 | 0.004 | 0.001 | 0.003 | 0.001 | 0.004 | 0.001 | 0.004 | 0.000 | 0.005 | 0.001 | 0.006 | 0.000 |
| | M+2 | 0.358 | 0.008 | 0.364 | 0.006 | 0.339 | 0.007 | 0.408 | 0.015 | 0.400 | 0.018 | 0.373 | 0.004 | 0.359 | 0.004 | 0.383 | 0.007 |
| | M+3 | 0.062 | 0.002 | 0.055 | 0.002 | 0.058 | 0.001 | 0.044 | 0.007 | 0.043 | 0.006 | 0.053 | 0.004 | 0.058 | 0.001 | 0.057 | 0.001 |
| | M+4 | 0.178 | 0.003 | 0.174 | 0.004 | 0.183 | 0.003 | 0.171 | 0.008 | 0.181 | 0.005 | 0.189 | 0.002 | 0.196 | 0.002 | 0.171 | 0.007 |
| | M+5 | 0.134 | 0.004 | 0.125 | 0.005 | 0.140 | 0.001 | 0.126 | 0.005 | 0.142 | 0.004 | 0.148 | 0.001 | 0.153 | 0.001 | 0.123 | 0.001 |
| | M+6 | 0.048 | 0.004 | 0.047 | 0.006 | 0.061 | 0.005 | 0.057 | 0.006 | 0.065 | 0.006 | 0.080 | 0.000 | 0.084 | 0.005 | 0.060 | 0.003 |
| α-ketoglutarate | M+0 | 0.685 | 0.017 | 0.680 | 0.021 | 0.646 | 0.019 | 0.776 | 0.017 | 0.748 | 0.020 | 0.676 | 0.007 | 0.667 | 0.003 | 0.744 | 0.010 |
| | M+1 | 0.000 | 0.000 | 0.000 | 0.000 | 0.000 | 0.000 | 0.000 | 0.000 | 0.000 | 0.000 | 0.000 | 0.000 | 0.000 | 0.000 | 0.000 | 0.000 |
| | M+2 | 0.116 | 0.004 | 0.137 | 0.008 | 0.127 | 0.008 | 0.095 | 0.006 | 0.101 | 0.006 | 0.121 | 0.007 | 0.117 | 0.005 | 0.119 | 0.007 |
| | M+3 | 0.051 | 0.004 | 0.047 | 0.004 | 0.050 | 0.004 | 0.030 | 0.003 | 0.037 | 0.003 | 0.040 | 0.000 | 0.046 | 0.007 | 0.031 | 0.003 |
| | M+4 | 0.077 | 0.004 | 0.078 | 0.007 | 0.088 | 0.005 | 0.052 | 0.005 | 0.056 | 0.009 | 0.085 | 0.004 | 0.080 | 0.003 | 0.057 | 0.000 |
| | M+5 | 0.071 | 0.007 | 0.059 | 0.004 | 0.089 | 0.004 | 0.047 | 0.004 | 0.058 | 0.004 | 0.078 | 0.002 | 0.090 | 0.005 | 0.048 | 0.003 |
| Fumarate | M+0 | 0.702 | 0.024 | 0.698 | 0.008 | 0.691 | 0.012 | 0.706 | 0.016 | 0.621 | 0.037 | 0.721 | 0.009 | 0.691 | 0.010 | 0.725 | 0.020 |
| | M+1 | 0.001 | 0.001 | 0.001 | 0.001 | 0.002 | 0.002 | 0.001 | 0.001 | 0.002 | 0.001 | 0.000 | 0.000 | 0.001 | 0.001 | 0.000 | 0.000 |
| | M+2 | 0.175 | 0.013 | 0.169 | 0.005 | 0.160 | 0.003 | 0.153 | 0.006 | 0.179 | 0.008 | 0.142 | 0.002 | 0.168 | 0.014 | 0.160 | 0.008 |
| | M+3 | 0.114 | 0.009 | 0.132 | 0.004 | 0.131 | 0.007 | 0.138 | 0.008 | 0.146 | 0.007 | 0.137 | 0.007 | 0.140 | 0.006 | 0.115 | 0.013 |
| | M+4 | 0.009 | 0.008 | 0.000 | 0.000 | 0.016 | 0.010 | 0.002 | 0.002 | 0.052 | 0.023 | 0.000 | 0.000 | 0.000 | 0.000 | 0.000 | 0.000 |
| Malate | M+0 | 0.547 | 0.012 | 0.550 | 0.012 | 0.562 | 0.024 | 0.604 | 0.029 | 0.563 | 0.030 | 0.500 | 0.006 | 0.486 | 0.009 | 0.559 | 0.008 |
| | M+1 | 0.017 | 0.001 | 0.015 | 0.003 | 0.017 | 0.001 | 0.009 | 0.004 | 0.009 | 0.004 | 0.011 | 0.000 | 0.011 | 0.002 | 0.022 | 0.001 |
| | M+2 | 0.188 | 0.004 | 0.190 | 0.004 | 0.167 | 0.010 | 0.178 | 0.005 | 0.187 | 0.004 | 0.195 | 0.001 | 0.194 | 0.005 | 0.184 | 0.002 |
| | M+3 | 0.157 | 0.004 | 0.160 | 0.005 | 0.153 | 0.008 | 0.134 | 0.010 | 0.152 | 0.010 | 0.173 | 0.002 | 0.186 | 0.003 | 0.147 | 0.002 |
| | M+4 | 0.092 | 0.004 | 0.085 | 0.007 | 0.101 | 0.005 | 0.076 | 0.011 | 0.090 | 0.013 | 0.121 | 0.005 | 0.123 | 0.003 | 0.088 | 0.006 |

Values represent the mean and standard error (n=6) for EGFRt, CD19, Leu16, Rituximab, and RFR-LCDR.
Values represent the mean and standard error (n=3) for Rituximab.AA, RFR-LCDR.AA, and GD2.

**Supplementary Table 7.** Labeling Patterns of the TCA Cycle Intermediates with 50% [U-$^{13}C_5$]Glutamine

***Donor 3-TCA labeling with 50% [U-$^{13}C_5$] glutamine and 50% unlabeled glutamine***

| Metabolite | Labeling | EGFRt | s.e.m. | CD19 | s.e.m. | Leu16 | s.e.m. | Rituximab | s.e.m. | RFR-LCDR | s.e.m. | GD2 | s.e.m. |
|---|---|---|---|---|---|---|---|---|---|---|---|---|---|
| Citrate | M+0 | 0.669 | 0.041 | 0.683 | 0.010 | 0.651 | 0.014 | 0.654 | 0.029 | 0.671 | 0.011 | 0.664 | 0.007 |
| | M+1 | 0.088 | 0.022 | 0.071 | 0.007 | 0.071 | 0.006 | 0.070 | 0.010 | 0.065 | 0.001 | 0.060 | 0.009 |
| | M+2 | 0.060 | 0.016 | 0.069 | 0.003 | 0.100 | 0.007 | 0.078 | 0.006 | 0.068 | 0.007 | 0.077 | 0.005 |
| | M+3 | 0.026 | 0.009 | 0.027 | 0.010 | 0.025 | 0.004 | 0.024 | 0.002 | 0.030 | 0.003 | 0.024 | 0.009 |
| | M+4 | 0.136 | 0.016 | 0.128 | 0.006 | 0.133 | 0.007 | 0.147 | 0.013 | 0.138 | 0.003 | 0.136 | 0.006 |
| | M+5 | 0.020 | 0.004 | 0.023 | 0.006 | 0.020 | 0.007 | 0.026 | 0.003 | 0.027 | 0.008 | 0.039 | 0.018 |
| | M+6 | 0.000 | 0.000 | 0.000 | 0.000 | 0.000 | 0.000 | 0.000 | 0.000 | 0.000 | 0.000 | 0.000 | 0.000 |
| α-ketoglutarate | M+0 | 0.694 | 0.009 | 0.676 | 0.010 | 0.691 | 0.010 | 0.666 | 0.003 | 0.709 | 0.006 | 0.700 | 0.005 |
| | M+1 | 0.041 | 0.003 | 0.032 | 0.003 | 0.032 | 0.004 | 0.037 | 0.003 | 0.021 | 0.002 | 0.031 | 0.003 |
| | M+2 | 0.007 | 0.001 | 0.007 | 0.001 | 0.007 | 0.000 | 0.009 | 0.001 | 0.004 | 0.001 | 0.008 | 0.001 |
| | M+3 | 0.061 | 0.004 | 0.066 | 0.004 | 0.058 | 0.002 | 0.067 | 0.001 | 0.051 | 0.002 | 0.054 | 0.002 |
| | M+4 | 0.000 | 0.000 | 0.000 | 0.000 | 0.000 | 0.000 | 0.000 | 0.000 | 0.000 | 0.000 | 0.000 | 0.000 |
| | M+5 | 0.197 | 0.003 | 0.218 | 0.002 | 0.212 | 0.007 | 0.221 | 0.001 | 0.215 | 0.003 | 0.208 | 0.005 |
| Fumarate | M+0 | 0.599 | 0.014 | 0.580 | 0.008 | 0.603 | 0.001 | 0.583 | 0.007 | 0.601 | 0.008 | 0.594 | 0.008 |
| | M+1 | 0.050 | 0.006 | 0.051 | 0.002 | 0.042 | 0.001 | 0.062 | 0.003 | 0.035 | 0.006 | 0.051 | 0.002 |
| | M+2 | 0.086 | 0.010 | 0.095 | 0.005 | 0.082 | 0.002 | 0.094 | 0.002 | 0.077 | 0.005 | 0.079 | 0.004 |
| | M+3 | 0.000 | 0.000 | 0.001 | 0.001 | 0.000 | 0.000 | 0.001 | 0.000 | 0.000 | 0.000 | 0.001 | 0.001 |
| | M+4 | 0.266 | 0.002 | 0.273 | 0.002 | 0.273 | 0.002 | 0.259 | 0.003 | 0.287 | 0.003 | 0.275 | 0.002 |
| Malate | M+0 | 0.638 | 0.008 | 0.619 | 0.012 | 0.652 | 0.011 | 0.613 | 0.007 | 0.648 | 0.008 | 0.631 | 0.008 |
| | M+1 | 0.063 | 0.001 | 0.063 | 0.005 | 0.055 | 0.004 | 0.066 | 0.002 | 0.049 | 0.001 | 0.059 | 0.004 |
| | M+2 | 0.086 | 0.005 | 0.090 | 0.004 | 0.084 | 0.005 | 0.094 | 0.003 | 0.079 | 0.001 | 0.087 | 0.003 |
| | M+3 | 0.006 | 0.000 | 0.009 | 0.001 | 0.005 | 0.001 | 0.009 | 0.000 | 0.008 | 0.001 | 0.008 | 0.000 |
| | M+4 | 0.208 | 0.003 | 0.219 | 0.004 | 0.203 | 0.003 | 0.218 | 0.007 | 0.216 | 0.006 | 0.215 | 0.002 |

Values represent the mean and standard error (n=3).

**Supplementary Table 8.** Carbon Mapping in Glycolysis, the Pentose Phosphate Pathway, the TCA Cycle, and Glutaminolysis

| *Carbon Mapping in Glycolysis, Pentose Phosphate Pathway, and TCA cycle* | | |
|---|---|---|
| **Reaction** | **Substrates** | **Products** |
| HK | GLC (ABCDEF) | G6P (ABCDEF) |
| PGI | G6P (ABCDEF) | F6P (ABCDEF) |
| PFK | F6P (ABCDEF) | FBP (ABCDEF) |
| FBA | FBP (ABCDEF) | DHAP (CBA) + GAP (DEF) |
| TPI | DHAP (ABC) | GAP (ABC) |
| GAPD | GAP (ABC) | 13BPG (ABC) |
| PGK | 13BPG (ABC) | 3PG (ABC) |
| ENO | 3PG (ABC) | PEP (ABC) |
| PYK | PEP (ABC) | PYR (ABC) |
| LDH | PYR (ABC) | LAC (ABC) |
| G6PDH | G6P (ABCDEF) | 6PG (ABCDEF) |
| GND | 6PG (ABCDEF) | RU5P (BCDEF) + CO2 (A) |
| RPI | RU5P (ABCDE) | R5P (ABCDE) |
| RPE | RU5P (ABCDE) | X5P (ABCDE) |
| TKT1 | X5P (ABCDE) + R5P (abcde) | GAP (CDE) + S7P (ABabcde) |
| TAL | GAP (ABC) + S7P (abcdefg) | F6P (abcABC) + E4P (defg) |
| TKT2 | X5P (ABCDE) + E4P (abcd) | GAP (CDE) + F6P (ABabcd) |
| PPCK | OAA (ABCD) | PEP (ABC) + CO2 (D) |
| ME | MAL (ABCD) | PYR (ABC) + CO2 (D) |
| PC | PYR (ABC) + CO2 (D) | OAA (ABCD) |
| PDH | PYR (ABC) | AcCoA (BC) + CO2 (A) |
| CS | OAA (ABCD) + AcCoA (ab) | CitICit (DCBAba) |
| ACITL | CitICit (DCBAba) | OAA (ABCD) + AcCoA_cyt (ab) |
| ICDH | CitICit (ABCDEF) | OGA (ABCEF) + CO2 (D) |
| AKGDH | OGA (ABCDE) | SuccCoA (BCDE) + CO2 (A) |
| SUCOAS | SuccCoA (ABCD) | Succ (ABCD) |
| SUCD | Succ (ABCD) | Fum (ABCD) |
| FUM | Fum (ABCD) | MAL (ABCD) |
| MDH | MAL (ABCD) | OAA (ABCD) |
| PYR_Ala | PYR (ABC) + Glu (DEFGH) | Ala (ABC) + OGA (DEFGH) |
| OGA_Glu | OGA (ABCDE) | Glu (ABCDE) |
| OGA_Gln | Glu (ABCDE) | Gln (ABCDE) |

for each reaction, the changes in letter positions indicate molecular transitions between substrates and product

G6P denotes glucose-6-phosphate; F6P, fructose-6-phosphate; FBP, fructose-1,6-bisphosphate; DHAP, dihydroxyacetone phosphate; GAP, glyceraldehyde-3-phosphate; 13BPG, 1,3-bisphosphoglycerate; 3PG, 3-phosphoglycerate; PEP, phosphoenolpyruvate; PYR, pyruvate, 6PG, 6-phosphogluconate; RU5P, ribulose-5-phosphate; R5P, ribose-5-phosphate; X5P, xyulose-5-phosphate; S7P, sedoheptulose-7-phosphate; E4P, erythrose-4-phosphate; OAA, oxaloacetate; MAL, malate; AcCoA, acetyl-CoA; CitICit, citrate/isocitrate; OGA, α-ketoglutarate; SuccCoA, succinyl-CoA; Succ, succinate; Fum, fumarate; Glu, glutamate; Ala, alanine; and Gln, glutamine. Succ and Fum are symmetric (i.e., ABCD = DCBA).

**Supplementary Table 9.** Central Carbon Metabolite Labeling Patterns from Various $^{13}C$ Tracers Used for Metabolic Flux Analysis

***Steady-state metabolite labeling from various $^{13}C$ tracers***

| | | **EGFRt T cells** | | | | | | | | | **Rituximab anti-CD20 CAR-T cells** | | | | | | | | |
|---|---|---|---|---|---|---|---|---|---|---|---|---|---|---|---|---|---|---|---|
| | | 1,2-$^{13}C_2$ Glucose | | | U-$^{13}C_6$ Glucose | | | 57% U-$^{13}C_5$ Glutamine | | | 1,2-$^{13}C_2$ Glucose | | | U-$^{13}C_6$ Glucose | | | 57% U-$^{13}C_5$ Glutamine | | |
| | | Measured | Simulated | Std. Dev. | Measured | Simulated | Std. Dev. | Measured | Simulated | Std. Dev. | Measured | Simulated | Std. Dev. | Measured | Simulated | Std. Dev. | Measured | Simulated | Std. Dev. |
| α-ketoglutarate | M+0 | 74.4% | 79.2% | 1.1% | 67.4% | 64.7% | 5.0% | 69.1% | 63.9% | 1.5% | 84.5% | 83.1% | 2.3% | 77.8% | 70.1% | 4.3% | 66.4% | 64.0% | 0.5% |
| | M+1 | 4.0% | 1.6% | 0.4% | 0.0% | 0.3% | 0.4% | 3.9% | 5.1% | 0.6% | 2.4% | 0.7% | 0.7% | 0.0% | 0.2% | 0.4% | 3.6% | 4.4% | 0.5% |
| | M+2 | 14.8% | 15.7% | 0.7% | 12.6% | 21.8% | 1.2% | 1.3% | 0.3% | 0.4% | 10.0% | 14.0% | 1.0% | 10.0% | 21.3% | 1.4% | 1.4% | 0.1% | 0.4% |
| | M+3 | 3.9% | 0.9% | 0.4% | 4.9% | 1.0% | 1.0% | 6.1% | 8.1% | 0.6% | 1.7% | 0.4% | 0.4% | 2.7% | 0.5% | 0.8% | 6.6% | 8.2% | 0.4% |
| | M+4 | 2.3% | 2.4% | 0.4% | 7.8% | 10.1% | 1.1% | 0.0% | 0.1% | 0.4% | 1.1% | 1.8% | 0.4% | 5.0% | 7.3% | 1.4% | 0.0% | 0.0% | 0.4% |
| | M+5 | 0.6% | 0.2% | 0.4% | 7.2% | 2.0% | 2.0% | 19.6% | 22.5% | 0.5% | 0.3% | 0.1% | 0.4% | 4.4% | 0.7% | 1.1% | 22.0% | 23.2% | 0.4% |
| alanine | M+0 | | | | 0.0% | 2.0% | 0.4% | 99.9% | 98.9% | 0.4% | | | | 0.0% | 0.9% | 0.4% | 99.7% | 99.5% | 0.4% |
| | M+1 | | | | 0.0% | 0.7% | 0.4% | 0.0% | 0.2% | 0.4% | | | | 0.0% | 0.3% | 0.4% | 0.0% | 0.1% | 0.4% |
| | M+2 | | | | 6.0% | 0.3% | 7.9% | 0.0% | 0.3% | 0.4% | | | | 0.0% | 0.1% | 0.4% | 0.0% | 0.1% | 0.4% |
| | M+3 | | | | 94.0% | 97.0% | 7.9% | 0.1% | 0.7% | 0.4% | | | | 100.0% | 98.7% | 0.4% | 0.2% | 0.3% | 0.4% |
| 2,3-bisphosphoglycerate | M+0 | | | | 0.0% | 0.1% | 0.4% | 99.7% | 99.9% | 0.5% | | | | 0.0% | 0.1% | 0.4% | 100.0% | 99.9% | 0.4% |
| | M+1 | | | | 0.0% | 0.0% | 0.4% | 0.2% | 0.0% | 0.4% | | | | 0.0% | 0.0% | 0.4% | 0.0% | 0.0% | 0.4% |
| | M+2 | | | | 0.0% | 0.0% | 0.4% | 0.0% | 0.0% | 0.4% | | | | 0.0% | 0.0% | 0.4% | 0.0% | 0.0% | 0.4% |
| | M+3 | | | | 100.0% | 99.8% | 0.4% | 0.0% | 0.0% | 0.4% | | | | 100.0% | 99.8% | 0.4% | 0.0% | 0.0% | 0.4% |
| citrate/isocitrate | M+0 | 48.9% | 48.6% | 1.5% | 22.1% | 14.6% | 3.4% | 66.0% | 65.0% | 7.0% | 54.7% | 55.2% | 5.5% | 18.9% | 22.4% | 3.0% | 64.6% | 64.6% | 4.9% |
| | M+1 | 6.2% | 5.0% | 0.4% | 0.6% | 2.1% | 0.4% | 8.7% | 6.3% | 3.7% | 5.4% | 3.1% | 0.9% | 0.4% | 1.7% | 0.4% | 7.0% | 5.1% | 1.6% |
| | M+2 | 31.3% | 35.9% | 1.8% | 35.3% | 47.6% | 1.9% | 6.9% | 7.2% | 2.6% | 29.5% | 34.4% | 3.0% | 40.6% | 50.0% | 3.6% | 8.6% | 7.0% | 0.9% |
| | M+3 | 6.0% | 3.4% | 0.7% | 6.0% | 4.9% | 0.6% | 2.7% | 1.0% | 1.7% | 4.8% | 2.2% | 1.1% | 4.4% | 4.5% | 1.6% | 2.5% | 1.6% | 0.4% |
| | M+4 | 5.8% | 6.0% | 0.4% | 17.8% | 15.3% | 1.2% | 13.5% | 18.4% | 2.8% | 4.3% | 4.5% | 0.4% | 17.4% | 13.0% | 1.9% | 14.6% | 19.3% | 2.3% |
| | M+5 | 1.4% | 0.8% | 0.4% | 13.2% | 11.0% | 1.1% | 2.1% | 1.8% | 0.7% | 1.0% | 0.5% | 0.4% | 12.5% | 6.8% | 1.1% | 2.7% | 2.2% | 0.5% |
| | M+6 | 0.4% | 0.2% | 0.4% | 5.1% | 4.5% | 1.0% | 0.0% | 0.3% | 0.4% | 0.3% | 0.1% | 0.4% | 5.9% | 1.5% | 1.4% | 0.0% | 0.2% | 0.4% |
| dihydroxyacetone phosphate | M+0 | 54.1% | 47.1% | 8.7% | 0.7% | 0.1% | 1.0% | 100.0% | 99.9% | 0.4% | 58.6% | 46.6% | 6.1% | 1.0% | 0.1% | 1.5% | 99.8% | 99.9% | 0.4% |
| | M+1 | 0.7% | 1.9% | 1.1% | 0.0% | 0.0% | 0.4% | 0.0% | 0.0% | 0.4% | 1.5% | 1.0% | 1.8% | 0.1% | 0.0% | 0.4% | 0.1% | 0.0% | 0.4% |
| | M+2 | 45.0% | 50.5% | 7.9% | 0.3% | 0.0% | 0.8% | 0.0% | 0.0% | 0.4% | 39.6% | 51.9% | 6.0% | 0.0% | 0.0% | 0.4% | 0.0% | 0.0% | 0.4% |
| | M+3 | 0.2% | 0.5% | 0.4% | 98.9% | 99.9% | 1.7% | 0.0% | 0.0% | 0.4% | 0.3% | 0.5% | 0.5% | 98.9% | 99.9% | 1.7% | 0.0% | 0.0% | 0.4% |
| fructose-1,6-bisphosphate* | M+0 | 13.8% | 11.3% | 4.6% | 0.0% | 0.0% | 0.4% | 100.0% | 99.9% | 0.4% | 13.9% | 11.8% | 3.2% | 0.0% | 0.0% | 0.4% | 100.0% | 99.9% | 0.4% |
| | M+1 | 3.4% | 2.0% | 3.7% | 0.0% | 0.0% | 0.4% | 0.0% | 0.0% | 0.4% | 3.2% | 1.0% | 4.0% | 0.0% | 0.0% | 0.4% | 0.0% | 0.0% | 0.4% |
| | M+2 | 67.9% | 64.3% | 10.4% | 0.0% | 0.0% | 0.4% | 0.0% | 0.0% | 0.4% | 67.5% | 67.8% | 4.0% | 0.0% | 0.0% | 0.4% | 0.0% | 0.0% | 0.4% |
| | M+3 | 1.2% | 2.4% | 1.8% | 0.0% | 0.1% | 0.4% | 0.0% | 0.0% | 0.4% | 1.6% | 1.4% | 1.9% | 0.0% | 0.1% | 0.4% | 0.0% | 0.0% | 0.4% |
| | M+4 | 13.7% | 19.6% | 4.6% | 0.0% | 0.0% | 0.4% | 0.0% | 0.0% | 0.4% | 13.7% | 17.6% | 3.2% | 0.0% | 0.0% | 0.4% | 0.0% | 0.0% | 0.4% |
| | M+5 | 0.0% | 0.6% | 0.4% | 0.0% | 0.0% | 0.4% | 0.0% | 0.0% | 0.4% | 0.0% | 0.5% | 0.4% | 0.1% | 0.0% | 0.4% | 0.0% | 0.0% | 0.4% |
| | M+6 | 0.0% | 0.0% | 0.4% | 99.9% | 99.8% | 0.4% | 0.0% | 0.0% | 0.4% | 0.0% | 0.0% | 0.4% | 99.8% | 99.8% | 0.4% | 0.0% | 0.0% | 0.4% |
| fumarate | M+0 | 81.0% | 79.3% | 0.7% | 71.3% | 64.8% | 5.3% | 59.9% | 64.5% | 2.4% | 88.3% | 83.1% | 3.0% | 76.4% | 70.1% | 4.1% | 58.3% | 64.2% | 1.2% |
| | M+1 | 3.8% | 4.5% | 0.4% | 0.1% | 2.2% | 0.4% | 5.0% | 4.7% | 1.0% | 0.7% | 3.4% | 1.0% | 0.0% | 2.4% | 0.4% | 6.3% | 4.3% | 0.6% |
| | M+2 | 11.9% | 13.2% | 0.7% | 16.1% | 20.5% | 1.9% | 8.6% | 8.3% | 1.7% | 9.1% | 11.5% | 2.5% | 13.7% | 19.3% | 1.4% | 9.4% | 8.3% | 0.4% |
| | M+3 | 3.3% | 2.3% | 0.4% | 12.3% | 8.2% | 3.3% | 0.1% | 0.1% | 0.4% | 1.8% | 1.6% | 0.4% | 9.8% | 6.4% | 3.0% | 0.1% | 0.0% | 0.4% |
| | M+4 | 0.0% | 0.8% | 0.4% | 0.2% | 4.3% | 0.4% | 26.5% | 22.5% | 0.4% | 0.0% | 0.4% | 0.4% | 0.0% | 1.8% | 0.4% | 25.9% | 23.2% | 0.6% |
| glucose-6-phosphate | M+0 | | | | 0.0% | 0.0% | 0.4% | 100.0% | 100.0% | 0.4% | | | | 0.0% | 0.0% | 0.4% | 100.0% | 100.0% | 0.4% |
| | M+1 | | | | 0.0% | 0.0% | 0.4% | 0.0% | 0.0% | 0.4% | | | | 0.0% | 0.0% | 0.4% | 0.0% | 0.0% | 0.4% |
| | M+2 | | | | 0.0% | 0.0% | 0.4% | 0.0% | 0.0% | 0.4% | | | | 0.0% | 0.0% | 0.4% | 0.0% | 0.0% | 0.4% |
| | M+3 | | | | 0.8% | 0.0% | 0.6% | 0.0% | 0.0% | 0.4% | | | | 1.2% | 0.0% | 0.8% | 0.0% | 0.0% | 0.4% |
| | M+4 | | | | 0.0% | 0.0% | 0.4% | 0.0% | 0.0% | 0.4% | | | | 0.0% | 0.0% | 0.4% | 0.0% | 0.0% | 0.4% |
| | M+5 | | | | 0.1% | 0.0% | 0.4% | 0.0% | 0.0% | 0.4% | | | | 0.6% | 0.0% | 0.8% | 0.0% | 0.0% | 0.4% |
| | M+6 | | | | 99.1% | 99.9% | 0.7% | 0.0% | 0.0% | 0.4% | | | | 98.1% | 100.0% | 1.4% | 0.0% | 0.0% | 0.4% |
| glutamine | M+0 | 95.0% | 99.9% | 5.0% | 99.9% | 99.9% | 0.4% | 43.6% | 43.1% | 1.4% | 97.1% | 100.0% | 2.9% | 99.9% | 100.0% | 0.4% | 41.7% | 43.0% | 0.4% |
| | M+1 | 1.0% | 0.0% | 1.1% | 0.0% | 0.0% | 0.4% | 0.0% | 0.0% | 0.4% | 0.2% | 0.0% | 0.4% | 0.0% | 0.0% | 0.4% | 0.0% | 0.0% | 0.4% |
| | M+2 | 2.7% | 0.1% | 2.7% | 0.0% | 0.1% | 0.4% | 0.0% | 0.0% | 0.4% | 1.9% | 0.0% | 1.9% | 0.0% | 0.0% | 0.4% | 0.0% | 0.0% | 0.4% |
| | M+3 | 0.8% | 0.0% | 0.8% | 0.0% | 0.0% | 0.4% | 0.0% | 0.0% | 0.4% | 0.5% | 0.0% | 0.5% | 0.0% | 0.0% | 0.4% | 0.0% | 0.0% | 0.4% |
| | M+4 | 0.4% | 0.0% | 0.4% | 0.0% | 0.0% | 0.4% | 0.0% | 0.0% | 0.4% | 0.2% | 0.0% | 0.4% | 0.0% | 0.0% | 0.4% | 0.0% | 0.0% | 0.4% |
| | M+5 | 0.1% | 0.0% | 0.4% | 0.0% | 0.0% | 0.4% | 56.4% | 56.8% | 1.4% | 0.1% | 0.0% | 0.4% | 0.0% | 0.0% | 0.4% | 58.3% | 57.0% | 0.4% |
| glutamate | M+0 | 78.7% | 82.0% | 9.9% | 54.6% | 69.4% | 3.9% | 64.6% | 63.8% | 3.1% | 87.4% | 85.1% | 5.3% | 65.6% | 73.7% | 6.1% | 63.8% | 63.9% | 0.5% |
| | M+1 | 4.3% | 1.4% | 1.7% | 0.4% | 0.3% | 0.4% | 5.2% | 4.4% | 1.0% | 2.1% | 0.6% | 1.1% | 0.4% | 0.1% | 0.4% | 5.0% | 3.9% | 0.4% |
| | M+2 | 10.7% | 13.6% | 4.3% | 16.0% | 18.9% | 0.6% | 2.6% | 0.2% | 0.6% | 7.2% | 12.3% | 2.9% | 14.2% | 18.7% | 1.1% | 2.3% | 0.1% | 0.4% |
| | M+3 | 3.6% | 0.8% | 2.0% | 7.4% | 0.9% | 0.9% | 7.7% | 7.1% | 1.0% | 1.5% | 0.3% | 1.1% | 4.8% | 0.4% | 1.4% | 7.7% | 7.2% | 0.4% |
| | M+4 | 2.3% | 2.1% | 1.5% | 11.1% | 8.8% | 1.2% | 0.0% | 0.0% | 0.4% | 1.0% | 1.5% | 0.8% | 8.1% | 6.4% | 1.7% | 0.0% | 0.0% | 0.4% |
| | M+5 | 0.5% | 0.1% | 0.5% | 10.5% | 1.8% | 1.7% | 19.9% | 24.5% | 0.6% | 0.8% | 0.1% | 1.2% | 7.0% | 0.6% | 1.9% | 21.2% | 24.9% | 0.8% |
| lactate | M+0 | 52.1% | 51.8% | 1.2% | 4.4% | 2.0% | 0.9% | 98.1% | 98.9% | 0.4% | 51.3% | 51.0% | 0.4% | 3.5% | 0.9% | 0.8% | 98.5% | 99.5% | 0.4% |
| | M+1 | 0.6% | 2.2% | 0.4% | 0.3% | 0.7% | 0.4% | 0.7% | 0.2% | 0.4% | 0.3% | 1.0% | 0.4% | 0.2% | 0.3% | 0.4% | 0.5% | 0.1% | 0.4% |
| | M+2 | 46.6% | 45.6% | 0.7% | 0.9% | 0.3% | 0.4% | 0.2% | 0.3% | 0.4% | 47.9% | 47.5% | 0.4% | 0.8% | 0.1% | 0.4% | 0.1% | 0.1% | 0.4% |
| | M+3 | 0.7% | 0.5% | 0.4% | 94.4% | 97.0% | 0.8% | 1.1% | 0.7% | 0.4% | 0.4% | 0.5% | 0.4% | 95.4% | 98.7% | 0.9% | 0.9% | 0.3% | 0.4% |
| malate | M+0 | 70.1% | 76.6% | 2.0% | 54.5% | 59.1% | 3.5% | 63.2% | 66.3% | 1.4% | 77.8% | 82.1% | 3.6% | 59.9% | 67.7% | 7.1% | 60.8% | 65.1% | 1.3% |
| | M+1 | 8.7% | 4.6% | 0.7% | 1.6% | 2.3% | 0.4% | 6.3% | 5.5% | 0.4% | 6.0% | 3.3% | 1.2% | 0.9% | 2.6% | 1.0% | 6.6% | 4.5% | 0.4% |
| | M+2 | 14.9% | 15.8% | 0.7% | 19.2% | 18.6% | 0.9% | 9.2% | 7.5% | 0.9% | 11.8% | 12.6% | 1.5% | 18.2% | 18.3% | 1.2% | 9.9% | 7.8% | 0.5% |
| | M+3 | 5.2% | 2.3% | 0.5% | 15.5% | 14.1% | 1.0% | 0.6% | 0.4% | 0.4% | 3.7% | 1.6% | 0.7% | 13.3% | 9.2% | 2.4% | 1.0% | 0.7% | 0.4% |
| | M+4 | 1.1% | 0.7% | 0.4% | 9.2% | 6.0% | 1.4% | 20.7% | 20.3% | 0.5% | 0.7% | 0.3% | 0.4% | 7.7% | 2.2% | 2.8% | 21.7% | 21.9% | 1.1% |
| phosphoenolpyruvate | M+0 | 62.6% | 51.0% | 9.9% | 1.0% | 0.2% | 0.9% | 100.0% | 99.9% | 0.4% | 61.8% | 50.9% | 10.7% | 0.8% | 0.6% | 0.9% | 100.0% | 99.7% | 0.4% |
| | M+1 | 0.0% | 1.8% | 0.4% | 0.0% | 0.1% | 0.4% | 0.0% | 0.0% | 0.4% | 0.2% | 1.0% | 0.6% | 0.0% | 0.2% | 0.4% | 0.0% | 0.0% | 0.4% |
| | M+2 | 37.1% | 46.7% | 9.6% | 0.1% | 0.0% | 0.4% | 0.0% | 0.0% | 0.4% | 37.9% | 47.7% | 10.5% | 0.0% | 0.1% | 0.4% | 0.0% | 0.1% | 0.4% |
| | M+3 | 0.2% | 0.5% | 0.4% | 98.9% | 99.7% | 1.1% | 0.0% | 0.1% | 0.4% | 0.0% | 0.5% | 0.4% | 99.2% | 99.2% | 0.9% | 0.0% | 0.2% | 0.4% |
| 6-phospho-D-gluconate | M+0 | 12.2% | 0.1% | 3.5% | 0.0% | 0.0% | 0.4% | 100.0% | 100.0% | 0.4% | 17.6% | 0.2% | 6.3% | 0.0% | 0.0% | 0.4% | 100.0% | 100.0% | 0.4% |
| | M+1 | 4.4% | 2.0% | 1.7% | 0.0% | 0.0% | 0.4% | 0.0% | 0.0% | 0.4% | 6.5% | 1.0% | 2.9% | 0.0% | 0.0% | 0.4% | 0.0% | 0.0% | 0.4% |
| | M+2 | 68.5% | 80.3% | 4.0% | 0.0% | 0.0% | 0.4% | 0.0% | 0.0% | 0.4% | 63.9% | 87.6% | 8.3% | 0.0% | 0.0% | 0.4% | 0.0% | 0.0% | 0.4% |
| | M+3 | 2.9% | 2.3% | 1.1% | 0.0% | 0.0% | 0.4% | 0.0% | 0.0% | 0.4% | 2.8% | 1.3% | 1.5% | 0.0% | 0.0% | 0.4% | 0.0% | 0.0% | 0.4% |
| | M+4 | 11.9% | 14.8% | 1.9% | 0.0% | 0.0% | 0.4% | 0.0% | 0.0% | 0.4% | 9.0% | 9.5% | 1.4% | 0.0% | 0.0% | 0.4% | 0.0% | 0.0% | 0.4% |
| | M+5 | 0.0% | 0.6% | 0.4% | 0.0% | 0.1% | 0.4% | 0.0% | 0.0% | 0.4% | 0.3% | 0.5% | 0.4% | 0.0% | 0.0% | 0.4% | 0.0% | 0.0% | 0.4% |
| | M+6 | 0.0% | 0.0% | 0.4% | 100.0% | 99.9% | 0.4% | 0.0% | 0.0% | 0.4% | 0.0% | 0.0% | 0.4% | 100.0% | 99.9% | 0.4% | 0.0% | 0.0% | 0.4% |
| 3-phosphoglycerate | M+0 | 56.5% | 51.0% | 3.5% | 0.4% | 0.2% | 0.5% | 100.0% | 99.9% | 0.4% | 58.1% | 50.8% | 1.8% | 0.5% | 0.5% | 0.7% | 99.7% | 99.7% | 0.4% |
| | M+1 | 0.4% | 1.8% | 0.6% | 0.0% | 0.1% | 0.4% | 0.0% | 0.0% | 0.4% | 0.2% | 1.0% | 0.4% | 0.0% | 0.2% | 0.4% | 0.3% | 0.0% | 0.4% |
| | M+2 | 42.8% | 46.7% | 2.3% | 0.0% | 0.0% | 0.4% | 0.0% | 0.0% | 0.4% | 41.5% | 47.7% | 1.4% | 0.8% | 0.1% | 1.2% | 0.0% | 0.1% | 0.4% |
| | M+3 | 0.3% | 0.5% | 0.5% | 99.5% | 99.7% | 0.5% | 0.0% | 0.1% | 0.4% | 0.2% | 0.5% | 0.4% | 98.7% | 99.2% | 1.3% | 0.0% | 0.2% | 0.4% |
| pyruvate | M+0 | 53.1% | 51.8% | 2.7% | 5.7% | 2.0% | 1.1% | 99.5% | 98.9% | 0.4% | 51.7% | 51.0% | 1.9% | 4.3% | 0.9% | 1.7% | 99.6% | 99.5% | 0.4% |
| | M+1 | 0.5% | 2.2% | 0.5% | 0.1% | 0.7% | 0.4% | 0.5% | 0.2% | 0.4% | 0.4% | 1.0% | 0.4% | 0.1% | 0.3% | 0.4% | 0.4% | 0.1% | 0.4% |
| | M+2 | 46.0% | 45.6% | 1.8% | 0.7% | 0.3% | 0.4% | 0.0% | 0.3% | 0.4% | 47.6% | 47.5% | 1.4% | 0.7% | 0.1% | 0.4% | 0.0% | 0.1% | 0.4% |
| | M+3 | 0.5% | 0.5% | 0.5% | 93.5% | 97.0% | 0.9% | 0.0% | 0.7% | 0.4% | 0.2% | 0.5% | 0.4% | 94.9% | 98.7% | 1.6% | 0.0% | 0.3% | 0.4% |
| ribose-5-phosphate | M+0 | 41.3% | 30.2% | 30.9% | 1.0% | 0.0% | 1.1% | 100.0% | 100.0% | 0.4% | 31.5% | 22.9% | 21.3% | 0.6% | 0.0% | 0.9% | 100.0% | 100.0% | 0.4% |
| | M+1 | 19.5% | 37.2% | 5.0% | 0.0% | 0.0% | 0.4% | 0.0% | 0.0% | 0.4% | 24.5% | 39.5% | 8.3% | 0.0% | 0.0% | 0.4% | 0.0% | 0.0% | 0.4% |
| | M+2 | 18.1% | 13.6% | 12.1% | 0.2% | 0.1% | 0.4% | 0.0% | 0.0% | 0.4% | 19.2% | 14.7% | 5.1% | 0.0% | 0.1% | 0.4% | 0.0% | 0.0% | 0.4% |
| | M+3 | 7.4% | 12.8% | 6.6% | 0.4% | 0.0% | 0.6% | 0.0% | 0.0% | 0.4% | 13.3% | 12.3% | 5.3% | 0.2% | 0.0% | 0.4% | 0.0% | 0.0% | 0.4% |
| | M+4 | 13.6% | 6.2% | 8.3% | 0.0% | 0.0% | 0.4% | 0.0% | 0.0% | 0.4% | 11.5% | 10.4% | 3.2% | 0.1% | 0.0% | 0.4% | 0.0% | 0.0% | 0.4% |
| | M+5 | 0.0% | 0.1% | 0.4% | 98.4% | 99.9% | 1.8% | 0.0% | 0.0% | 0.4% | 0.0% | 0.1% | 0.4% | 99.1% | 99.9% | 1.2% | 0.0% | 0.0% | 0.4% |
| sedoheptulose-7-phosphate | M+0 | | | | 0.5% | 0.0% | 0.7% | 100.0% | 100.0% | 0.4% | | | | 1.1% | 0.0% | 1.7% | 99.9% | 100.0% | 0.4% |
| | M+1 | | | | 0.0% | 0.0% | 0.4% | 0.0% | 0.0% | 0.4% | | | | 0.0% | 0.0% | 0.4% | 0.0% | 0.0% | 0.4% |
| | M+2 | | | | 0.0% | 0.0% | 0.4% | 0.0% | 0.0% | 0.4% | | | | 0.0% | 0.0% | 0.4% | 0.0% | 0.0% | 0.4% |
| | M+3 | | | | 0.2% | 0.0% | 0.4% | 0.0% | 0.0% | 0.4% | | | | 0.7% | 0.0% | 0.8% | 0.0% | 0.0% | 0.4% |
| | M+4 | | | | 0.3% | 0.1% | 0.5% | 0.0% | 0.0% | 0.4% | | | | 0.2% | 0.1% | 0.5% | 0.0% | 0.0% | 0.4% |
| | M+5 | | | | 0.4% | 0.0% | 0.6% | 0.0% | 0.0% | 0.4% | | | | 1.3% | 0.0% | 1.0% | 0.0% | 0.0% | 0.4% |
| | M+6 | | | | 0.9% | 0.0% | 1.2% | 0.0% | 0.0% | 0.4% | | | | 1.1% | 0.0% | 1.3% | 0.0% | 0.0% | 0.4% |
| | M+7 | | | | 97.7% | 99.9% | 2.0% | 0.0% | 0.0% | 0.4% | | | | 95.5% | 99.9% | 3.5% | 0.0% | 0.0% | 0.4% |

*fructose-1,6-bisphosphate M+0 and M+4 assumed to be equal for 1,2-$^{13}C_2$ Glucose labeling

Values represent the mean and standard deviation (n=3 or 6).

**Supplementary Table 10.** Metabolic Flux Distributions (nmol/million cells/hr)

| ***Metabolic flux distributions (nmol/million cells/hr) determined using $^{13}C$ tracers*** | | | | | | | | |
|---|---|---|---|---|---|---|---|---|
| | | | **EGFRt T cells** | | | **Rituximab CAR-T cells** | | |
| **Reaction** | **Substrates** | **Products** | **flux** | L.B. | U.B. | **flux** | L.B. | U.B. |
| 'IN_GLC' | | GLC | 63.36 | 63.36 | 64.64 | 123.19 | 114.81 | 138.64 |
| 'hk' | GLC | G6P | 63.36 | 63.36 | 64.64 | 123.19 | 114.81 | 138.64 |
| 'pgi' | G6P | F6P | 57.28 | 56.35 | 58.43 | 116.98 | 108.55 | 131.74 |
| 'pfk' | F6P | FBP | 60.90 | 60.63 | 62.14 | 120.40 | 112.01 | 135.62 |
| 'fba' | FBP | DHAP + GAP | 60.90 | 60.63 | 62.14 | 120.40 | 112.01 | 135.62 |
| 'tpi' | DHAP | GAP | 60.54 | 60.27 | 61.78 | 119.80 | 111.41 | 135.03 |
| 'gapd' | GAP | 13BPG | 123.26 | 122.99 | 125.76 | 241.91 | 225.14 | 272.60 |
| 'pgk' | 13BPG | 3PG | 123.26 | 122.99 | 125.76 | 241.91 | 225.14 | 272.60 |
| 'eno' | 2PG | PEP | 112.01 | 111.74 | 114.52 | 220.93 | 204.16 | 251.63 |
| 'pyk' | PEP | Pyr | 112.45 | 111.98 | 114.86 | 223.01 | 206.00 | 253.98 |
| 'ldh' | Pyr | Lac | 82.37 | 79.29 | 84.03 | 183.99 | 158.23 | 207.94 |
| 'EX_PYR' | Pyr | | 1.58 | 1.56 | 1.75 | 2.06 | 1.86 | 2.27 |
| 'EX_LAC' | Lactate | | 82.37 | 79.29 | 84.03 | 183.99 | 158.23 | 207.94 |
| 'ppck' | OAA | PEP + CO2 | 0.44 | 0.06 | 0.52 | 2.09 | 0.99 | 2.63 |
| 'me' | Mal | Pyr + CO2 | 3.31 | 3.17 | 3.64 | 1.13 | 0.19 | 2.21 |
| 'pc' | Pyr + CO2 | OAA | 5.05 | 4.29 | 6.42 | 2.92 | 1.55 | 4.12 |
| 'g6pdh' | G6P | 6PG | 5.73 | 4.70 | 6.76 | 5.61 | 3.41 | 8.02 |
| 'gnd' | 6PG | Ru5P + CO2 | 5.73 | 4.70 | 6.76 | 5.61 | 3.41 | 8.02 |
| 'rpi' | Ru5P | R5P | 2.10 | 1.76 | 2.45 | 2.20 | 1.46 | 3.00 |
| 'rpe' | Ru5P | X5P | 3.62 | 2.93 | 4.31 | 3.42 | 1.95 | 5.02 |
| 'tkt2' | E4P + X5P | F6P + GAP | 1.81 | 1.47 | 2.16 | 1.71 | 0.97 | 2.51 |
| 'tkt1' | R5P + X5P | S7P + GAP | 1.81 | 1.47 | 2.16 | 1.71 | 0.97 | 2.51 |
| 'tal' | S7P + GAP | E4P + F6P | 1.81 | 1.47 | 2.16 | 1.71 | 0.97 | 2.51 |
| 'EX_OAA' | OAA | | 25.90 | 20.77 | 30.70 | 28.28 | 25.46 | 31.11 |
| 'pdh' | Pyr | AcCoA + CO2 | 25.73 | 24.60 | 30.46 | 32.10 | 26.45 | 48.52 |
| 'cs' | AcCoA + OAA | Cit/Icit | 31.27 | 30.34 | 37.60 | 43.61 | 35.22 | 68.67 |
| 'icdh' | Cit/Icit | OGA + CO2 | 23.96 | 19.59 | 28.58 | 26.13 | 23.81 | 28.53 |
| 'akgdh' | OGA | SuccCoA + CO2 | 48.55 | 39.81 | 57.01 | 54.72 | 50.06 | 59.37 |
| 'sucoas' | SuccCoA | Succ | 48.55 | 39.81 | 57.01 | 54.72 | 50.06 | 59.37 |
| 'sucd' | Succ | Fum | 48.55 | 39.81 | 57.01 | 54.72 | 50.06 | 59.37 |
| 'fum' | Fum | Mal | 48.55 | 39.81 | 57.01 | 54.72 | 50.06 | 59.37 |
| 'mdh' | Mal | OAA | 45.24 | 36.44 | 53.83 | 53.58 | 48.76 | 58.32 |
| 'PYR_Ala' | Pyr | Ala | 1.03 | 1.03 | 1.03 | 3.07 | 2.73 | 3.62 |
| 'EX_PYR_Ala' | Ala | | 1.03 | 1.03 | 1.03 | 3.07 | 2.73 | 3.62 |
| 'IN_AC' | | AcCoA | 5.55 | 4.93 | 7.14 | 11.51 | 7.94 | 20.15 |
| 'EX_AC_cyt' | AcCoA | | 7.32 | 1.95 | 15.44 | 17.48 | 7.87 | 43.06 |
| 'OGA_Glu' | OGA | Glu | -23.56 | -27.62 | -19.18 | -25.51 | -28.06 | -22.96 |
| 'OGA_Gln' | Glu | Gln | -25.55 | -29.60 | -21.18 | -32.63 | -34.91 | -30.46 |
| IN_Gln-EX_Gln | | Gln | 25.55 | 21.18 | 29.60 | 32.63 | 30.46 | 34.91 |
| EX_OGA_Glu-IN_Glu | Glu | | 0.96 | 0.96 | 0.97 | 4.04 | 3.70 | 4.59 |

L.B. and U.B. represent the lower and upper bounds of the 95% confidence interval.

**Supplementary Table 11.** Labeling Patterns of Nitrogenous Metabolites in CAR-T Cells with 50% [γ-$^{15}$N]Glutamine

| ***Donor 4- CAR-T cell metabolite labeling with [γ-$^{15}$N] glutamine*** | | | | | | | | | | | | | | | |
|---|---|---|---|---|---|---|---|---|---|---|---|---|---|---|---|
| **Metabolite** | **Labeling** | **EGFRt** | **s.e.m.** | **CD19** | **s.e.m.** | **Leu16** | **s.e.m.** | **Rituximab** | **s.e.m.** | **RFR-LCDR** | **s.e.m.** | **Rituximab.AA** | **s.e.m.** | **RFR-LCDR.AA** | **s.e.m.** | **GD2** | **s.e.m.** |
| ATP | M+0 | 0.407 | 0.009 | 0.406 | 0.010 | 0.371 | 0.007 | 0.324 | 0.005 | 0.319 | 0.003 | 0.315 | 0.005 | 0.330 | 0.004 | 0.374 | 0.004 |
| | M+1 | 0.343 | 0.006 | 0.356 | 0.023 | 0.340 | 0.002 | 0.363 | 0.003 | 0.360 | 0.004 | 0.338 | 0.002 | 0.349 | 0.008 | 0.354 | 0.004 |
| | M+2 | 0.250 | 0.013 | 0.238 | 0.013 | 0.289 | 0.006 | 0.313 | 0.003 | 0.321 | 0.007 | 0.347 | 0.006 | 0.321 | 0.012 | 0.272 | 0.008 |
| | M+3 | 0.000 | 0.000 | 0.000 | 0.000 | 0.000 | 0.000 | 0.000 | 0.000 | 0.000 | 0.000 | 0.000 | 0.000 | 0.000 | 0.000 | 0.000 | 0.000 |
| | M+4 | 0.000 | 0.000 | 0.000 | 0.000 | 0.000 | 0.000 | 0.000 | 0.000 | 0.000 | 0.000 | 0.000 | 0.000 | 0.000 | 0.000 | 0.000 | 0.000 |
| | M+5 | 0.000 | 0.000 | 0.000 | 0.000 | 0.000 | 0.000 | 0.000 | 0.000 | 0.000 | 0.000 | 0.000 | 0.000 | 0.000 | 0.000 | 0.000 | 0.000 |
| ADP | M+0 | 0.258 | 0.001 | 0.246 | 0.003 | 0.230 | 0.004 | 0.194 | 0.002 | 0.185 | 0.003 | 0.179 | 0.004 | 0.184 | 0.011 | 0.224 | 0.004 |
| | M+1 | 0.449 | 0.001 | 0.454 | 0.009 | 0.430 | 0.006 | 0.440 | 0.005 | 0.452 | 0.003 | 0.457 | 0.001 | 0.472 | 0.002 | 0.422 | 0.003 |
| | M+2 | 0.293 | 0.002 | 0.300 | 0.009 | 0.340 | 0.005 | 0.365 | 0.004 | 0.362 | 0.001 | 0.364 | 0.004 | 0.343 | 0.011 | 0.354 | 0.003 |
| | M+3 | 0.000 | 0.000 | 0.000 | 0.000 | 0.000 | 0.000 | 0.000 | 0.000 | 0.000 | 0.000 | 0.000 | 0.000 | 0.000 | 0.000 | 0.000 | 0.000 |
| | M+4 | 0.000 | 0.000 | 0.000 | 0.000 | 0.000 | 0.000 | 0.000 | 0.000 | 0.000 | 0.000 | 0.000 | 0.000 | 0.000 | 0.000 | 0.000 | 0.000 |
| | M+5 | 0.000 | 0.000 | 0.000 | 0.000 | 0.000 | 0.000 | 0.000 | 0.000 | 0.000 | 0.000 | 0.000 | 0.000 | 0.000 | 0.000 | 0.000 | 0.000 |
| AMP | M+0 | 0.341 | 0.009 | 0.305 | 0.005 | 0.300 | 0.006 | 0.253 | 0.005 | 0.248 | 0.003 | 0.253 | 0.005 | 0.261 | 0.002 | 0.280 | 0.003 |
| | M+1 | 0.417 | 0.004 | 0.433 | 0.004 | 0.404 | 0.009 | 0.419 | 0.001 | 0.433 | 0.005 | 0.426 | 0.006 | 0.440 | 0.007 | 0.409 | 0.003 |
| | M+2 | 0.242 | 0.008 | 0.262 | 0.006 | 0.295 | 0.004 | 0.328 | 0.005 | 0.319 | 0.004 | 0.320 | 0.002 | 0.299 | 0.008 | 0.311 | 0.002 |
| | M+3 | 0.000 | 0.000 | 0.000 | 0.000 | 0.000 | 0.000 | 0.000 | 0.000 | 0.000 | 0.000 | 0.000 | 0.000 | 0.000 | 0.000 | 0.000 | 0.000 |
| | M+4 | 0.000 | 0.000 | 0.000 | 0.000 | 0.000 | 0.000 | 0.000 | 0.000 | 0.000 | 0.000 | 0.000 | 0.000 | 0.000 | 0.000 | 0.000 | 0.000 |
| | M+5 | 0.000 | 0.000 | 0.000 | 0.000 | 0.000 | 0.000 | 0.000 | 0.000 | 0.000 | 0.000 | 0.000 | 0.000 | 0.000 | 0.000 | 0.000 | 0.000 |
| CTP | M+0 | 0.333 | 0.015 | 0.327 | 0.014 | 0.299 | 0.015 | 0.280 | 0.008 | 0.261 | 0.012 | 0.265 | 0.004 | 0.298 | 0.018 | 0.286 | 0.021 |
| | M+1 | 0.574 | 0.019 | 0.576 | 0.012 | 0.567 | 0.009 | 0.571 | 0.003 | 0.584 | 0.014 | 0.567 | 0.007 | 0.562 | 0.024 | 0.533 | 0.021 |
| | M+2 | 0.094 | 0.010 | 0.097 | 0.004 | 0.134 | 0.011 | 0.149 | 0.006 | 0.155 | 0.005 | 0.168 | 0.008 | 0.141 | 0.008 | 0.181 | 0.003 |
| | M+3 | 0.000 | 0.000 | 0.000 | 0.000 | 0.000 | 0.000 | 0.000 | 0.000 | 0.000 | 0.000 | 0.000 | 0.000 | 0.000 | 0.000 | 0.000 | 0.000 |
| CDP | M+0 | 0.337 | 0.014 | 0.254 | 0.024 | 0.274 | 0.028 | 0.263 | 0.015 | 0.222 | 0.002 | 0.277 | 0.022 | 0.268 | 0.023 | 0.230 | 0.024 |
| | M+1 | 0.481 | 0.007 | 0.509 | 0.018 | 0.486 | 0.029 | 0.500 | 0.015 | 0.524 | 0.018 | 0.472 | 0.018 | 0.512 | 0.020 | 0.503 | 0.019 |
| | M+2 | 0.182 | 0.016 | 0.236 | 0.006 | 0.240 | 0.001 | 0.237 | 0.003 | 0.254 | 0.017 | 0.251 | 0.009 | 0.220 | 0.008 | 0.267 | 0.005 |
| | M+3 | 0.000 | 0.000 | 0.000 | 0.000 | 0.000 | 0.000 | 0.000 | 0.000 | 0.000 | 0.000 | 0.000 | 0.000 | 0.000 | 0.000 | 0.000 | 0.000 |
| CMP | M+0 | 0.220 | 0.001 | 0.295 | 0.041 | 0.491 | 0.079 | 0.295 | 0.043 | 0.246 | 0.058 | 0.389 | 0.010 | 0.256 | 0.019 | 0.320 | 0.017 |
| | M+1 | 0.470 | 0.073 | 0.493 | 0.080 | 0.332 | 0.073 | 0.499 | 0.040 | 0.508 | 0.066 | 0.350 | 0.059 | 0.516 | 0.120 | 0.415 | 0.018 |
| | M+2 | 0.310 | 0.073 | 0.211 | 0.040 | 0.177 | 0.095 | 0.206 | 0.029 | 0.246 | 0.074 | 0.261 | 0.049 | 0.228 | 0.121 | 0.265 | 0.024 |
| | M+3 | 0.000 | 0.000 | 0.000 | 0.000 | 0.000 | 0.000 | 0.000 | 0.000 | 0.000 | 0.000 | 0.000 | 0.000 | 0.000 | 0.000 | 0.000 | 0.000 |
| GTP | M+0 | 0.404 | 0.009 | 0.395 | 0.015 | 0.374 | 0.013 | 0.353 | 0.012 | 0.397 | 0.017 | 0.361 | 0.023 | 0.430 | 0.019 | 0.369 | 0.027 |
| | M+1 | 0.282 | 0.021 | 0.313 | 0.014 | 0.247 | 0.032 | 0.278 | 0.011 | 0.268 | 0.019 | 0.262 | 0.017 | 0.264 | 0.037 | 0.243 | 0.016 |
| | M+2 | 0.271 | 0.015 | 0.209 | 0.009 | 0.266 | 0.032 | 0.278 | 0.025 | 0.284 | 0.006 | 0.284 | 0.045 | 0.270 | 0.043 | 0.268 | 0.014 |
| | M+3 | 0.042 | 0.021 | 0.083 | 0.011 | 0.114 | 0.009 | 0.091 | 0.019 | 0.051 | 0.012 | 0.093 | 0.006 | 0.036 | 0.024 | 0.120 | 0.013 |
| | M+4 | 0.000 | 0.000 | 0.000 | 0.000 | 0.000 | 0.000 | 0.000 | 0.000 | 0.000 | 0.000 | 0.000 | 0.000 | 0.000 | 0.000 | 0.000 | 0.000 |
| | M+5 | 0.000 | 0.000 | 0.000 | 0.000 | 0.000 | 0.000 | 0.000 | 0.000 | 0.000 | 0.000 | 0.000 | 0.000 | 0.000 | 0.000 | 0.000 | 0.000 |
| GDP | M+0 | 0.260 | 0.025 | 0.333 | 0.019 | 0.277 | 0.003 | 0.240 | 0.016 | 0.234 | 0.007 | 0.217 | 0.008 | 0.204 | 0.011 | 0.259 | 0.016 |
| | M+1 | 0.233 | 0.039 | 0.224 | 0.030 | 0.191 | 0.023 | 0.204 | 0.008 | 0.217 | 0.012 | 0.209 | 0.009 | 0.212 | 0.009 | 0.218 | 0.005 |
| | M+2 | 0.291 | 0.010 | 0.280 | 0.007 | 0.319 | 0.020 | 0.321 | 0.012 | 0.360 | 0.014 | 0.309 | 0.015 | 0.301 | 0.030 | 0.347 | 0.003 |
| | M+3 | 0.217 | 0.071 | 0.163 | 0.011 | 0.213 | 0.037 | 0.235 | 0.029 | 0.189 | 0.019 | 0.264 | 0.015 | 0.282 | 0.037 | 0.176 | 0.018 |
| | M+4 | 0.000 | 0.000 | 0.000 | 0.000 | 0.000 | 0.000 | 0.000 | 0.000 | 0.000 | 0.000 | 0.000 | 0.000 | 0.000 | 0.000 | 0.000 | 0.000 |
| | M+5 | 0.000 | 0.000 | 0.000 | 0.000 | 0.000 | 0.000 | 0.000 | 0.000 | 0.000 | 0.000 | 0.000 | 0.000 | 0.000 | 0.000 | 0.000 | 0.000 |
| GMP | M+0 | 0.431 | 0.127 | 0.422 | 0.042 | 0.361 | 0.030 | 0.446 | 0.022 | 0.365 | 0.050 | 0.317 | 0.020 | 0.409 | 0.054 | 0.361 | 0.022 |
| | M+1 | 0.403 | 0.091 | 0.362 | 0.017 | 0.348 | 0.033 | 0.288 | 0.032 | 0.357 | 0.066 | 0.380 | 0.042 | 0.360 | 0.024 | 0.351 | 0.008 |
| | M+2 | 0.109 | 0.041 | 0.104 | 0.007 | 0.103 | 0.015 | 0.073 | 0.010 | 0.104 | 0.028 | 0.116 | 0.018 | 0.087 | 0.013 | 0.095 | 0.002 |
| | M+3 | 0.056 | 0.056 | 0.112 | 0.022 | 0.187 | 0.047 | 0.193 | 0.049 | 0.174 | 0.045 | 0.188 | 0.056 | 0.144 | 0.021 | 0.193 | 0.020 |
| | M+4 | 0.000 | 0.000 | 0.000 | 0.000 | 0.000 | 0.000 | 0.000 | 0.000 | 0.000 | 0.000 | 0.000 | 0.000 | 0.000 | 0.000 | 0.000 | 0.000 |
| | M+5 | 0.000 | 0.000 | 0.000 | 0.000 | 0.000 | 0.000 | 0.000 | 0.000 | 0.000 | 0.000 | 0.000 | 0.000 | 0.000 | 0.000 | 0.000 | 0.000 |
| UTP | M+0 | 0.408 | 0.008 | 0.405 | 0.005 | 0.351 | 0.008 | 0.346 | 0.003 | 0.339 | 0.005 | 0.358 | 0.008 | 0.350 | 0.004 | 0.342 | 0.004 |
| | M+1 | 0.592 | 0.008 | 0.595 | 0.005 | 0.649 | 0.008 | 0.654 | 0.003 | 0.661 | 0.005 | 0.642 | 0.008 | 0.650 | 0.004 | 0.658 | 0.004 |
| | M+2 | 0.000 | 0.000 | 0.000 | 0.000 | 0.000 | 0.000 | 0.000 | 0.000 | 0.000 | 0.000 | 0.000 | 0.000 | 0.000 | 0.000 | 0.000 | 0.000 |
| UDP | M+0 | 0.431 | 0.007 | 0.442 | 0.018 | 0.395 | 0.001 | 0.379 | 0.008 | 0.385 | 0.006 | 0.396 | 0.001 | 0.398 | 0.011 | 0.387 | 0.002 |
| | M+1 | 0.569 | 0.007 | 0.558 | 0.018 | 0.605 | 0.001 | 0.621 | 0.008 | 0.615 | 0.006 | 0.604 | 0.001 | 0.602 | 0.011 | 0.613 | 0.002 |
| | M+2 | 0.000 | 0.000 | 0.000 | 0.000 | 0.000 | 0.000 | 0.000 | 0.000 | 0.000 | 0.000 | 0.000 | 0.000 | 0.000 | 0.000 | 0.000 | 0.000 |
| UMP | M+0 | 0.401 | 0.005 | 0.455 | 0.006 | 0.418 | 0.019 | 0.363 | 0.011 | 0.397 | 0.012 | 0.374 | 0.015 | 0.387 | 0.009 | 0.380 | 0.014 |
| | M+1 | 0.599 | 0.005 | 0.545 | 0.006 | 0.582 | 0.019 | 0.637 | 0.011 | 0.603 | 0.012 | 0.626 | 0.015 | 0.613 | 0.009 | 0.620 | 0.014 |
| | M+2 | 0.000 | 0.000 | 0.000 | 0.000 | 0.000 | 0.000 | 0.000 | 0.000 | 0.000 | 0.000 | 0.000 | 0.000 | 0.000 | 0.000 | 0.000 | 0.000 |

Values represent the mean and standard error (n=3).

**Supplementary Table 12.** Labeling Patterns of Nucleotides and the Hexosamine Pathway Intermediates in CAR-T Cells with [1,2-$^{13}C_2$]Glucose

***Donor 5- CAR-T cell metabolite labeling with [1,2-$^{13}C_2$] glucose***

| Metabolite | Labeling | EGFRt | s.e.m. | CD19 | s.e.m. | Leu16 | s.e.m. | Rituximab | s.e.m. | RFR-LCDR | s.e.m. | Rituximab.AA | s.e.m. | RFR-LCDR.AA | s.e.m. |
|---|---|---|---|---|---|---|---|---|---|---|---|---|---|---|---|
| ATP | M+0 | 0.410 | 0.009 | 0.356 | 0.033 | 0.352 | 0.002 | 0.201 | 0.014 | 0.250 | 0.006 | 0.227 | 0.005 | 0.287 | 0.004 |
| | M+1 | 0.211 | 0.005 | 0.212 | 0.007 | 0.208 | 0.004 | 0.264 | 0.008 | 0.236 | 0.010 | 0.252 | 0.006 | 0.228 | 0.003 |
| | M+2 | 0.172 | 0.009 | 0.184 | 0.014 | 0.188 | 0.006 | 0.213 | 0.007 | 0.208 | 0.006 | 0.213 | 0.006 | 0.195 | 0.002 |
| | M+3 | 0.085 | 0.005 | 0.102 | 0.005 | 0.104 | 0.001 | 0.141 | 0.006 | 0.128 | 0.003 | 0.130 | 0.004 | 0.120 | 0.006 |
| | M+4 | 0.109 | 0.008 | 0.137 | 0.006 | 0.138 | 0.001 | 0.164 | 0.006 | 0.162 | 0.005 | 0.162 | 0.003 | 0.154 | 0.003 |
| | M+5 | 0.011 | 0.001 | 0.009 | 0.002 | 0.011 | 0.000 | 0.015 | 0.005 | 0.016 | 0.002 | 0.016 | 0.001 | 0.016 | 0.001 |
| | M+6 | 0.000 | 0.000 | 0.000 | 0.000 | 0.000 | 0.000 | 0.000 | 0.000 | 0.000 | 0.000 | 0.000 | 0.000 | 0.000 | 0.000 |
| | M+7 | 0.000 | 0.000 | 0.000 | 0.000 | 0.000 | 0.000 | 0.000 | 0.000 | 0.000 | 0.000 | 0.000 | 0.000 | 0.000 | 0.000 |
| | M+8 | 0.000 | 0.000 | 0.000 | 0.000 | 0.000 | 0.000 | 0.000 | 0.000 | 0.000 | 0.000 | 0.000 | 0.000 | 0.000 | 0.000 |
| | M+9 | 0.000 | 0.000 | 0.000 | 0.000 | 0.000 | 0.000 | 0.000 | 0.000 | 0.000 | 0.000 | 0.000 | 0.000 | 0.000 | 0.000 |
| | M+10 | 0.000 | 0.000 | 0.000 | 0.000 | 0.000 | 0.000 | 0.000 | 0.000 | 0.000 | 0.000 | 0.000 | 0.000 | 0.000 | 0.000 |
| ADP | M+0 | 0.447 | 0.005 | 0.367 | 0.023 | 0.390 | 0.007 | 0.179 | 0.004 | 0.262 | 0.003 | 0.225 | 0.003 | 0.309 | 0.006 |
| | M+1 | 0.188 | 0.004 | 0.199 | 0.008 | 0.187 | 0.005 | 0.270 | 0.008 | 0.248 | 0.001 | 0.246 | 0.002 | 0.224 | 0.006 |
| | M+2 | 0.150 | 0.003 | 0.174 | 0.014 | 0.177 | 0.003 | 0.212 | 0.002 | 0.196 | 0.001 | 0.207 | 0.002 | 0.188 | 0.001 |
| | M+3 | 0.085 | 0.001 | 0.099 | 0.006 | 0.089 | 0.004 | 0.144 | 0.006 | 0.119 | 0.002 | 0.135 | 0.003 | 0.108 | 0.002 |
| | M+4 | 0.123 | 0.003 | 0.156 | 0.004 | 0.149 | 0.004 | 0.181 | 0.006 | 0.162 | 0.003 | 0.176 | 0.002 | 0.161 | 0.003 |
| | M+5 | 0.004 | 0.000 | 0.001 | 0.000 | 0.004 | 0.001 | 0.009 | 0.000 | 0.009 | 0.001 | 0.008 | 0.001 | 0.005 | 0.001 |
| | M+6 | 0.000 | 0.000 | 0.000 | 0.000 | 0.000 | 0.000 | 0.000 | 0.000 | 0.000 | 0.000 | 0.000 | 0.000 | 0.000 | 0.000 |
| | M+7 | 0.000 | 0.000 | 0.000 | 0.000 | 0.000 | 0.000 | 0.000 | 0.000 | 0.000 | 0.000 | 0.000 | 0.000 | 0.000 | 0.000 |
| | M+8 | 0.000 | 0.000 | 0.000 | 0.000 | 0.000 | 0.000 | 0.000 | 0.000 | 0.000 | 0.000 | 0.000 | 0.000 | 0.000 | 0.000 |
| | M+9 | 0.000 | 0.000 | 0.000 | 0.000 | 0.000 | 0.000 | 0.000 | 0.000 | 0.000 | 0.000 | 0.000 | 0.000 | 0.000 | 0.000 |
| | M+10 | 0.000 | 0.000 | 0.000 | 0.000 | 0.000 | 0.000 | 0.000 | 0.000 | 0.000 | 0.000 | 0.000 | 0.000 | 0.000 | 0.000 |
| AMP | M+0 | 0.509 | 0.007 | 0.387 | 0.005 | 0.435 | 0.006 | 0.212 | 0.005 | 0.299 | 0.006 | 0.260 | 0.005 | 0.334 | 0.004 |
| | M+1 | 0.140 | 0.005 | 0.162 | 0.007 | 0.145 | 0.004 | 0.242 | 0.010 | 0.221 | 0.004 | 0.224 | 0.009 | 0.207 | 0.008 |
| | M+2 | 0.135 | 0.005 | 0.179 | 0.004 | 0.177 | 0.009 | 0.206 | 0.009 | 0.188 | 0.009 | 0.206 | 0.006 | 0.178 | 0.006 |
| | M+3 | 0.087 | 0.006 | 0.103 | 0.005 | 0.082 | 0.006 | 0.144 | 0.004 | 0.107 | 0.004 | 0.126 | 0.009 | 0.106 | 0.004 |
| | M+4 | 0.120 | 0.005 | 0.159 | 0.002 | 0.144 | 0.006 | 0.182 | 0.005 | 0.167 | 0.002 | 0.171 | 0.008 | 0.164 | 0.006 |
| | M+5 | 0.004 | 0.001 | 0.008 | 0.004 | 0.011 | 0.004 | 0.010 | 0.004 | 0.012 | 0.000 | 0.009 | 0.002 | 0.008 | 0.004 |
| | M+6 | 0.000 | 0.000 | 0.000 | 0.000 | 0.000 | 0.000 | 0.000 | 0.000 | 0.000 | 0.000 | 0.000 | 0.000 | 0.000 | 0.000 |
| | M+7 | 0.000 | 0.000 | 0.000 | 0.000 | 0.000 | 0.000 | 0.000 | 0.000 | 0.000 | 0.000 | 0.000 | 0.000 | 0.000 | 0.000 |
| | M+8 | 0.000 | 0.000 | 0.000 | 0.000 | 0.000 | 0.000 | 0.000 | 0.000 | 0.000 | 0.000 | 0.000 | 0.000 | 0.000 | 0.000 |
| | M+9 | 0.000 | 0.000 | 0.000 | 0.000 | 0.000 | 0.000 | 0.000 | 0.000 | 0.000 | 0.000 | 0.000 | 0.000 | 0.000 | 0.000 |
| | M+10 | 0.000 | 0.000 | 0.000 | 0.000 | 0.000 | 0.000 | 0.000 | 0.000 | 0.000 | 0.000 | 0.000 | 0.000 | 0.000 | 0.000 |
| CTP | M+0 | 0.710 | 0.084 | 0.686 | 0.031 | 0.547 | 0.020 | 0.308 | 0.023 | 0.357 | 0.024 | 0.387 | 0.045 | 0.503 | 0.063 |
| | M+1 | 0.126 | 0.063 | 0.145 | 0.017 | 0.190 | 0.016 | 0.205 | 0.027 | 0.207 | 0.014 | 0.192 | 0.030 | 0.179 | 0.023 |
| | M+2 | 0.090 | 0.009 | 0.095 | 0.028 | 0.115 | 0.025 | 0.166 | 0.013 | 0.154 | 0.015 | 0.176 | 0.001 | 0.145 | 0.032 |
| | M+3 | 0.045 | 0.006 | 0.039 | 0.020 | 0.071 | 0.020 | 0.177 | 0.035 | 0.120 | 0.010 | 0.104 | 0.012 | 0.086 | 0.010 |
| | M+4 | 0.029 | 0.029 | 0.035 | 0.018 | 0.061 | 0.011 | 0.132 | 0.015 | 0.141 | 0.009 | 0.118 | 0.019 | 0.079 | 0.012 |
| | M+5 | 0.000 | 0.000 | 0.000 | 0.000 | 0.016 | 0.016 | 0.011 | 0.011 | 0.022 | 0.011 | 0.023 | 0.013 | 0.008 | 0.008 |
| | M+6 | 0.000 | 0.000 | 0.000 | 0.000 | 0.000 | 0.000 | 0.000 | 0.000 | 0.000 | 0.000 | 0.000 | 0.000 | 0.000 | 0.000 |
| | M+7 | 0.000 | 0.000 | 0.000 | 0.000 | 0.000 | 0.000 | 0.000 | 0.000 | 0.000 | 0.000 | 0.000 | 0.000 | 0.000 | 0.000 |
| | M+8 | 0.000 | 0.000 | 0.000 | 0.000 | 0.000 | 0.000 | 0.000 | 0.000 | 0.000 | 0.000 | 0.000 | 0.000 | 0.000 | 0.000 |
| | M+9 | 0.000 | 0.000 | 0.000 | 0.000 | 0.000 | 0.000 | 0.000 | 0.000 | 0.000 | 0.000 | 0.000 | 0.000 | 0.000 | 0.000 |
| GTP | M+0 | 0.487 | 0.027 | 0.462 | 0.045 | 0.387 | 0.027 | 0.250 | 0.026 | 0.257 | 0.017 | 0.237 | 0.022 | 0.339 | 0.034 |
| | M+1 | 0.231 | 0.035 | 0.176 | 0.016 | 0.217 | 0.020 | 0.284 | 0.021 | 0.273 | 0.006 | 0.234 | 0.012 | 0.266 | 0.012 |
| | M+2 | 0.152 | 0.016 | 0.184 | 0.051 | 0.162 | 0.019 | 0.205 | 0.008 | 0.197 | 0.006 | 0.244 | 0.027 | 0.151 | 0.033 |
| | M+3 | 0.031 | 0.017 | 0.043 | 0.006 | 0.066 | 0.016 | 0.149 | 0.027 | 0.115 | 0.018 | 0.132 | 0.006 | 0.136 | 0.048 |
| | M+4 | 0.092 | 0.019 | 0.135 | 0.033 | 0.158 | 0.018 | 0.112 | 0.056 | 0.157 | 0.006 | 0.141 | 0.009 | 0.108 | 0.003 |
| | M+5 | 0.006 | 0.006 | 0.000 | 0.000 | 0.010 | 0.010 | 0.000 | 0.000 | 0.000 | 0.000 | 0.012 | 0.012 | 0.000 | 0.000 |
| | M+6 | 0.000 | 0.000 | 0.000 | 0.000 | 0.000 | 0.000 | 0.000 | 0.000 | 0.000 | 0.000 | 0.000 | 0.000 | 0.000 | 0.000 |
| | M+7 | 0.000 | 0.000 | 0.000 | 0.000 | 0.000 | 0.000 | 0.000 | 0.000 | 0.000 | 0.000 | 0.000 | 0.000 | 0.000 | 0.000 |
| | M+8 | 0.000 | 0.000 | 0.000 | 0.000 | 0.000 | 0.000 | 0.000 | 0.000 | 0.000 | 0.000 | 0.000 | 0.000 | 0.000 | 0.000 |
| | M+9 | 0.000 | 0.000 | 0.000 | 0.000 | 0.000 | 0.000 | 0.000 | 0.000 | 0.000 | 0.000 | 0.000 | 0.000 | 0.000 | 0.000 |
| | M+10 | 0.000 | 0.000 | 0.000 | 0.000 | 0.000 | 0.000 | 0.000 | 0.000 | 0.000 | 0.000 | 0.000 | 0.000 | 0.000 | 0.000 |
| GDP | M+0 | 0.343 | 0.016 | 0.337 | 0.034 | 0.347 | 0.019 | 0.194 | 0.002 | 0.261 | 0.013 | 0.205 | 0.019 | 0.275 | 0.015 |
| | M+1 | 0.186 | 0.009 | 0.150 | 0.020 | 0.168 | 0.015 | 0.176 | 0.020 | 0.214 | 0.017 | 0.222 | 0.011 | 0.226 | 0.057 |
| | M+2 | 0.119 | 0.020 | 0.120 | 0.007 | 0.127 | 0.016 | 0.149 | 0.016 | 0.170 | 0.008 | 0.180 | 0.012 | 0.116 | 0.003 |
| | M+3 | 0.222 | 0.021 | 0.238 | 0.028 | 0.239 | 0.030 | 0.270 | 0.006 | 0.200 | 0.006 | 0.223 | 0.005 | 0.243 | 0.017 |
| | M+4 | 0.103 | 0.022 | 0.125 | 0.006 | 0.120 | 0.017 | 0.165 | 0.013 | 0.150 | 0.025 | 0.135 | 0.014 | 0.131 | 0.027 |
| | M+5 | 0.000 | 0.000 | 0.000 | 0.000 | 0.000 | 0.000 | 0.006 | 0.006 | 0.006 | 0.006 | 0.005 | 0.005 | 0.000 | 0.000 |
| | M+6 | 0.000 | 0.000 | 0.000 | 0.000 | 0.000 | 0.000 | 0.000 | 0.000 | 0.000 | 0.000 | 0.000 | 0.000 | 0.000 | 0.000 |
| | M+7 | 0.000 | 0.000 | 0.000 | 0.000 | 0.000 | 0.000 | 0.000 | 0.000 | 0.000 | 0.000 | 0.000 | 0.000 | 0.000 | 0.000 |
| | M+8 | 0.000 | 0.000 | 0.000 | 0.000 | 0.000 | 0.000 | 0.000 | 0.000 | 0.000 | 0.000 | 0.000 | 0.000 | 0.000 | 0.000 |
| | M+9 | 0.000 | 0.000 | 0.000 | 0.000 | 0.000 | 0.000 | 0.000 | 0.000 | 0.000 | 0.000 | 0.000 | 0.000 | 0.000 | 0.000 |
| | M+10 | 0.000 | 0.000 | 0.000 | 0.000 | 0.000 | 0.000 | 0.000 | 0.000 | 0.000 | 0.000 | 0.000 | 0.000 | 0.000 | 0.000 |
| UTP | M+0 | 0.313 | 0.016 | 0.351 | 0.040 | 0.274 | 0.013 | 0.160 | 0.024 | 0.178 | 0.003 | 0.167 | 0.004 | 0.212 | 0.007 |
| | M+1 | 0.197 | 0.003 | 0.201 | 0.020 | 0.194 | 0.010 | 0.245 | 0.005 | 0.229 | 0.004 | 0.249 | 0.008 | 0.224 | 0.014 |
| | M+2 | 0.177 | 0.010 | 0.157 | 0.010 | 0.190 | 0.020 | 0.196 | 0.011 | 0.210 | 0.002 | 0.191 | 0.005 | 0.186 | 0.011 |
| | M+3 | 0.129 | 0.013 | 0.118 | 0.008 | 0.151 | 0.004 | 0.178 | 0.010 | 0.158 | 0.007 | 0.179 | 0.000 | 0.157 | 0.009 |
| | M+4 | 0.145 | 0.012 | 0.139 | 0.005 | 0.156 | 0.014 | 0.186 | 0.004 | 0.173 | 0.001 | 0.174 | 0.006 | 0.186 | 0.006 |
| | M+5 | 0.039 | 0.006 | 0.034 | 0.003 | 0.036 | 0.007 | 0.037 | 0.003 | 0.052 | 0.001 | 0.039 | 0.006 | 0.035 | 0.010 |
| | M+6 | 0.000 | 0.000 | 0.000 | 0.000 | 0.000 | 0.000 | 0.000 | 0.000 | 0.000 | 0.000 | 0.000 | 0.000 | 0.000 | 0.000 |
| | M+7 | 0.000 | 0.000 | 0.000 | 0.000 | 0.000 | 0.000 | 0.000 | 0.000 | 0.000 | 0.000 | 0.000 | 0.000 | 0.000 | 0.000 |
| | M+8 | 0.000 | 0.000 | 0.000 | 0.000 | 0.000 | 0.000 | 0.000 | 0.000 | 0.000 | 0.000 | 0.000 | 0.000 | 0.000 | 0.000 |
| | M+9 | 0.000 | 0.000 | 0.000 | 0.000 | 0.000 | 0.000 | 0.000 | 0.000 | 0.000 | 0.000 | 0.000 | 0.000 | 0.000 | 0.000 |
| UDP | M+0 | 0.417 | 0.068 | 0.486 | 0.045 | 0.380 | 0.025 | 0.184 | 0.025 | 0.244 | 0.011 | 0.206 | 0.019 | 0.245 | 0.012 |
| | M+1 | 0.122 | 0.010 | 0.141 | 0.015 | 0.146 | 0.024 | 0.254 | 0.020 | 0.229 | 0.009 | 0.227 | 0.010 | 0.206 | 0.037 |
| | M+2 | 0.138 | 0.015 | 0.116 | 0.021 | 0.169 | 0.028 | 0.189 | 0.014 | 0.158 | 0.009 | 0.175 | 0.020 | 0.181 | 0.010 |
| | M+3 | 0.120 | 0.039 | 0.101 | 0.020 | 0.077 | 0.019 | 0.139 | 0.008 | 0.145 | 0.008 | 0.147 | 0.012 | 0.134 | 0.012 |
| | M+4 | 0.160 | 0.009 | 0.144 | 0.028 | 0.211 | 0.012 | 0.197 | 0.020 | 0.183 | 0.005 | 0.194 | 0.019 | 0.196 | 0.030 |
| | M+5 | 0.000 | 0.000 | 0.010 | 0.010 | 0.007 | 0.007 | 0.017 | 0.015 | 0.032 | 0.011 | 0.018 | 0.006 | 0.031 | 0.011 |
| | M+6 | 0.000 | 0.000 | 0.000 | 0.000 | 0.000 | 0.000 | 0.000 | 0.000 | 0.000 | 0.000 | 0.000 | 0.000 | 0.000 | 0.000 |
| | M+7 | 0.000 | 0.000 | 0.000 | 0.000 | 0.000 | 0.000 | 0.000 | 0.000 | 0.000 | 0.000 | 0.000 | 0.000 | 0.000 | 0.000 |
| | M+8 | 0.000 | 0.000 | 0.000 | 0.000 | 0.000 | 0.000 | 0.000 | 0.000 | 0.000 | 0.000 | 0.000 | 0.000 | 0.000 | 0.000 |
| | M+9 | 0.000 | 0.000 | 0.000 | 0.000 | 0.000 | 0.000 | 0.000 | 0.000 | 0.000 | 0.000 | 0.000 | 0.000 | 0.000 | 0.000 |
| UMP | M+0 | 0.467 | 0.017 | 0.432 | 0.017 | 0.424 | 0.010 | 0.260 | 0.020 | 0.268 | 0.012 | 0.313 | 0.010 | 0.354 | 0.016 |
| | M+1 | 0.153 | 0.003 | 0.154 | 0.010 | 0.162 | 0.004 | 0.249 | 0.010 | 0.234 | 0.003 | 0.200 | 0.005 | 0.182 | 0.006 |
| | M+2 | 0.129 | 0.019 | 0.147 | 0.011 | 0.138 | 0.016 | 0.171 | 0.010 | 0.173 | 0.005 | 0.171 | 0.004 | 0.174 | 0.006 |
| | M+3 | 0.098 | 0.008 | 0.121 | 0.011 | 0.104 | 0.003 | 0.141 | 0.003 | 0.142 | 0.005 | 0.131 | 0.005 | 0.113 | 0.007 |
| | M+4 | 0.117 | 0.019 | 0.125 | 0.011 | 0.140 | 0.010 | 0.159 | 0.011 | 0.153 | 0.004 | 0.148 | 0.003 | 0.144 | 0.005 |
| | M+5 | 0.029 | 0.005 | 0.012 | 0.008 | 0.024 | 0.003 | 0.018 | 0.010 | 0.027 | 0.005 | 0.031 | 0.005 | 0.027 | 0.005 |
| | M+6 | 0.000 | 0.000 | 0.000 | 0.000 | 0.000 | 0.000 | 0.000 | 0.000 | 0.000 | 0.000 | 0.000 | 0.000 | 0.000 | 0.000 |
| | M+7 | 0.000 | 0.000 | 0.000 | 0.000 | 0.000 | 0.000 | 0.000 | 0.000 | 0.000 | 0.000 | 0.000 | 0.000 | 0.000 | 0.000 |
| | M+8 | 0.000 | 0.000 | 0.000 | 0.000 | 0.000 | 0.000 | 0.000 | 0.000 | 0.000 | 0.000 | 0.000 | 0.000 | 0.000 | 0.000 |
| | M+9 | 0.000 | 0.000 | 0.000 | 0.000 | 0.000 | 0.000 | 0.000 | 0.000 | 0.000 | 0.000 | 0.000 | 0.000 | 0.000 | 0.000 |
| GlcNAc-1/6P | M+0 | 0.640 | 0.089 | 0.493 | 0.081 | 0.684 | 0.082 | 0.328 | 0.024 | 0.362 | 0.047 | 0.326 | 0.014 | 0.489 | 0.026 |
| | M+1 | 0.000 | 0.000 | 0.000 | 0.000 | 0.000 | 0.000 | 0.000 | 0.000 | 0.000 | 0.000 | 0.000 | 0.000 | 0.000 | 0.000 |
| | M+2 | 0.307 | 0.039 | 0.467 | 0.099 | 0.316 | 0.082 | 0.568 | 0.032 | 0.522 | 0.054 | 0.530 | 0.004 | 0.402 | 0.034 |
| | M+3 | 0.000 | 0.000 | 0.000 | 0.000 | 0.000 | 0.000 | 0.000 | 0.000 | 0.000 | 0.000 | 0.000 | 0.000 | 0.000 | 0.000 |
| | M+4 | 0.053 | 0.053 | 0.040 | 0.040 | 0.000 | 0.000 | 0.104 | 0.012 | 0.116 | 0.011 | 0.144 | 0.018 | 0.110 | 0.032 |
| | M+5 | 0.000 | 0.000 | 0.000 | 0.000 | 0.000 | 0.000 | 0.000 | 0.000 | 0.000 | 0.000 | 0.000 | 0.000 | 0.000 | 0.000 |
| | M+6 | 0.000 | 0.000 | 0.000 | 0.000 | 0.000 | 0.000 | 0.000 | 0.000 | 0.000 | 0.000 | 0.000 | 0.000 | 0.000 | 0.000 |
| | M+7 | 0.000 | 0.000 | 0.000 | 0.000 | 0.000 | 0.000 | 0.000 | 0.000 | 0.000 | 0.000 | 0.000 | 0.000 | 0.000 | 0.000 |
| | M+8 | 0.000 | 0.000 | 0.000 | 0.000 | 0.000 | 0.000 | 0.000 | 0.000 | 0.000 | 0.000 | 0.000 | 0.000 | 0.000 | 0.000 |
| UDP-GlcNAc | M+0 | 0.075 | 0.001 | 0.046 | 0.002 | 0.061 | 0.002 | 0.015 | 0.001 | 0.028 | 0.001 | 0.022 | 0.000 | 0.011 | 0.000 |
| | M+1 | 0.041 | 0.001 | 0.030 | 0.001 | 0.037 | 0.001 | 0.013 | 0.001 | 0.023 | 0.001 | 0.018 | 0.000 | 0.062 | 0.003 |
| | M+2 | 0.150 | 0.001 | 0.148 | 0.003 | 0.159 | 0.003 | 0.088 | 0.002 | 0.107 | 0.003 | 0.103 | 0.000 | 0.044 | 0.002 |
| | M+3 | 0.120 | 0.002 | 0.134 | 0.002 | 0.128 | 0.001 | 0.161 | 0.002 | 0.141 | 0.001 | 0.144 | 0.004 | 0.206 | 0.011 |
| | M+4 | 0.181 | 0.004 | 0.193 | 0.002 | 0.192 | 0.002 | 0.177 | 0.001 | 0.178 | 0.001 | 0.182 | 0.004 | 0.220 | 0.013 |
| | M+5 | 0.132 | 0.002 | 0.143 | 0.000 | 0.132 | 0.002 | 0.185 | 0.001 | 0.164 | 0.002 | 0.171 | 0.002 | 0.295 | 0.017 |
| | M+6 | 0.134 | 0.001 | 0.156 | 0.001 | 0.139 | 0.001 | 0.174 | 0.003 | 0.170 | 0.001 | 0.169 | 0.004 | 0.250 | 0.017 |
| | M+7 | 0.074 | 0.002 | 0.071 | 0.002 | 0.066 | 0.002 | 0.086 | 0.001 | 0.085 | 0.002 | 0.088 | 0.001 | 0.252 | 0.010 |
| | M+8 | 0.058 | 0.001 | 0.056 | 0.003 | 0.057 | 0.002 | 0.069 | 0.001 | 0.069 | 0.001 | 0.070 | 0.001 | 0.125 | 0.007 |
| | M+9 | 0.022 | 0.000 | 0.017 | 0.001 | 0.020 | 0.001 | 0.020 | 0.001 | 0.023 | 0.000 | 0.022 | 0.001 | 0.096 | 0.005 |
| | M+10 | 0.012 | 0.001 | 0.009 | 0.001 | 0.010 | 0.000 | 0.011 | 0.000 | 0.012 | 0.000 | 0.011 | 0.001 | 0.034 | 0.003 |
| | M+11 | 0.000 | 0.000 | 0.000 | 0.000 | 0.000 | 0.000 | 0.000 | 0.000 | 0.000 | 0.000 | 0.000 | 0.000 | 0.000 | 0.000 |
| | M+12 | 0.000 | 0.000 | 0.000 | 0.000 | 0.000 | 0.000 | 0.000 | 0.000 | 0.000 | 0.000 | 0.000 | 0.000 | 0.000 | 0.000 |
| | M+13 | 0.000 | 0.000 | 0.000 | 0.000 | 0.000 | 0.000 | 0.000 | 0.000 | 0.000 | 0.000 | 0.000 | 0.000 | 0.000 | 0.000 |
| | M+14 | 0.000 | 0.000 | 0.000 | 0.000 | 0.000 | 0.000 | 0.000 | 0.000 | 0.000 | 0.000 | 0.000 | 0.000 | 0.000 | 0.000 |
| | M+15 | 0.000 | 0.000 | 0.000 | 0.000 | 0.000 | 0.000 | 0.000 | 0.000 | 0.000 | 0.000 | 0.000 | 0.000 | 0.000 | 0.000 |
| | M+16 | 0.000 | 0.000 | 0.000 | 0.000 | 0.000 | 0.000 | 0.000 | 0.000 | 0.000 | 0.000 | 0.000 | 0.000 | 0.000 | 0.000 |
| | M+17 | 0.000 | 0.000 | 0.000 | 0.000 | 0.000 | 0.000 | 0.000 | 0.000 | 0.000 | 0.000 | 0.000 | 0.000 | 0.000 | 0.000 |

Values represent the mean and standard error (n=3).

**Supplementary Table 13.** Nucleobase Secretion by CAR-T Cells

| ***Donor 4- CAR-T cell $^{15}$N Ion Counts*** | | | | | | | | | | | | | | | | | |
|---|---|---|---|---|---|---|---|---|---|---|---|---|---|---|---|---|---|
| **Nucleobase** | **Day** | **EGFRt** | **s.e.m.** | **CD19** | **s.e.m.** | **Leu16** | **s.e.m.** | **Rituximab** | **s.e.m.** | **RFR-LCDR** | **s.e.m.** | **Rituximab.AA** | **s.e.m.** | **RFR-LCDR.AA** | **s.e.m.** | **GD2** | **s.e.m.** |
| Uracil | 1 | 948.60 | 490.11 | 893.85 | 296.05 | 346.07 | 346.07 | 1563.25 | 81.32 | 1790.83 | 302.60 | 1950.46 | 251.50 | 1917.49 | 19.12 | 1165.45 | 583.74 |
| | 2 | 947.33 | 188.16 | 1387.51 | 66.85 | 716.42 | 358.63 | 2423.85 | 480.24 | 2199.59 | 176.99 | 2646.81 | 333.64 | 2556.56 | 69.96 | 2508.30 | 108.51 |
| | 3 | 1489.71 | 271.74 | 1493.54 | 132.64 | 1852.78 | 375.00 | 3649.70 | 386.69 | 3202.40 | 134.41 | 2612.60 | 277.56 | 2714.32 | 398.27 | 3375.98 | 168.09 |
| Thymine | 1 | 775.62 | 147.81 | 1389.45 | 192.21 | 1359.49 | 92.70 | 1401.03 | 540.52 | 1630.56 | 254.67 | 1224.41 | 404.22 | 1369.82 | 140.11 | 1863.53 | 237.77 |
| | 2 | 750.76 | 126.75 | 1642.89 | 379.54 | 1053.88 | 30.48 | 2961.76 | 310.95 | 2443.25 | 157.68 | 2946.84 | 625.24 | 1916.14 | 187.33 | 3556.00 | 663.77 |
| | 3 | 613.39 | 140.23 | 1402.02 | 506.01 | 316.67 | 180.00 | 4686.03 | 408.47 | 2228.05 | 67.05 | 1537.90 | 631.81 | 1112.32 | 31.91 | 4073.44 | 902.23 |
| Xanthine | 1 | 4255.28 | 1083.82 | 6979.07 | 1559.44 | 7436.83 | 982.35 | 10628.17 | 4431.74 | 15308.39 | 957.20 | 7320.93 | 425.95 | 8123.80 | 1083.43 | 6515.34 | 1065.75 |
| | 2 | 8844.29 | 978.89 | 14641.97 | 1478.87 | 13135.27 | 1335.12 | 15221.08 | 199.94 | 12113.26 | 1985.64 | 16605.38 | 344.78 | 11503.16 | 1600.20 | 15106.84 | 483.69 |
| | 3 | 20211.96 | 1682.55 | 30902.21 | 1014.27 | 26162.02 | 876.10 | 40891.40 | 2824.78 | 28143.47 | 1178.94 | 30732.81 | 1253.36 | 23864.83 | 1848.95 | 38395.48 | 2487.68 |

Values represent the mean and standard error (n=3).

**Supplementary Table 14.** Summary of Experimental Conditions

| ***Experimental Conditions*** | | | |
|---|---|---|---|
| **Isotope Tracers** | **Donor** | **Media Condition** | **CAR-T Cells** |
| [U-$^{13}C_6$] glucose, 50% [$^{15}N_2$]glutamine | 1 | RPMI +10% dFBS | EGFRt, CD19, Leu16, Rituximab, RFR-LCDR |
| 50% [U-$^{13}C_5$$^{15}N_2$]glutamine | 3 | RPMI +10% dFBS | EGFRt, CD19, Leu16, Rituximab, RFR-LCDR, GD2 |
| [U-$^{13}C_6$] glucose, 50% [γ-$^{15}$N]glutamine | 4 | RPMI +10% dFBS | EGFRt, CD19, Leu16, Rituximab, RFR-LCDR, Rituximab.AA, RFR-LCDR.AA, GD2 |
| [1,2-$^{13}C_2$]glucose | 5 | RPMI +10% dFBS | EGFRt, CD19, Leu16, Rituximab, RFR-LCDR, Rituximab.AA, RFR-LCDR.AA |
| [1,2-$^{13}C_2$]glucose | 6 | RPMI +10% dFBS | EGFRt, CD19, Rituximab |
| [U-$^{13}C_6$] glucose, 50% [γ-$^{15}$N]glutamine | 4 | RPMI +10% dFBS + 300 $\mu$M alanine | EGFRt, CD19, Leu16, Rituximab, RFR-LCDR, Rituximab.AA, RFR-LCDR.AA, GD2 |
| [U-$^{13}C_6$] glucose, 50% [$^{15}N_2$]glutamine | 1 | RPMI +10% dFBS + 800 $\mu$M ammonia | EGFRt, CD19, Leu16, Rituximab, RFR-LCDR |
| 50% [U-$^{13}C_5$$^{15}N_2$]glutamine | 3 | RPMI +10% dFBS + 800 $\mu$M ammonia | EGFRt, CD19, Leu16, Rituximab, RFR-LCDR, GD2 |
| [U-$^{13}C_6$] glucose, $^{15}NH_4Cl$ | 4 | RPMI +10% dFBS + 800 $\mu$M ammonia | EGFRt, CD19, Leu16, Rituximab, RFR-LCDR, Rituximab.AA, RFR-LCDR.AA, GD2 |
| [U-$^{13}C_6$] glucose, 50% [$^{15}N_2$]glutamine | 1 | RPMI +10% dFBS + 30 mM lactate | EGFRt, CD19, Leu16, Rituximab, RFR-LCDR |
| 50% [U-$^{13}C_5$$^{15}N_2$]glutamine | 3 | RPMI +10% dFBS + 30 mM lactate | EGFRt, CD19, Leu16, Rituximab, RFR-LCDR, GD2 |
| [U-$^{13}C_6$] glucose, 50% [$^{15}N_2$]glutamine | 1 | RPMI +10% dFBS + 800 $\mu$M ammonia + 30 mM lactate | EGFRt, CD19, Leu16, Rituximab, RFR-LCDR |
| 50% [U-$^{13}C_5$$^{15}N_2$]glutamine | 3 | RPMI +10% dFBS + 800 $\mu$M ammonia + 30 mM lactate | EGFRt, CD19, Leu16, Rituximab, RFR-LCDR, GD2 |
